# Bioinformatics Analysis of Key Genes and Pathways for Obesity Associated Type 2 Diabetes Mellitus as a Therapeutic Target

**DOI:** 10.1101/2020.12.25.424383

**Authors:** Basavaraj Vastrad, Anandkumar Tengli, Chanabasayya Vastrad, Iranna Kotturshetti

## Abstract

Obesity associated type 2 diabetes mellitus is one of the most common metabolic disorder worldwide. The prognosis of obesity associated type 2 diabetes mellitus patients has remained poor, though considerable efforts have been made to improve the treatment of this metabolic disorder. Therefore, identifying significant differentially expressed genes (DEGs) associated in metabolic disorder advancement and exploiting them as new biomarkers or potential therapeutic targets for metabolic disorder is highly valuable. Differentially expressed genes (DEGs) were screened out from gene expression omnibus (GEO) dataset (GSE132831) and subjected to GO and REACTOME pathway enrichment analyses. The protein - protein interactions network, module analysis, target gene - miRNA regulatory network and target gene - TF regulatory network were constructed, and the top ten hub genes were selected. The relative expression of hub genes was detected in RT-PCR. Furthermore, diagnostic value of hub genes in obesity associated type 2 diabetes mellitus patients was investigated using the receiver operating characteristic (ROC) analysis. Small molecules were predicted for obesity associated type 2 diabetes mellitus by using molecular docking studies. A total of 872 DEGs, including 439 up regulated genes and 432 down regulated genes were observed. Second, functional enrichment analysis showed that these DEGs are mainly involved in the axon guidance, neutrophil degranulation, plasma membrane bounded cell projection organization and cell activation. The top ten hub genes (MYH9, FLNA, DCTN1, CLTC, ERBB2, TCF4, VIM, LRRK2, IFI16 and CAV1) could be utilized as potential diagnostic indicators for obesity associated type 2 diabetes mellitus. The hub genes were validated in obesity associated type 2 diabetes mellitus. This investigation found effective and reliable molecular biomarkers for diagnosis and prognosis by integrated bioinformatics analysis, suggesting new and key therapeutic targets for obesity associated type 2 diabetes mellitus.

## Introduction

Obesity and type 2 diabetes mellitus are a major metabolic or endocrine disorder and are dramatically increasing throughout the globe [1]. The prevalence of obesity associated type 2 diabetes mellitus is 86% [2]. Obesity associated type 2 diabetes mellitus might be responsible for causing cardiovascular diseases [3], hypertension [4], and neurological and neuropsychiatric disorders [5] and asthma [6]. Till today, there is no cure for obesity associated type 2 diabetes mellitus, and treatment and mediation tailored to clinical features are endorsed. Genetic and environmental factors are two initial contributors to this disorder [7]. Exploration of the molecular mechanisms of obesity associated type 2 diabetes mellitus will develop the considerate of its pathogenesis and has key implications for designing new therapy.

Molecular mechanisms of obesity associated type 2 diabetes mellitus have been increasingly studied. Previous investigations showed that obesity associated type 2 diabetes mellitus in human was related to genes and signaling pathways. Key genes such as ENPP1 [8] and FTO [9] were responsible for development of obesity associated type 2 diabetes mellitus. Recent investigations showed that PI3K/AKT pathway [10] and TLR pathway [11] as a potential target for obesity associated type 2 diabetes mellitus. However, certain key genes and pathways of obesity associated type 2 diabetes mellitus have not been completely investigated. Further studies are necessary to elucidate these essential genes and pathways to provide novel therapeutic targets for the treatment of obesity associated type 2 diabetes mellitus.

Therefore, in this investigation, we downloaded the expression profiling by high throughput sequencing data GSE132831, provided by Osinski et al. [12], from Gene Expression Omnibus (GEO, http://www.ncbi.nlm.nih.gov/geo/) [13] database to identify the differentially expressed genes (DEGs) between diabetic obese samples and non diabetic obese samples. With the identified DEGs, we performed Gene Ontology (GO) and pathway enrichment analyses to investigate the functions and pathways enriched by the DEGs. Additionally, we constructed a protein–protein interaction (PPI) network and modules screened out some important gene nodes to perform clustering analysis. Furthermore, we constructed target gene - miRNA regulatory network and target gene - TF regulatory network based on these key genes to investigate the potential relationships between genes and obesity associated type 2 diabetes mellitus. Finally, hub genes were validated by using receiver operating characteristic (ROC) curve analysis and RT-PCR.

## Materials and Methods

### RNA sequencing data

The expression profiling by high throughput sequencing data GSE132831 was downloaded from the GEO database, which was based on the platform of GPL1857 Illumina NextSeq 500 (Homo sapiens). This dataset, including 104 diabetic obese samples and 120 non diabetic obese samples, was deposited by Osinski et al. [12].

### Identification of DEGs

The limma R/Bioconductor software package was used to perform the identification of DEGs between diabetic obese samples and non diabetic obese samples in R software [14]. The cutoff criteria were |logFC| > 1.112 for up regulated genes, |logFC| < −0.64 for down regulated genes and a P-value <0.05.

### Gene Ontology (GO) and Reactome pathway enrichment analysis of DEGs

The ToppGene (ToppFun) (https://toppgene.cchmc.org/enrichment.jsp) [15] is a gene functional enrichment program, providing a large series of functional annotation tools for researchers to decipher the biological implications behind huge amounts of genes. To understand the deeper biological meaning of DEGs, GO (http://geneontology.org/) [16] and REACTOME (https://reactome.org/) [17] pathway enrichment analysis of identified DEGs was performed using ToppGene. GO functional analysis consists of three categories: Biological process (BP), cellular component (CC) and molecular function (MF). P<0.05 was set as the threshold value.

### Protein-protein interaction (PPI) network and module analysis

The IID interactome (http://iid.ophid.utoronto.ca/) [18] is an online database containing known and predicted PPI networks. In this investigation, a PPI network of identified DEGs in dataset was identified using the IID interactome database (combined score >0.4) and subsequently visualized using Cytoscape (http://www.cytoscape.org/) software (version 3.8.2) [19]. The regulatory relationship between genes were analyzed through topological property of computing network including the node degree [20], betweenness centrality [21], stress centrality [22] and closeness centrality [23] by using the Network Analyzer app within Cytoscape. The PEWCC1 (http://apps.cytoscape.org/apps/PEWCC1) [24] program within Cytoscape was used to detect modules of the PPI network. The cut-off criteria were set as follows: Degree, ≥2; node score, ≥ 0.2; k-score, 0.2; maximum depth, 100. The GO and pathway enrichment analysis of the identified modules was then performed using the ToppGene database.

### Target gene - miRNA regulatory network

In the current investigation, the miRNAs associated with DEGs were searched utilizing miRNet database (https://www.mirnet.ca/) [25] online tool, and target gene - miRNA regulatory network was visualized by Cytoscape software.

### Target gene - TF regulatory network

In the current investigation, the TFs associated with DEGs were searched utilizing NetworkAnalyst database (https://www.networkanalyst.ca/) [26] online tool, and target gene - TF regulatory network was visualized by Cytoscape software.

### Receiver operating characteristic (ROC) analysis

A ROC analysis is a technique for visualizing, construct and determining classifiers based on their achievement. A diagnostic test was firstly performed in order to measure the diagnostic value of candidate biomarkers in obesity associated type 2 diabetes mellitus. Sensitivity and specificity of each biomarker in this diagnostic test were determined. ROC curves were retrieved by plotting the sensitivity, against the specificity using the pROC in R software [27]. Area under the ROC curve (AUC) was determined to predict the efficiency of this diagnostic test. A test with AUC bigger than 0.9 gets great efficiency, 0.7-0.9, modest efficiency and 0.5-0.7, small efficiency.

### Validation of the expression levels of candidate genes by RT-PCR

The TRI Reagent® (Sigma, USA) was used to isolate total RNA from the Pre-adipocytes; Normal, Human (ATCC® PCS-210-010™) and obesity 3T3-L1 cells (ATCC® CL-173). Then, RNA was reversely transcribed by using Reverse transcription cDNA kit (Thermo Fisher Scientific, Waltham, MA, USA), according to the manufacturer’s instructions. The expression of the top hub genes was analyzed on the 7 Flex real-time PCR system (Thermo Fisher Scientific, Waltham, MA, USA) connect real-time detection system. The expression levels of hub genes were normalized to The expression levels of 10 DEGs were normalized to beta-actin. Relative gene expression levels were calculated using the 2^-ΔΔCT^ (CT, cycle threshold) method [28]. All primers are listed in Table 1.

**Table 1.**
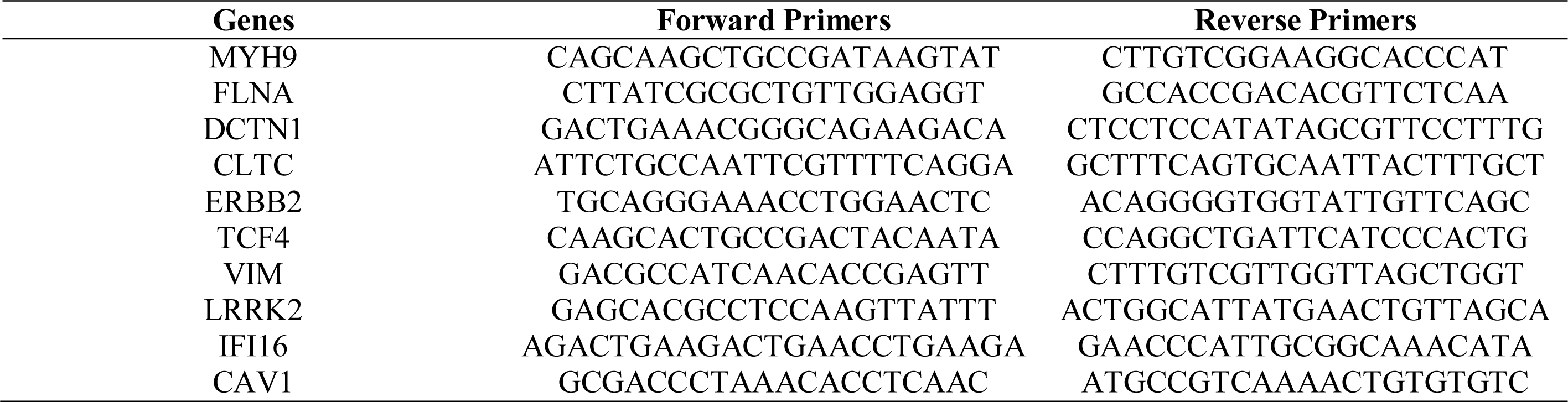
The sequences of primers for quantitative RT-PCR.

**Table 2.**
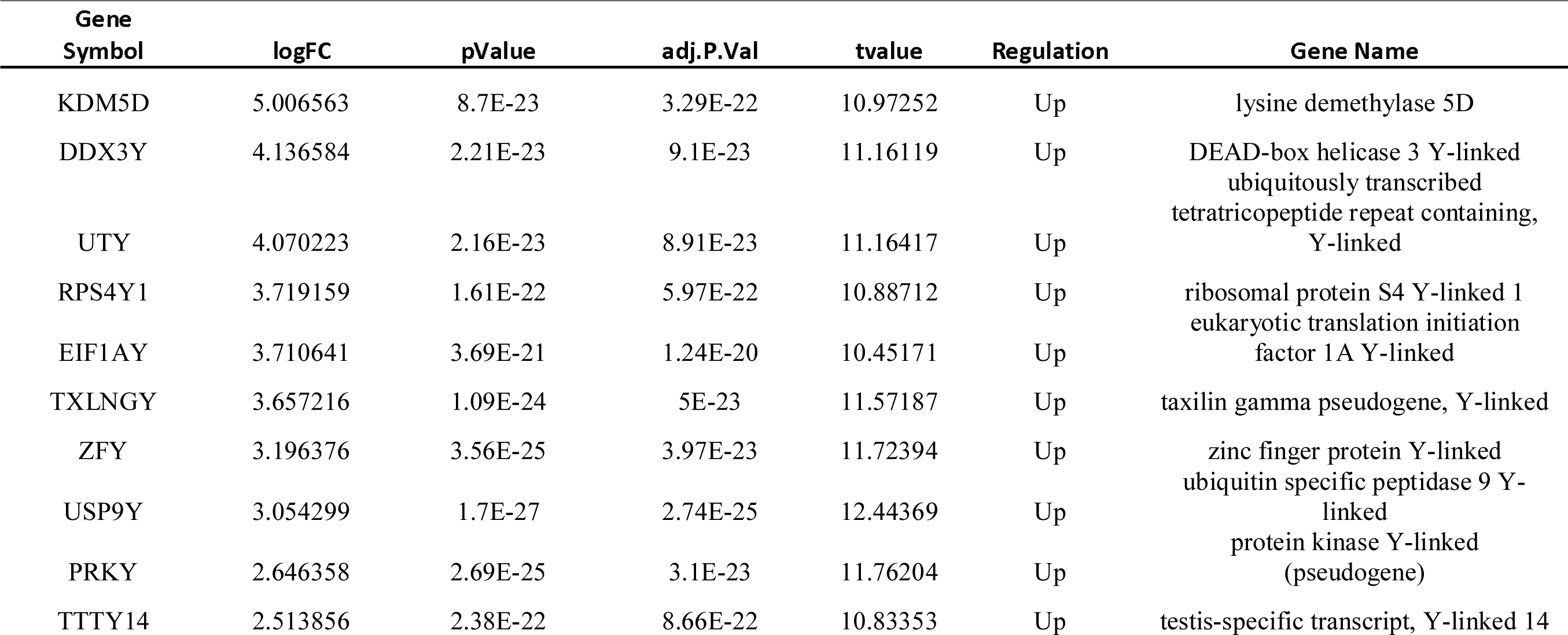
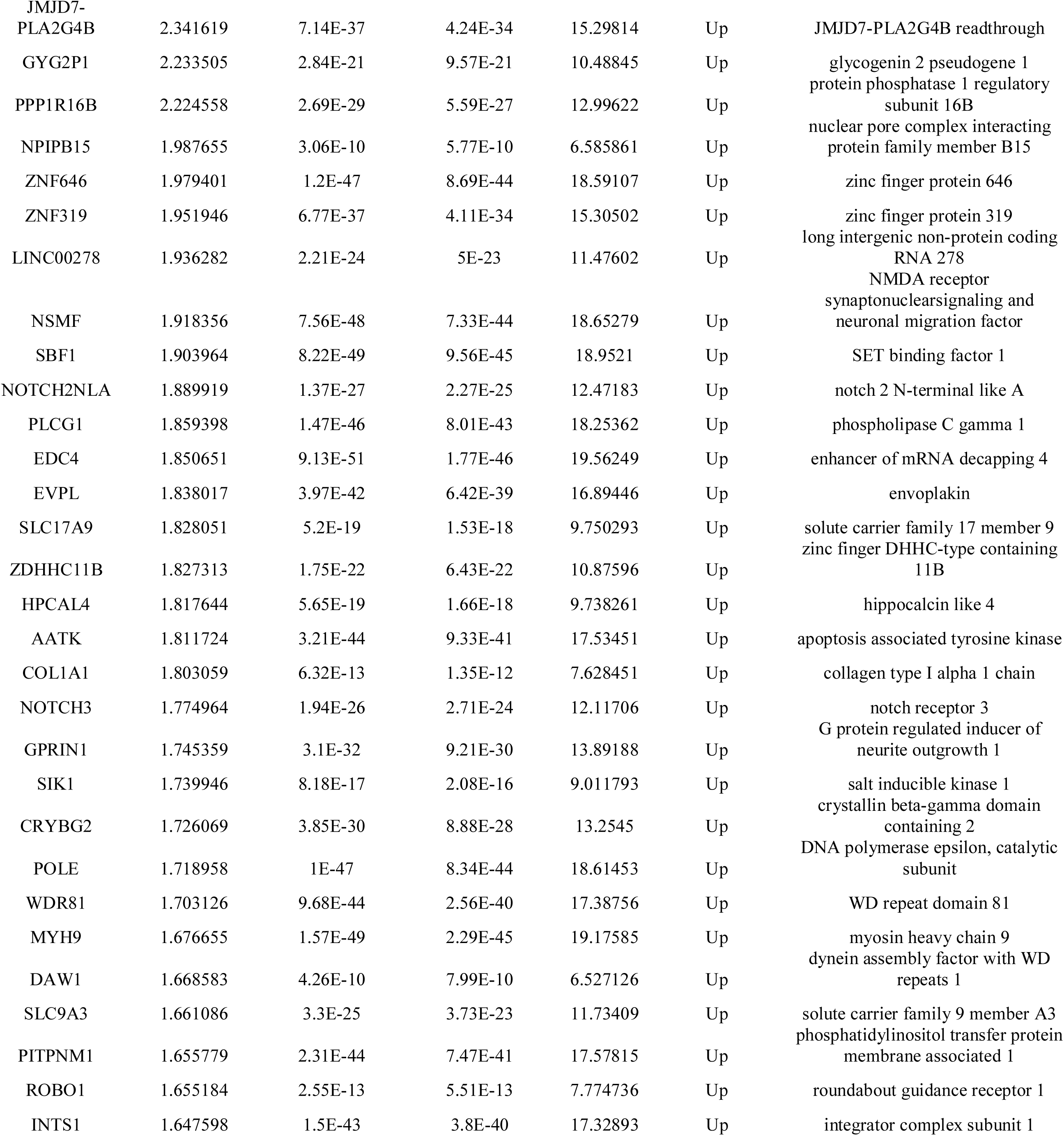
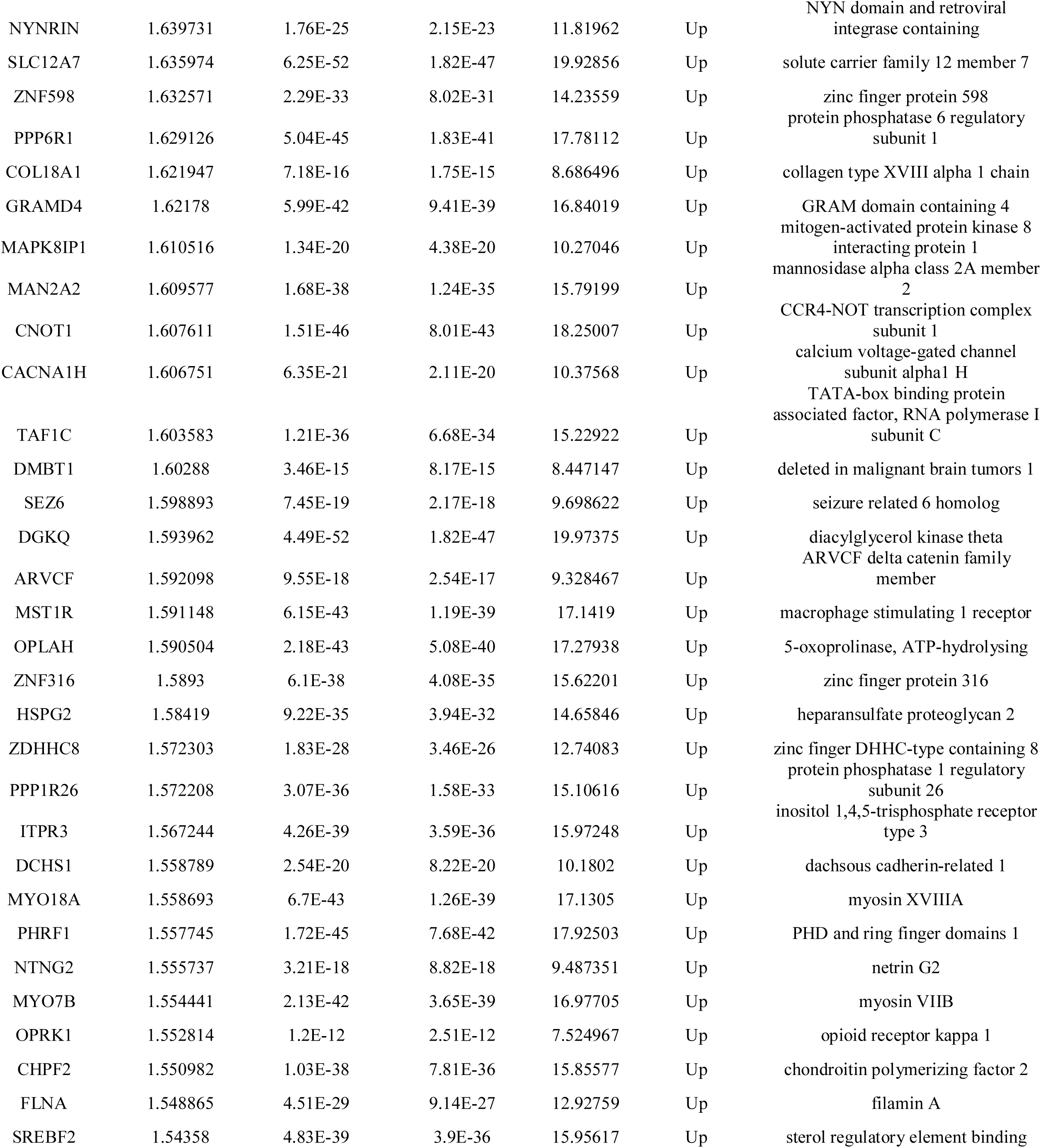
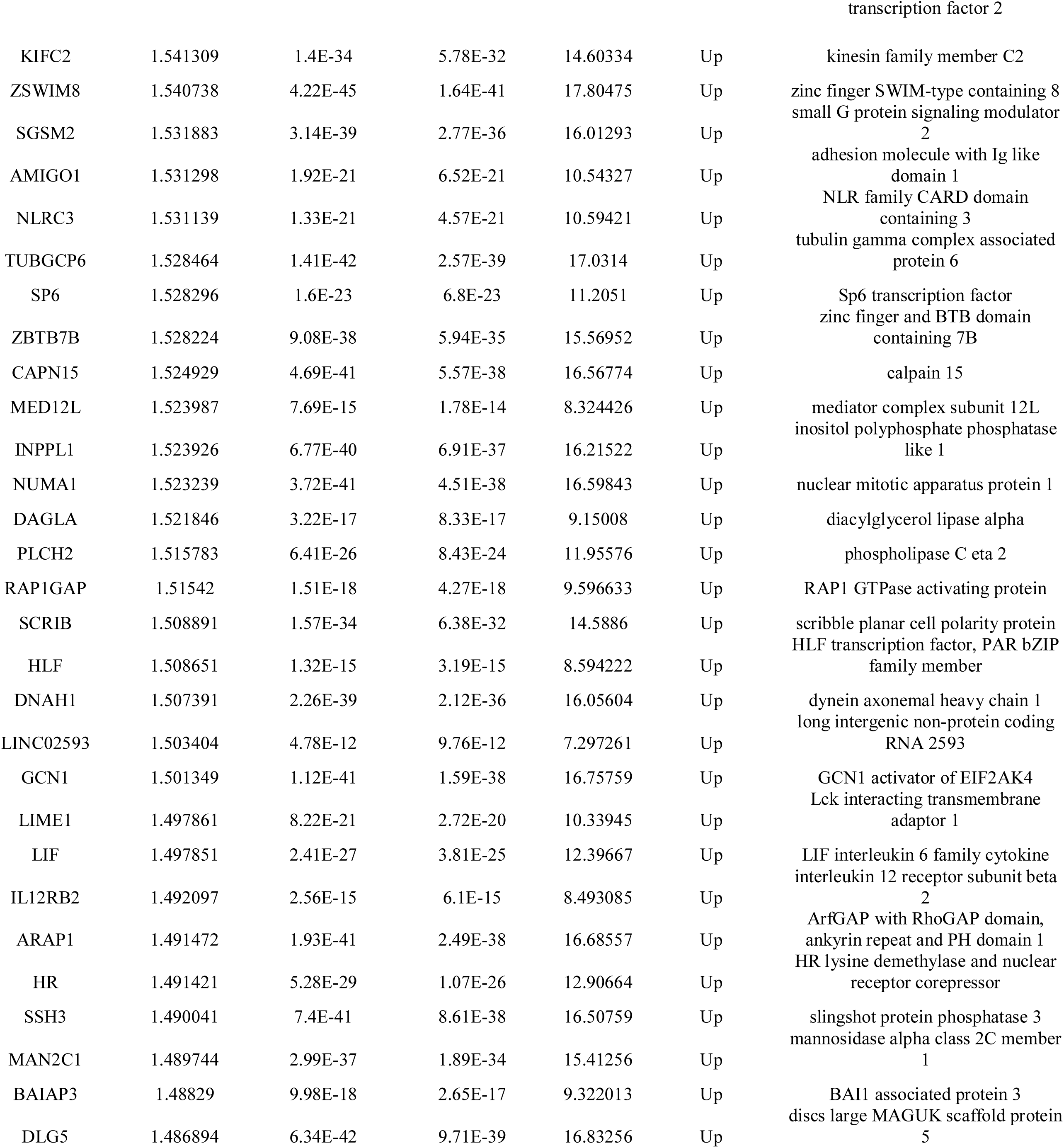
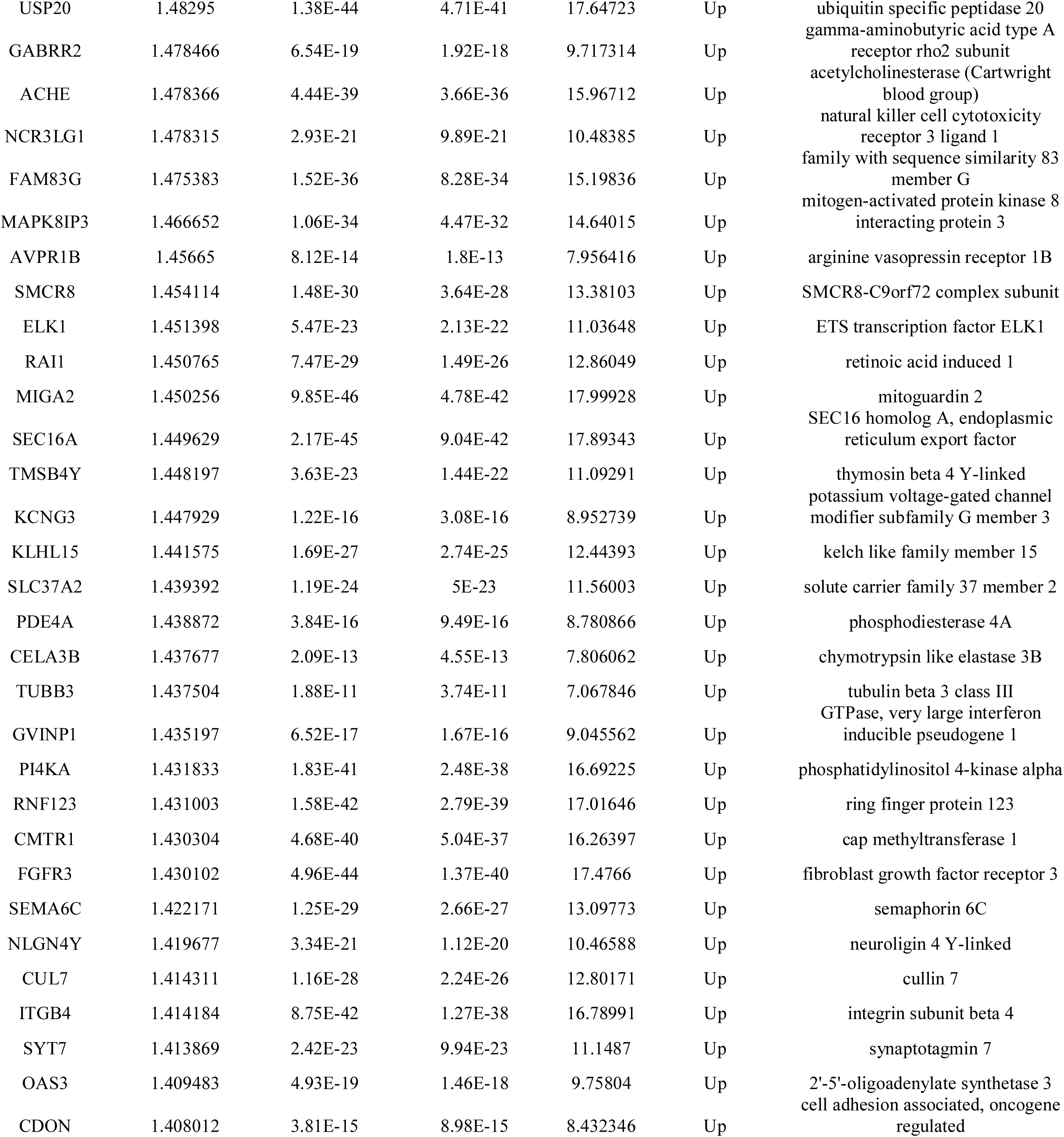
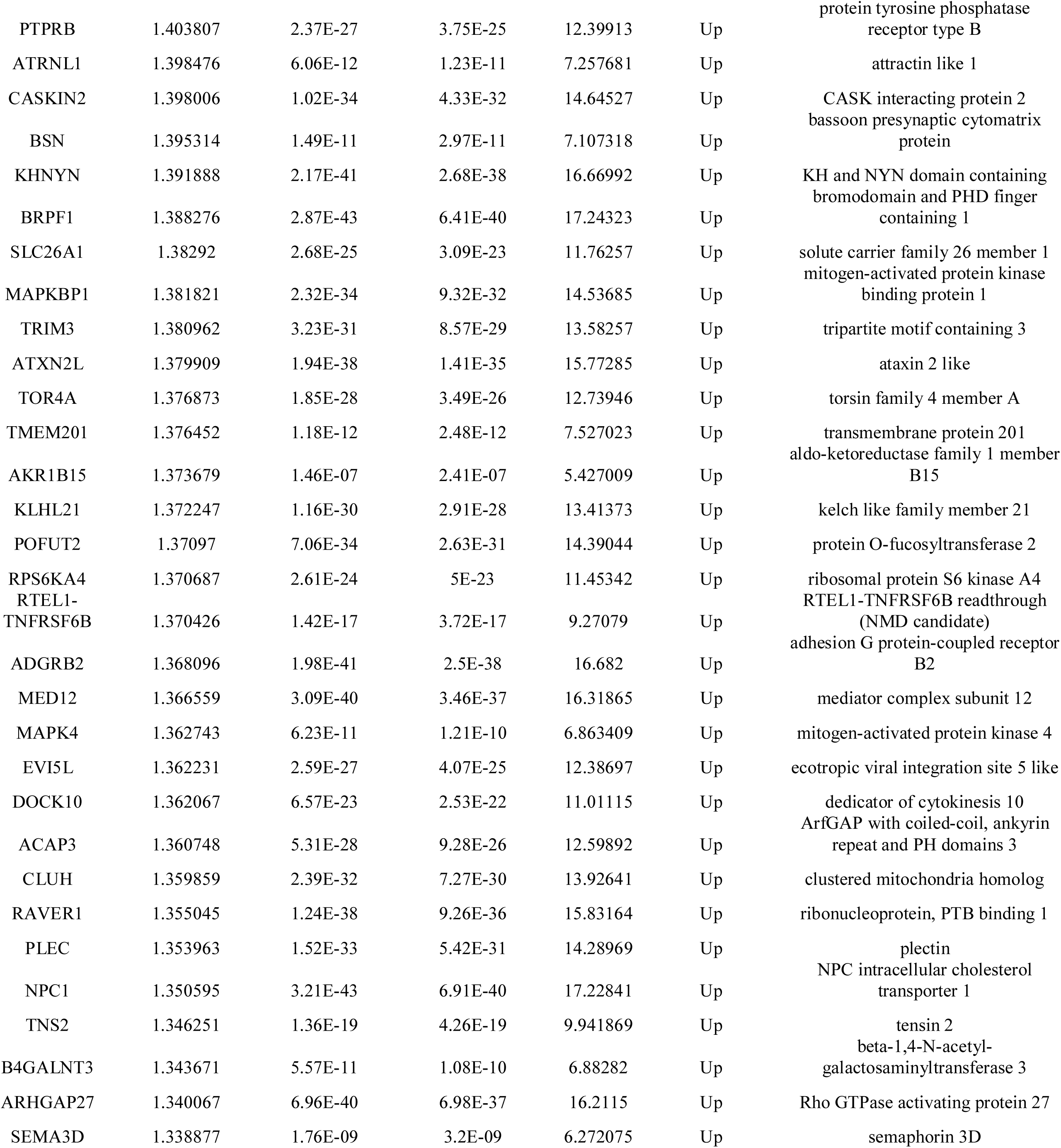
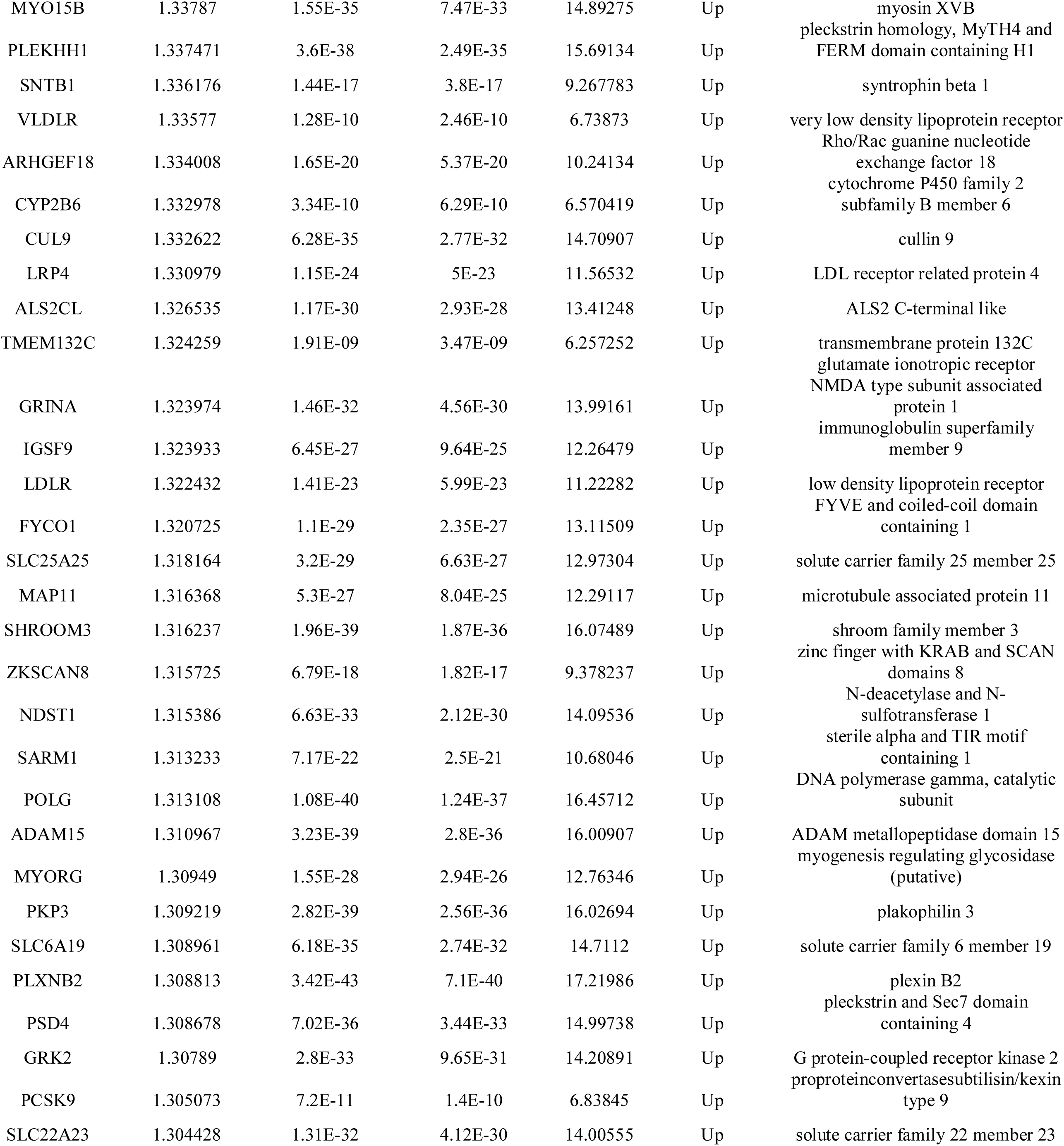
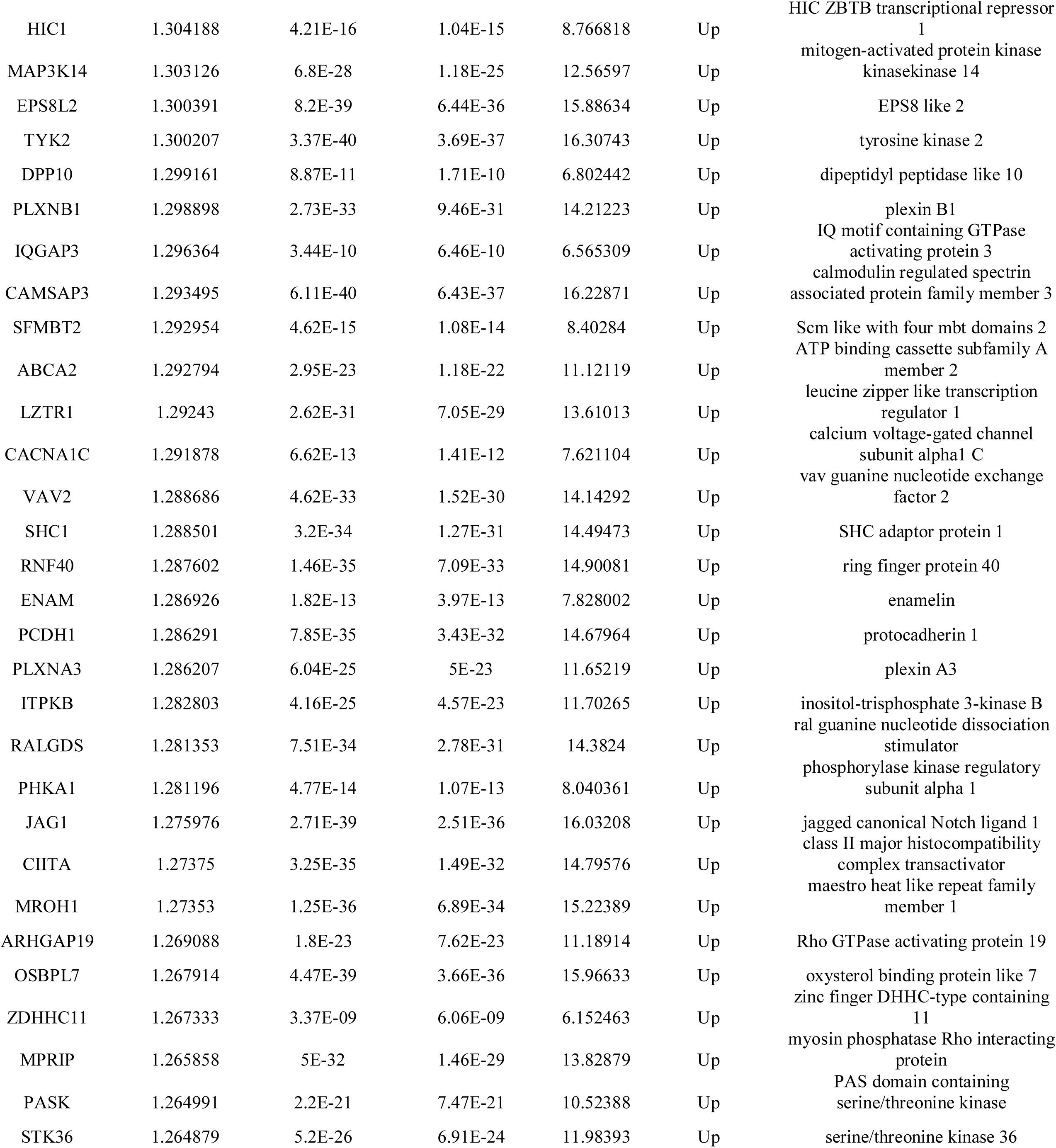
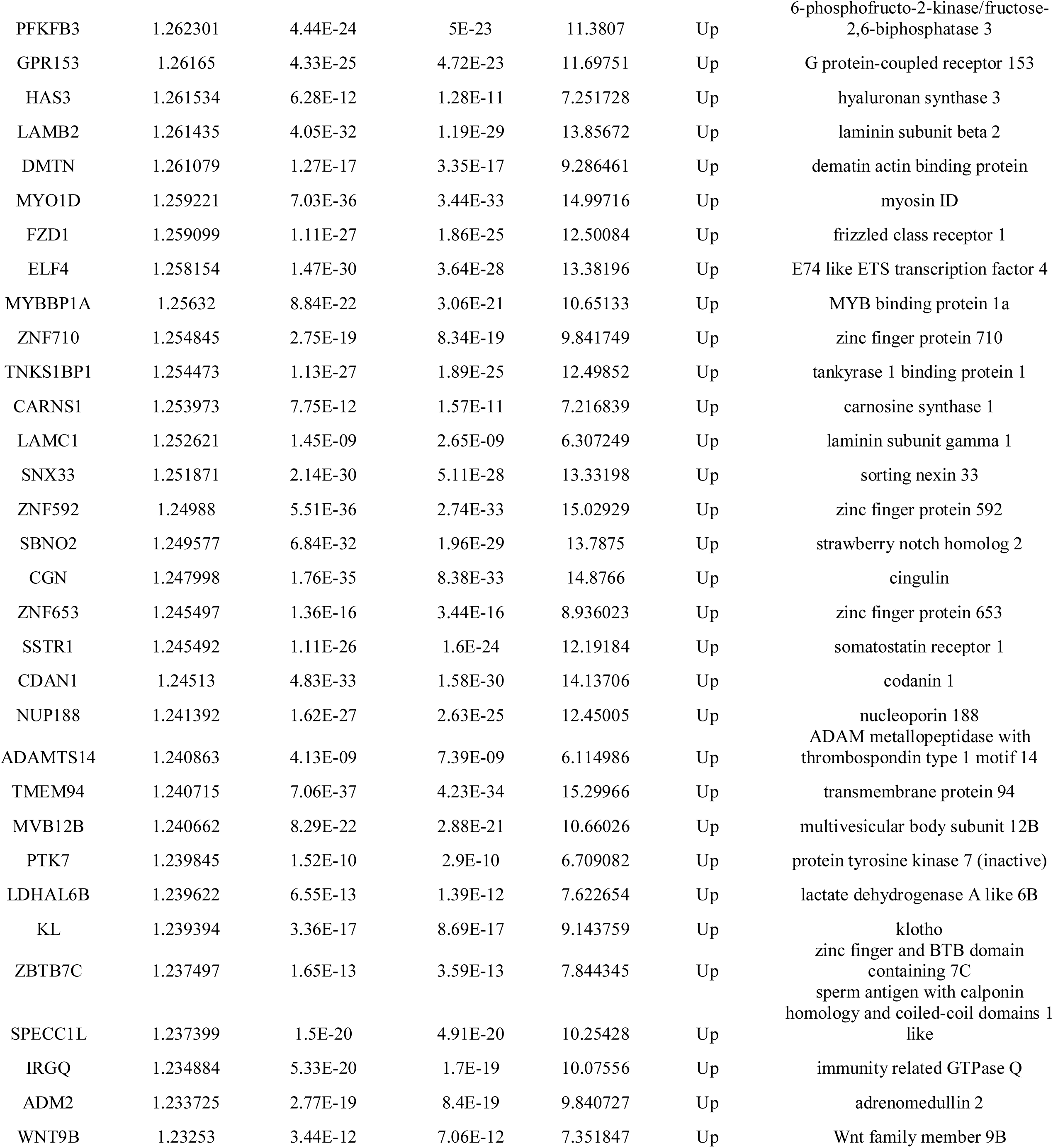
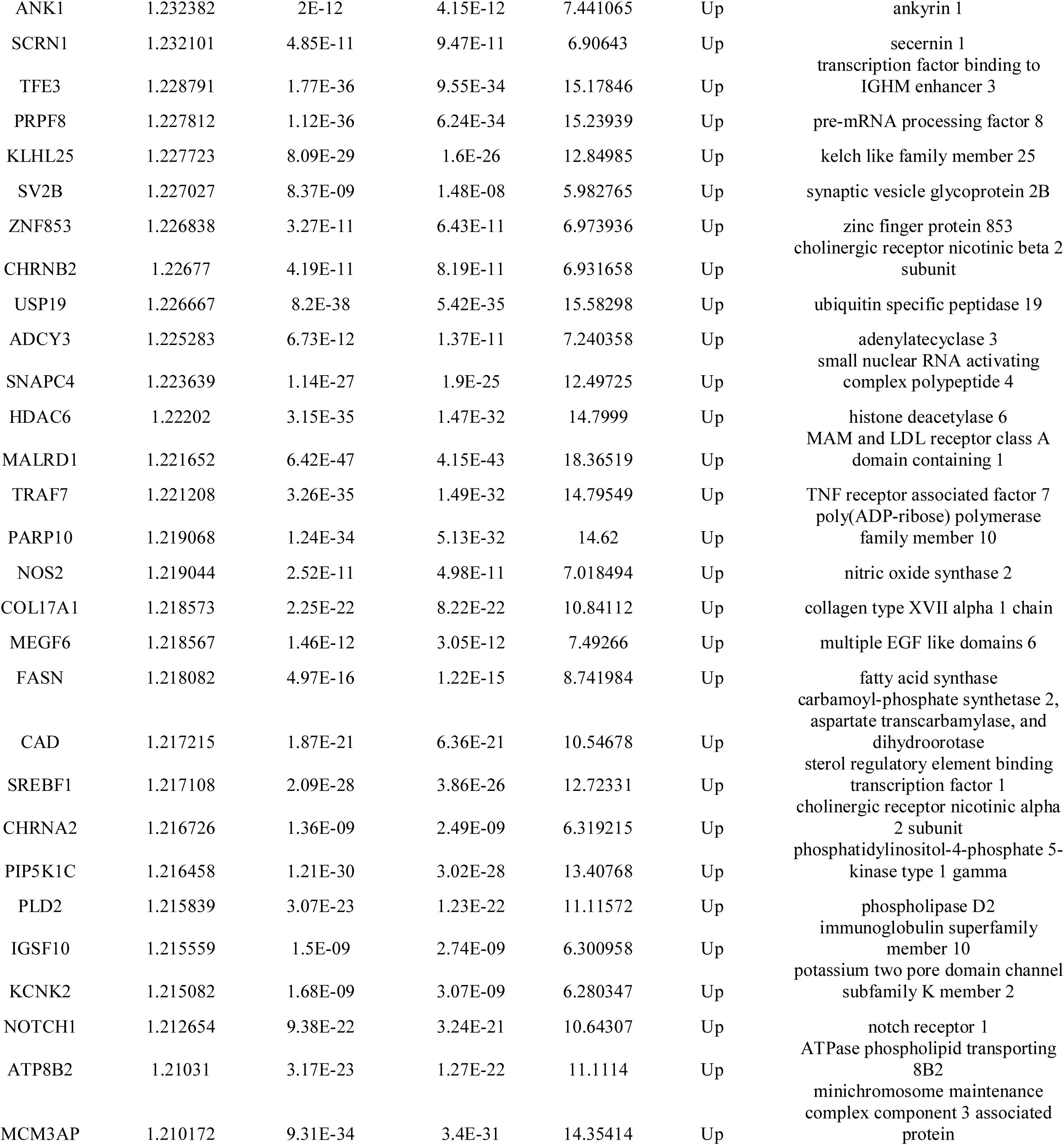
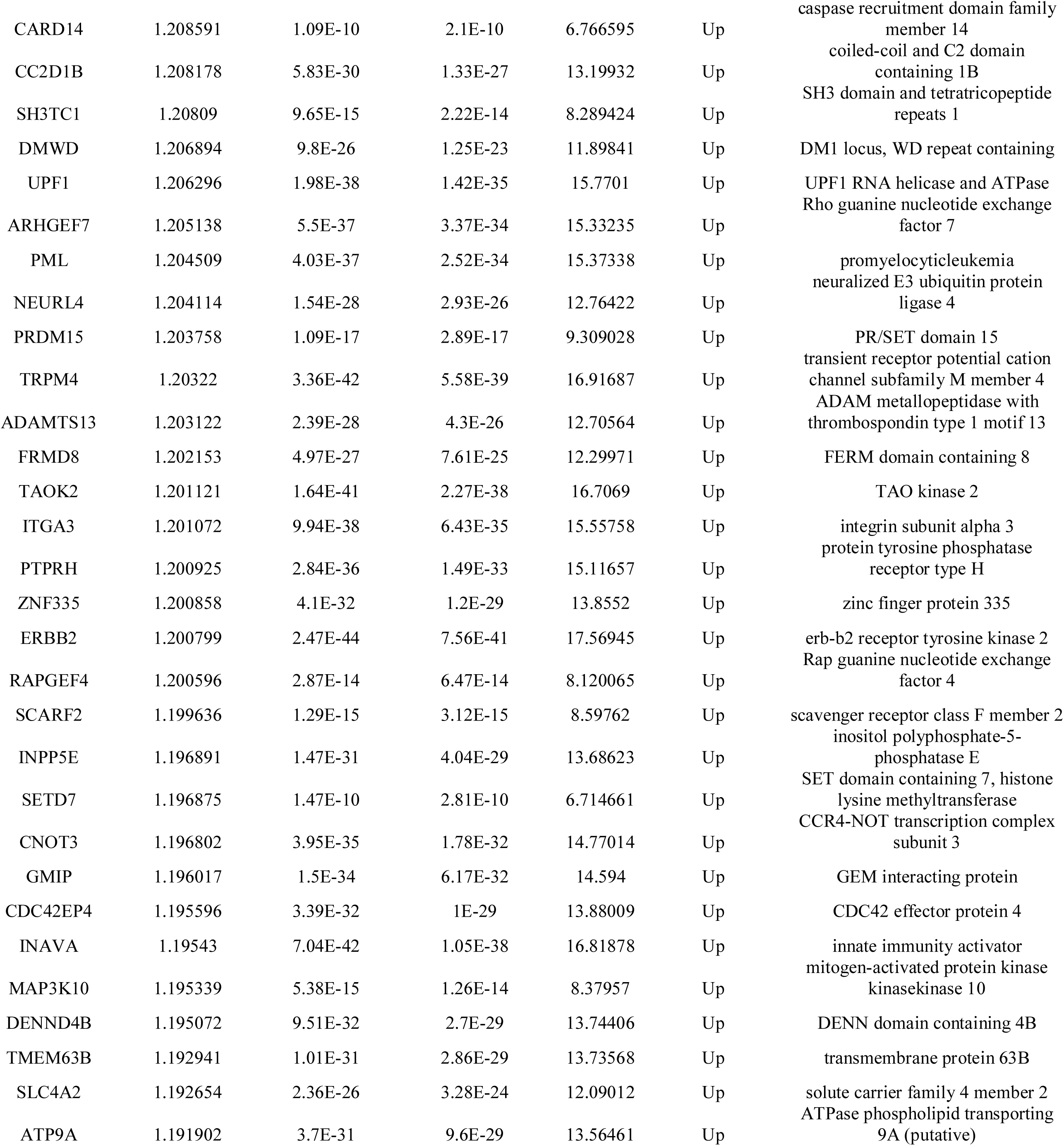
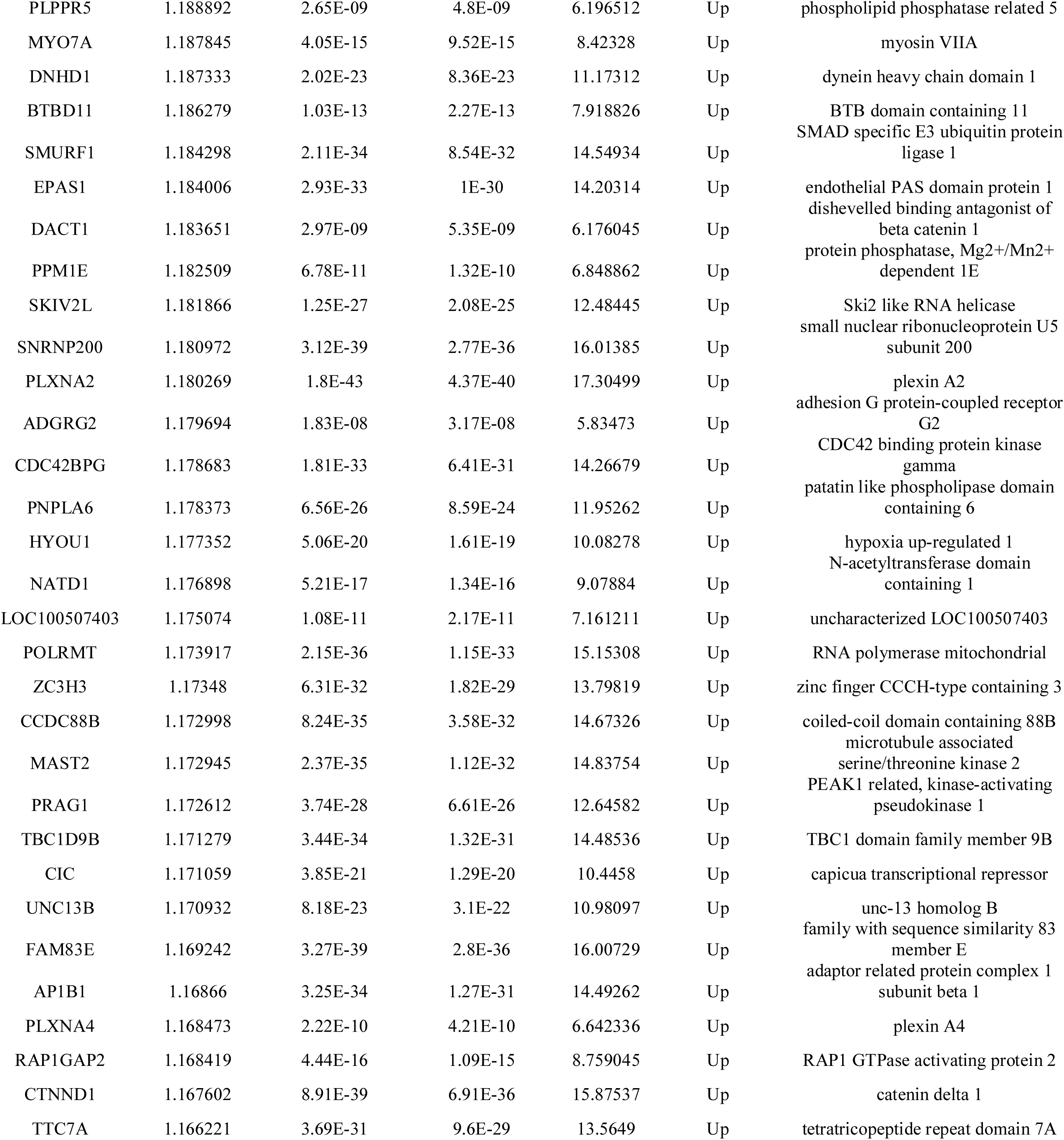
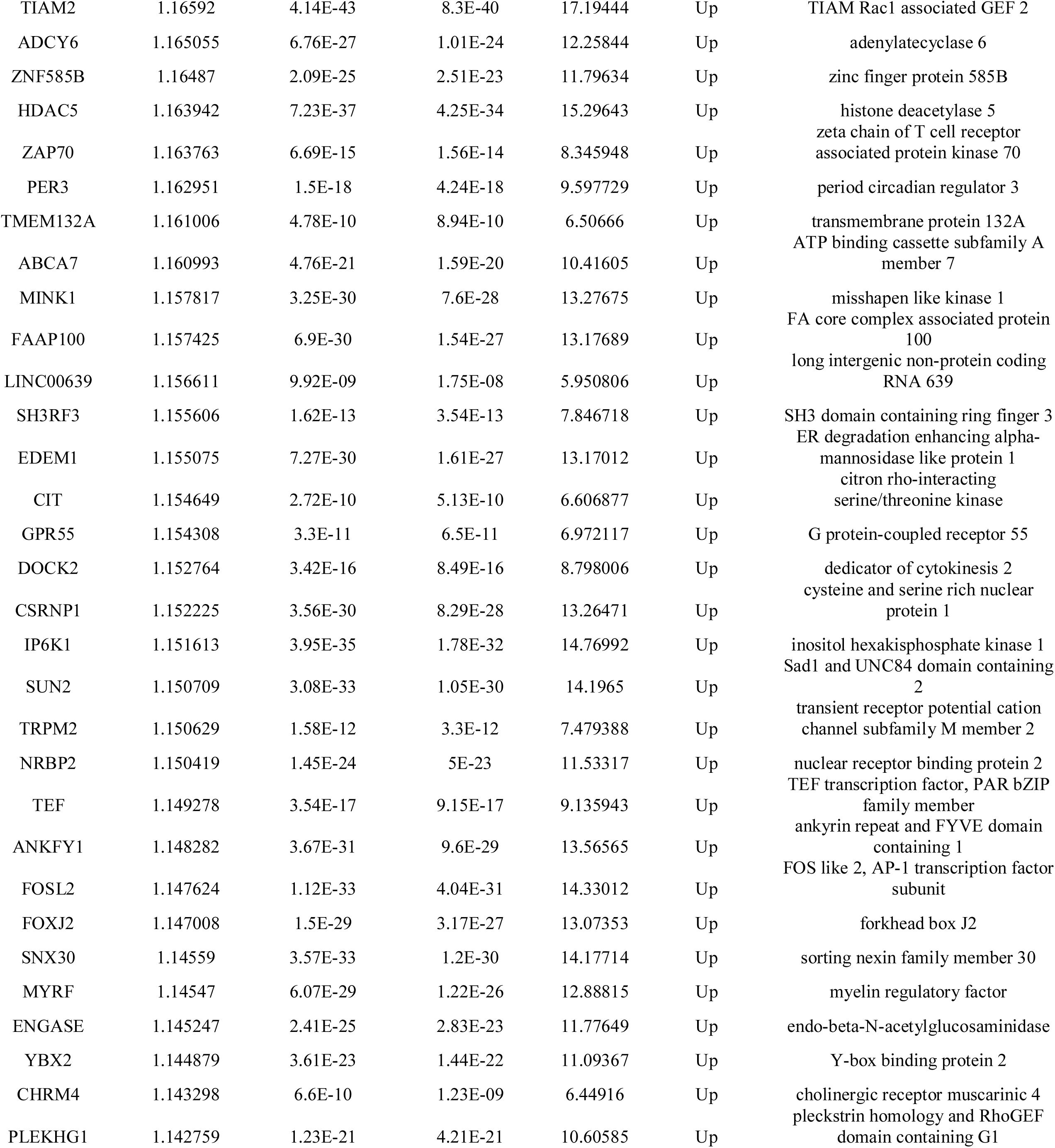
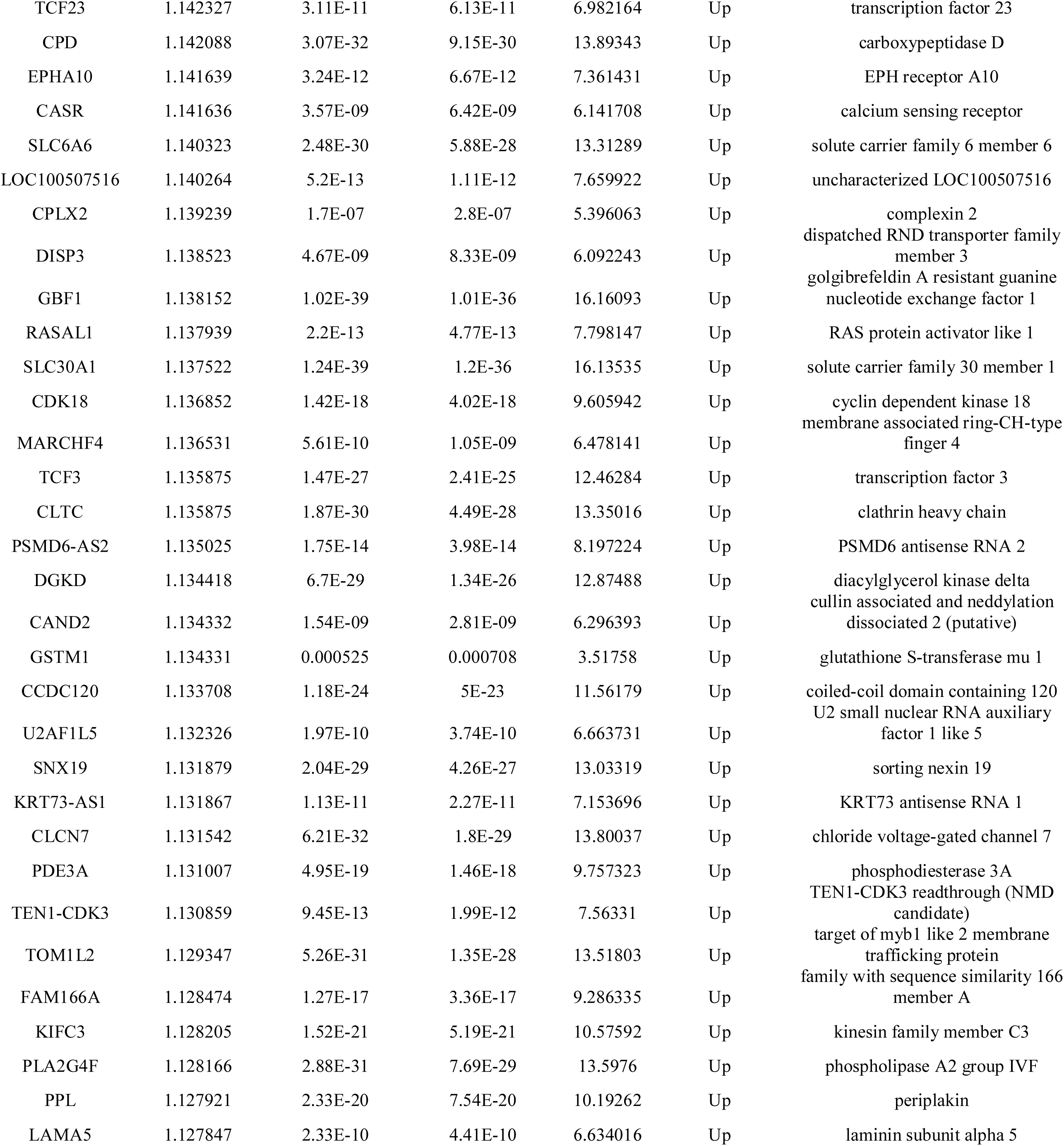
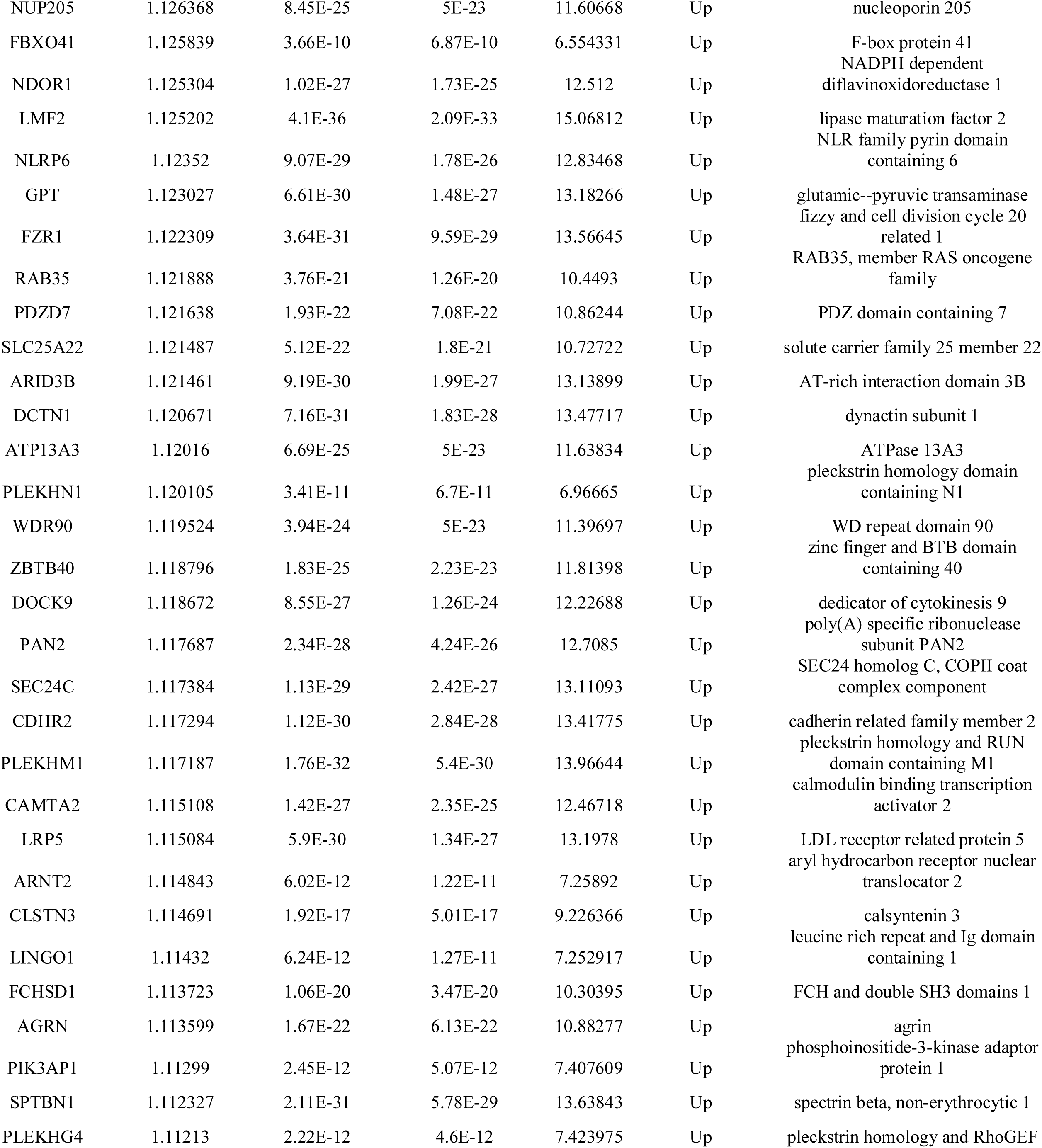
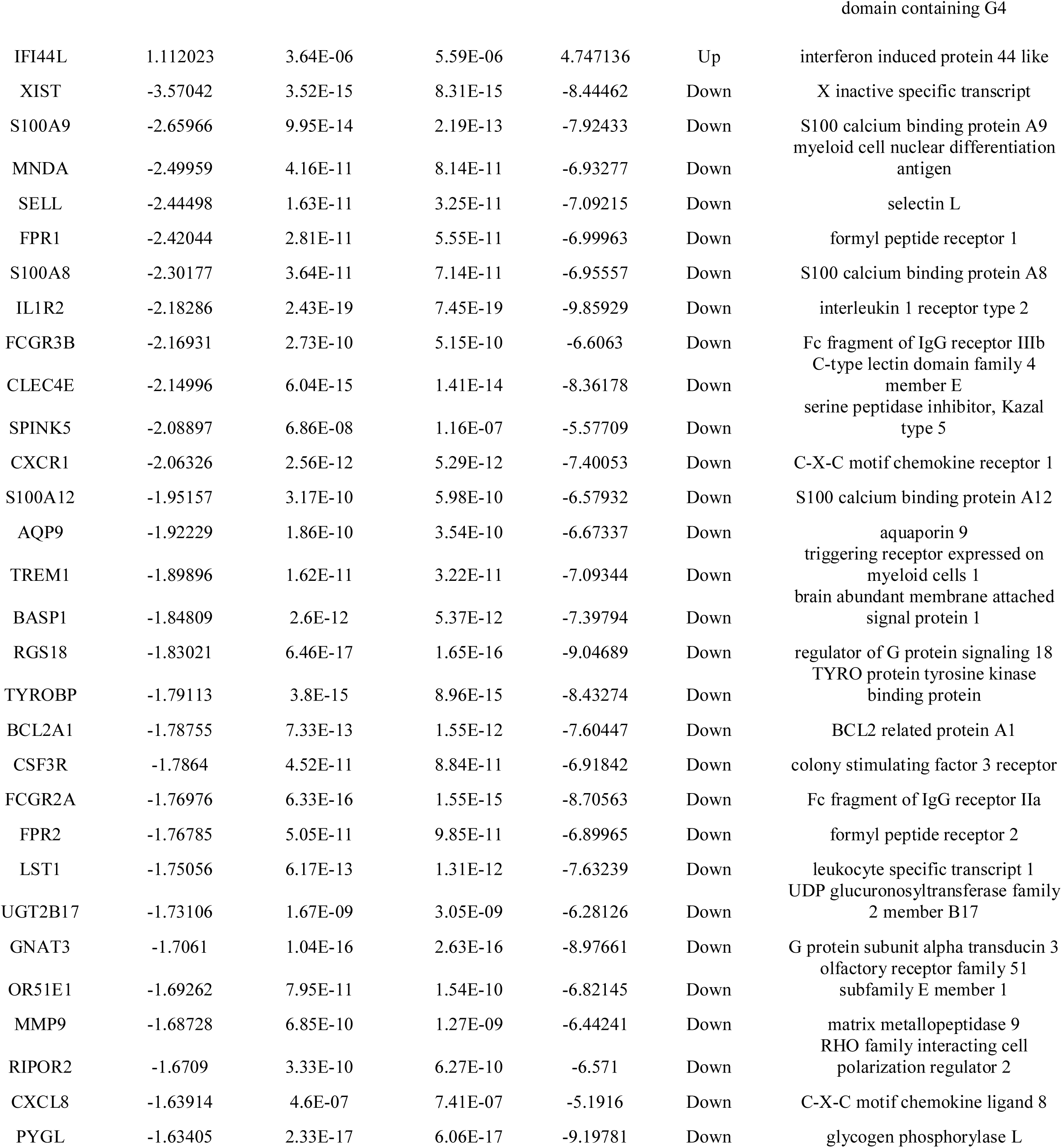
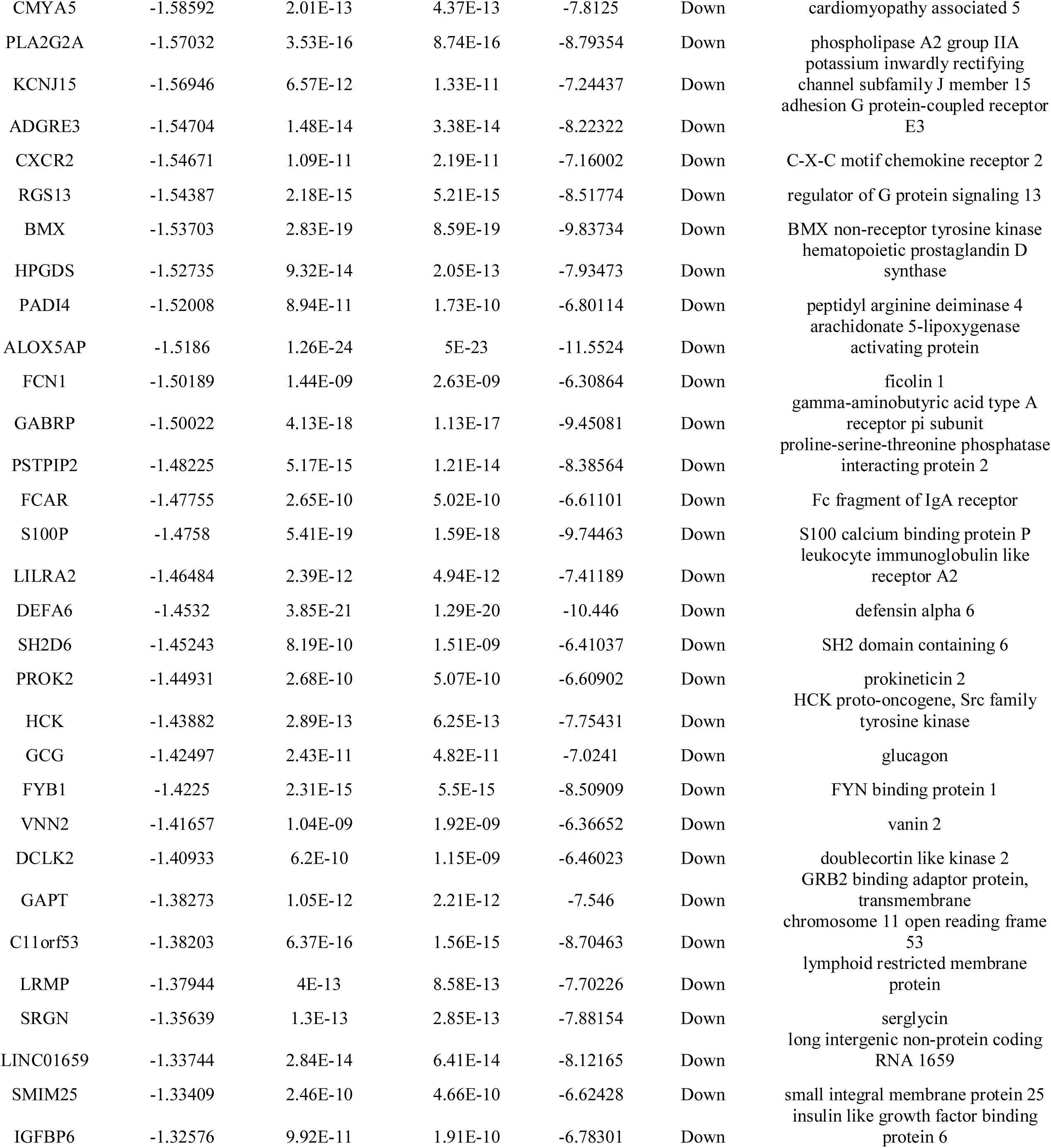
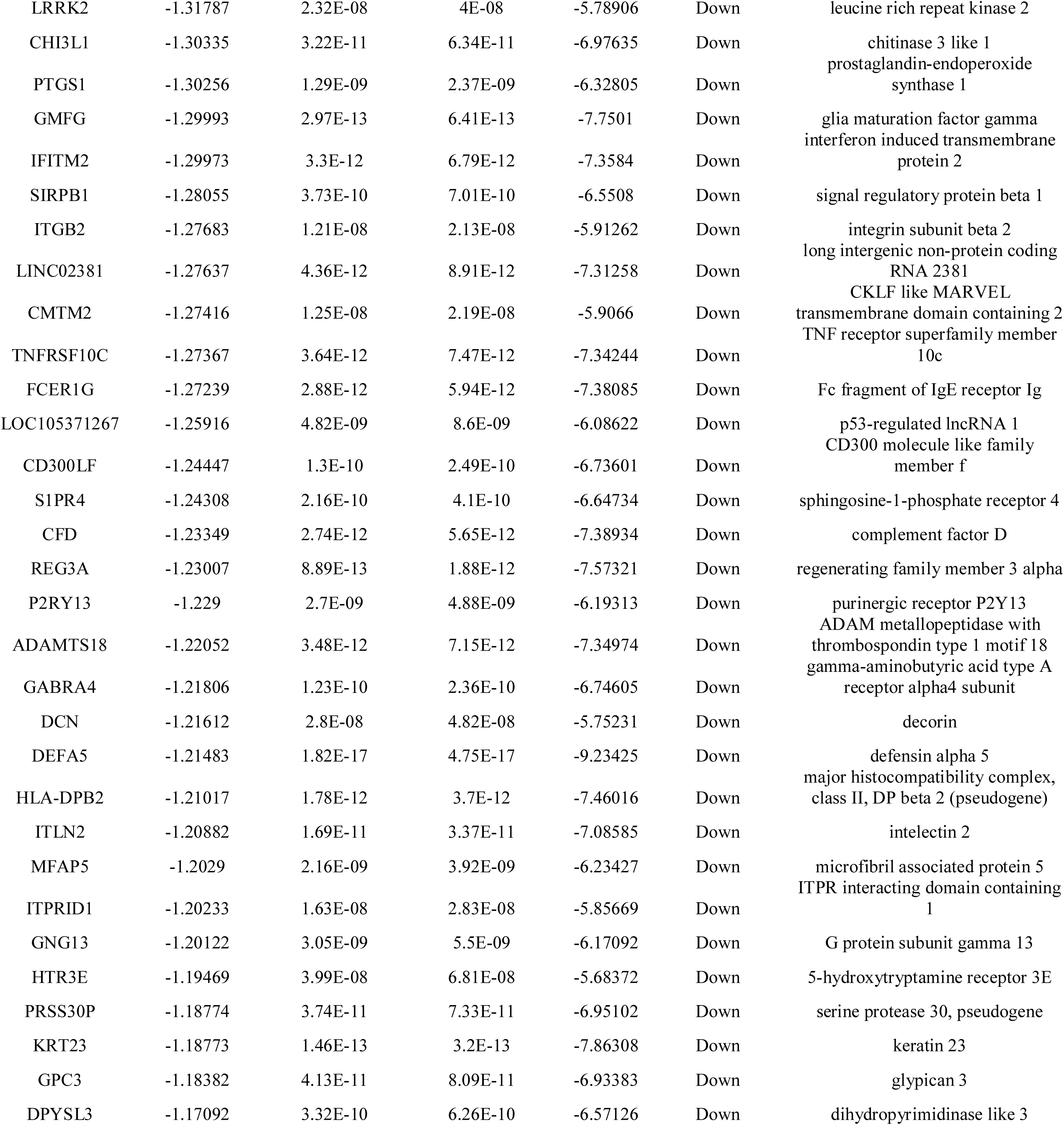
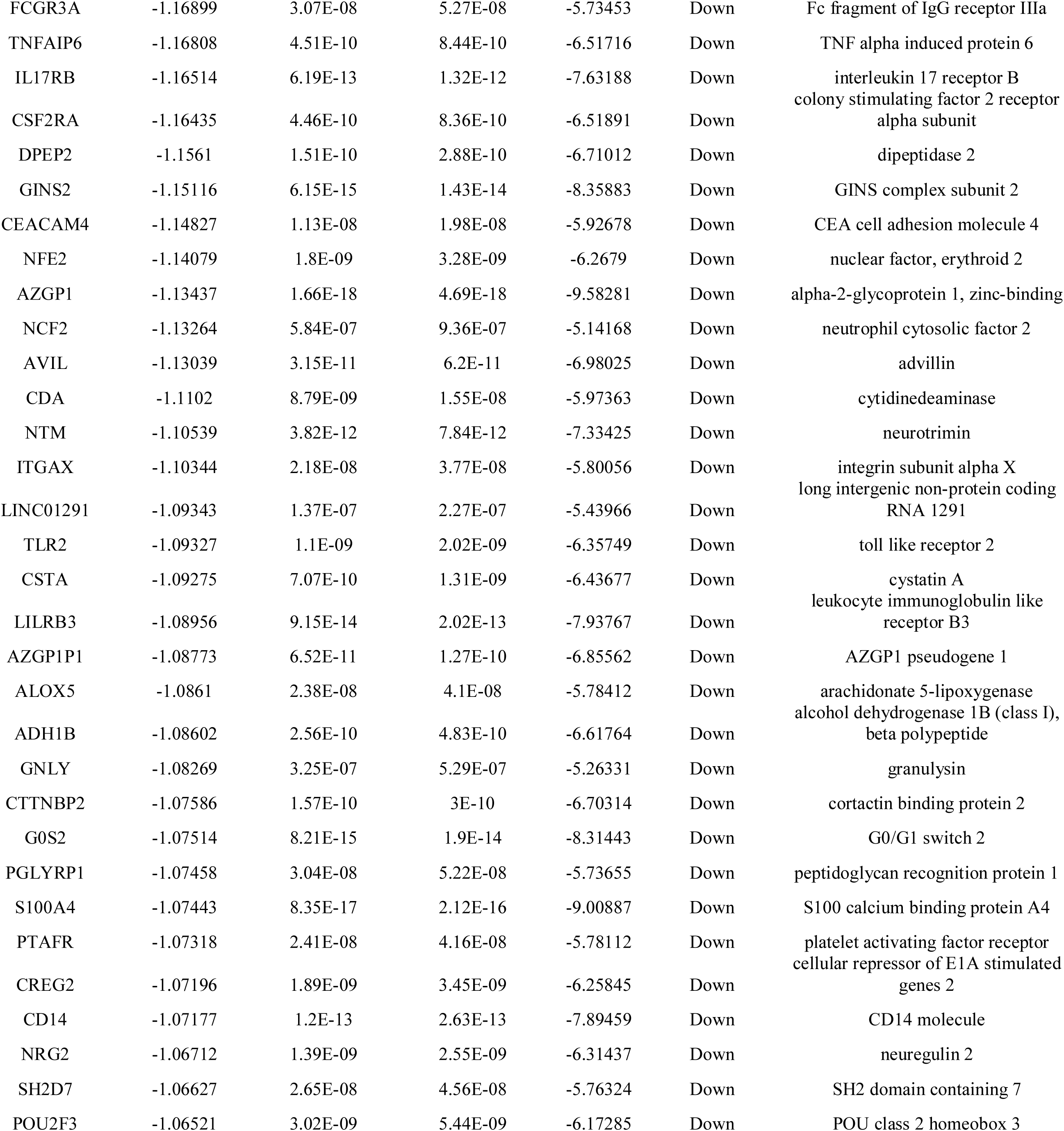
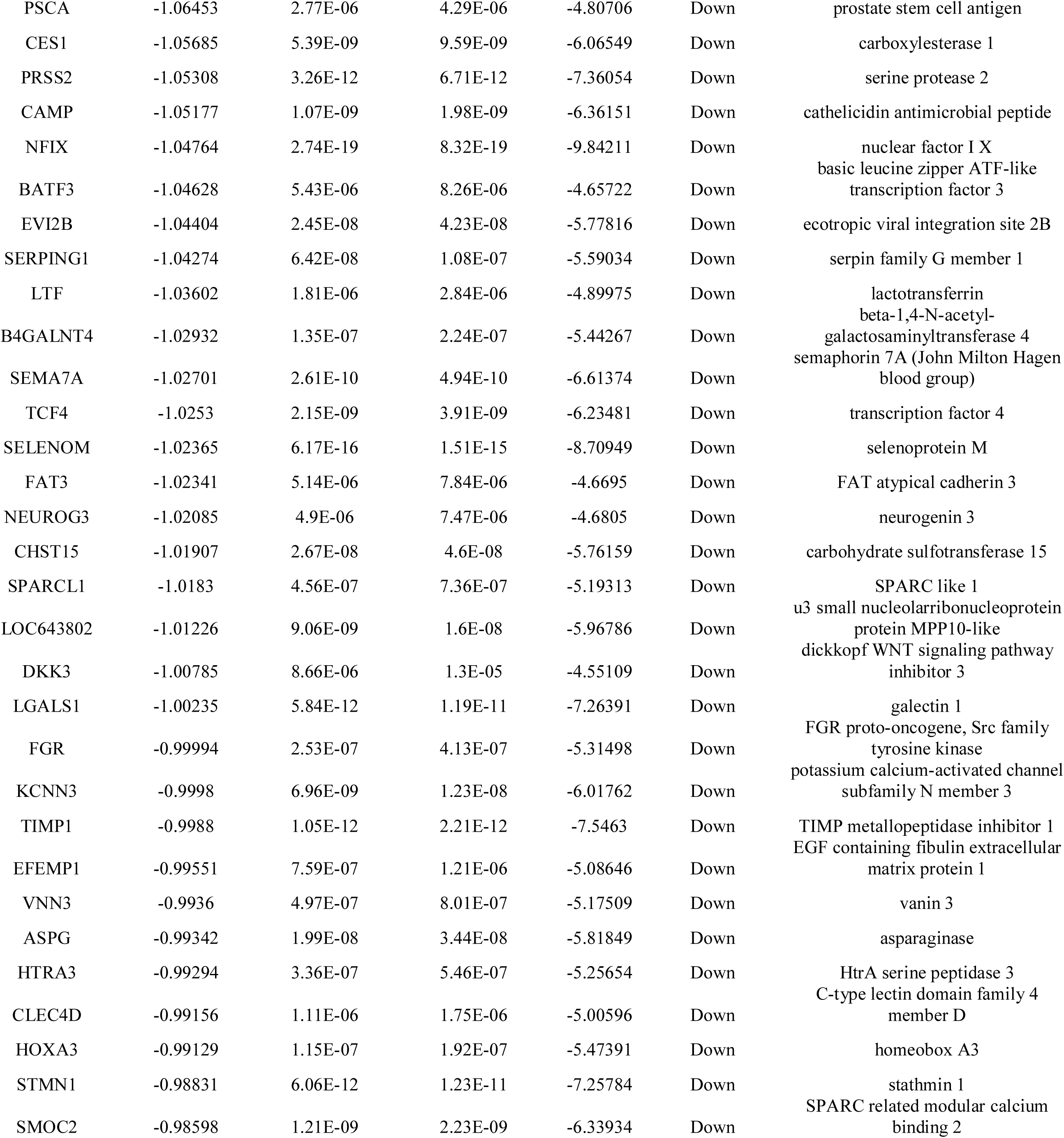
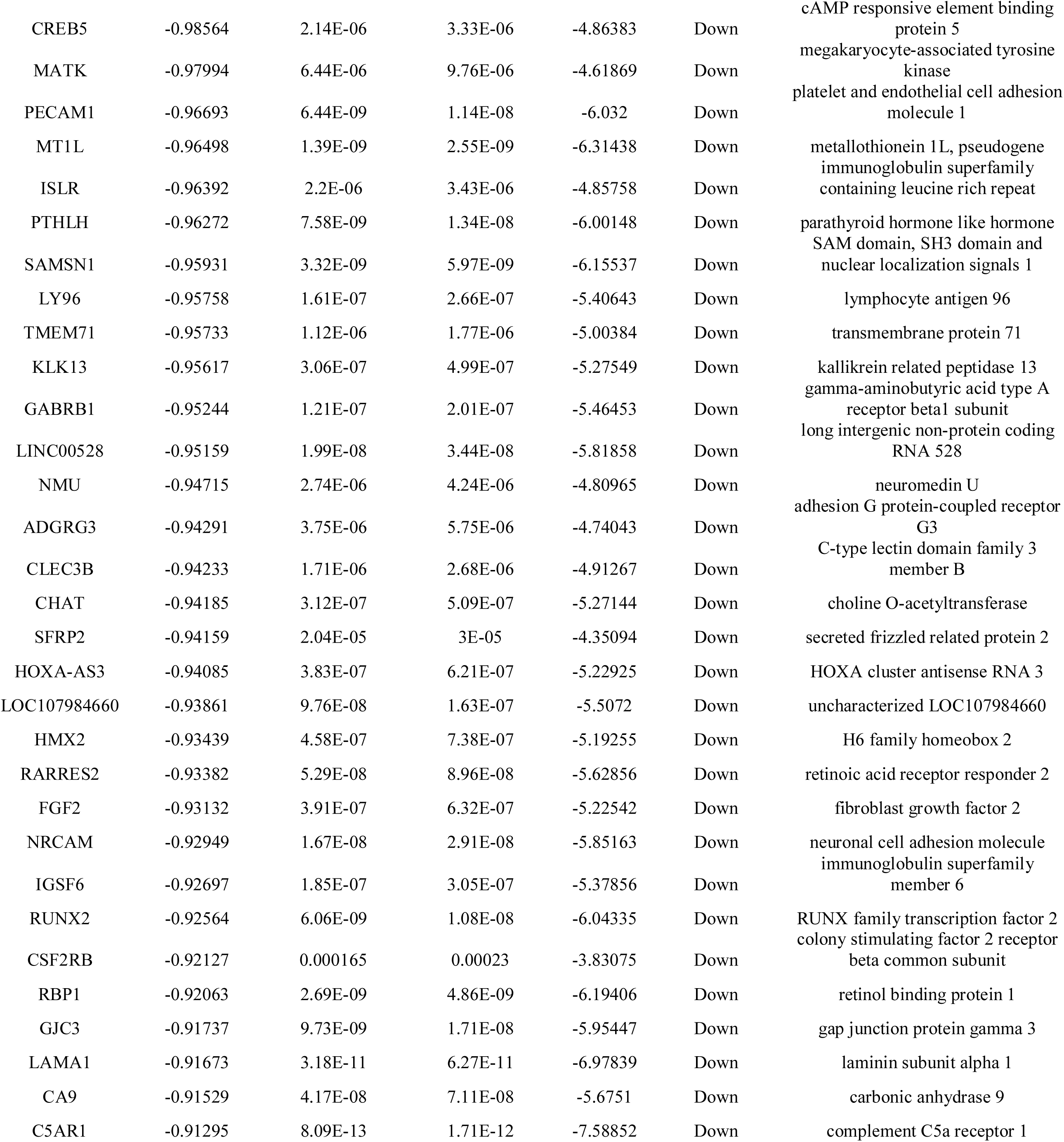
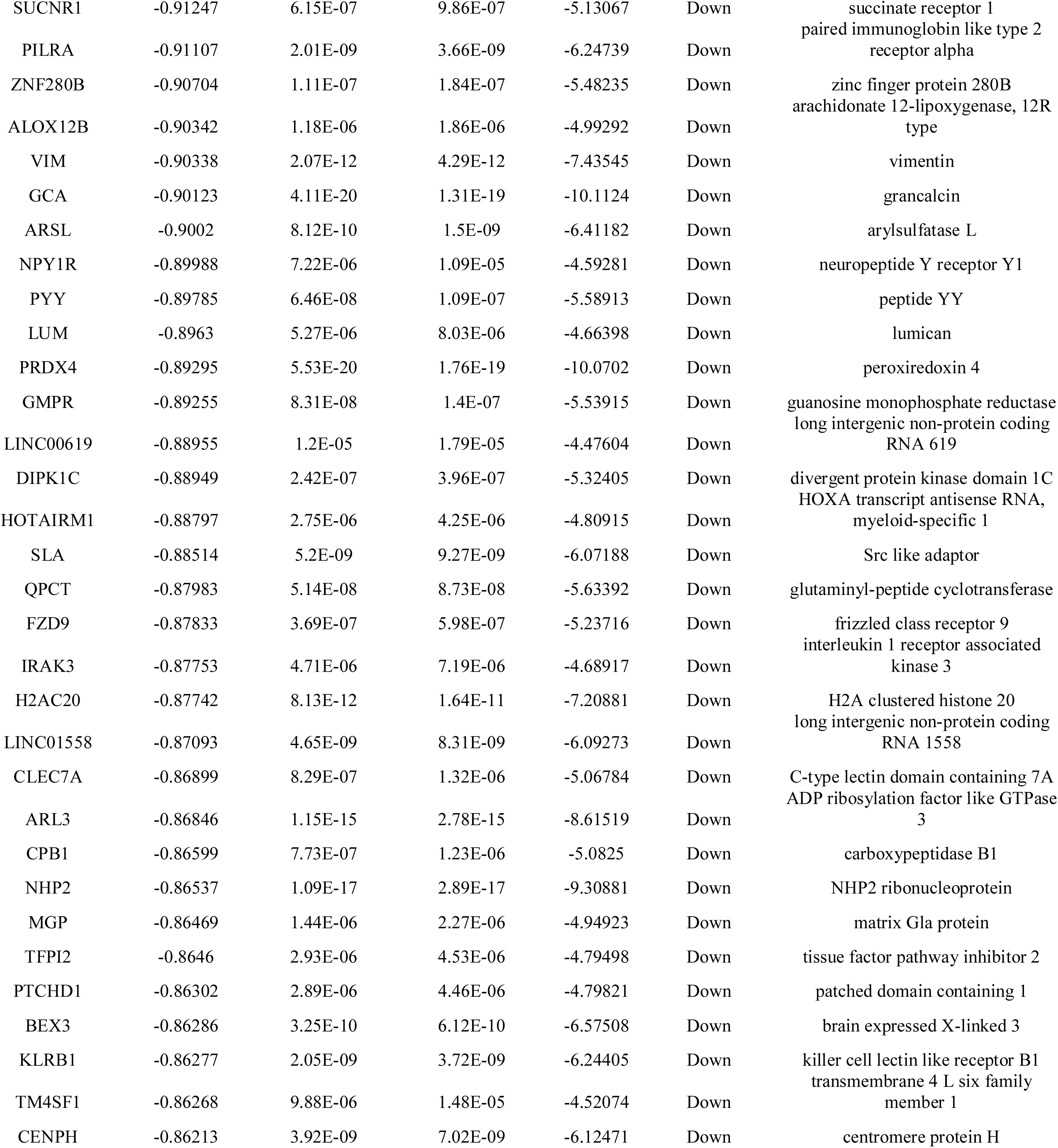
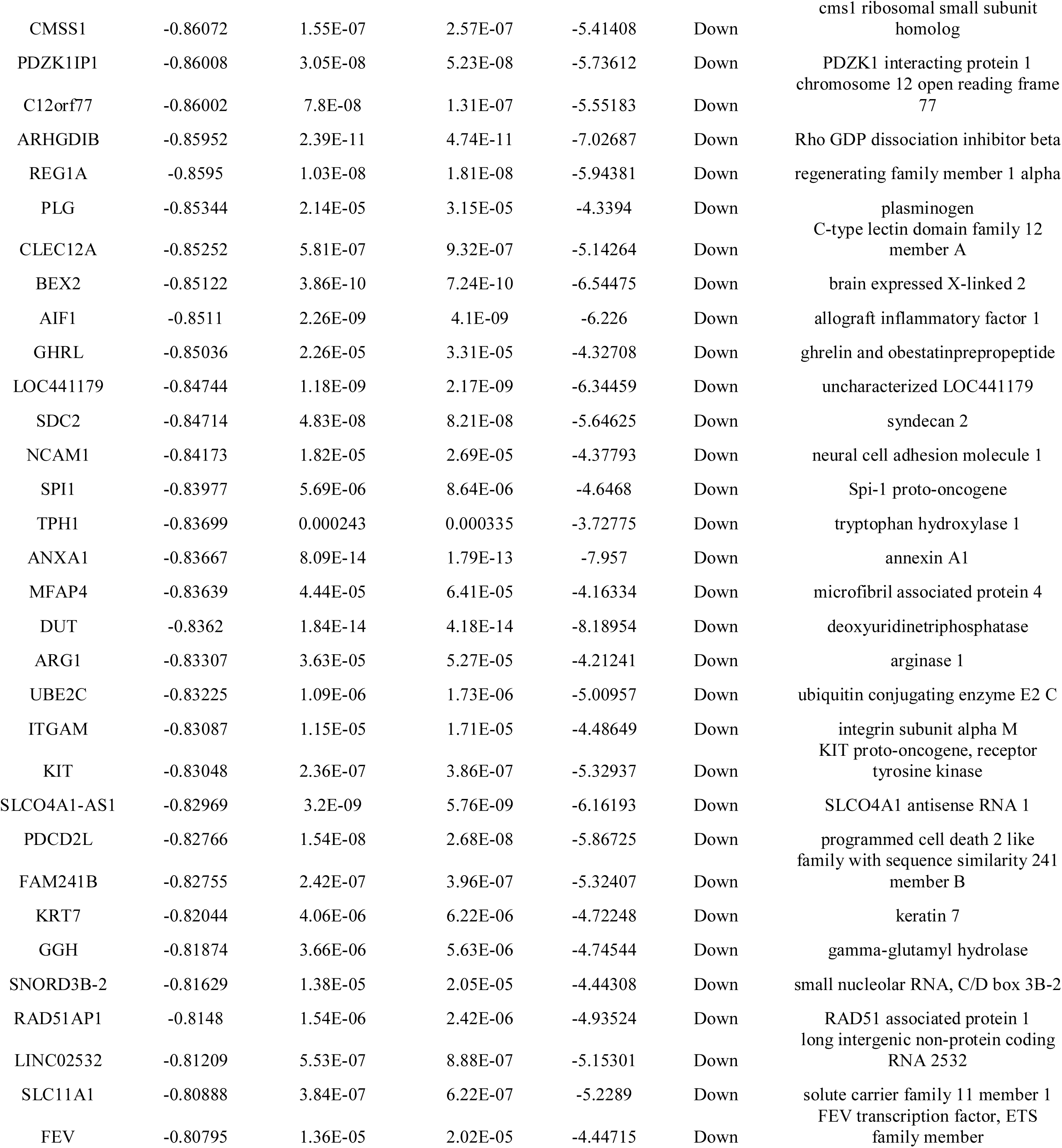
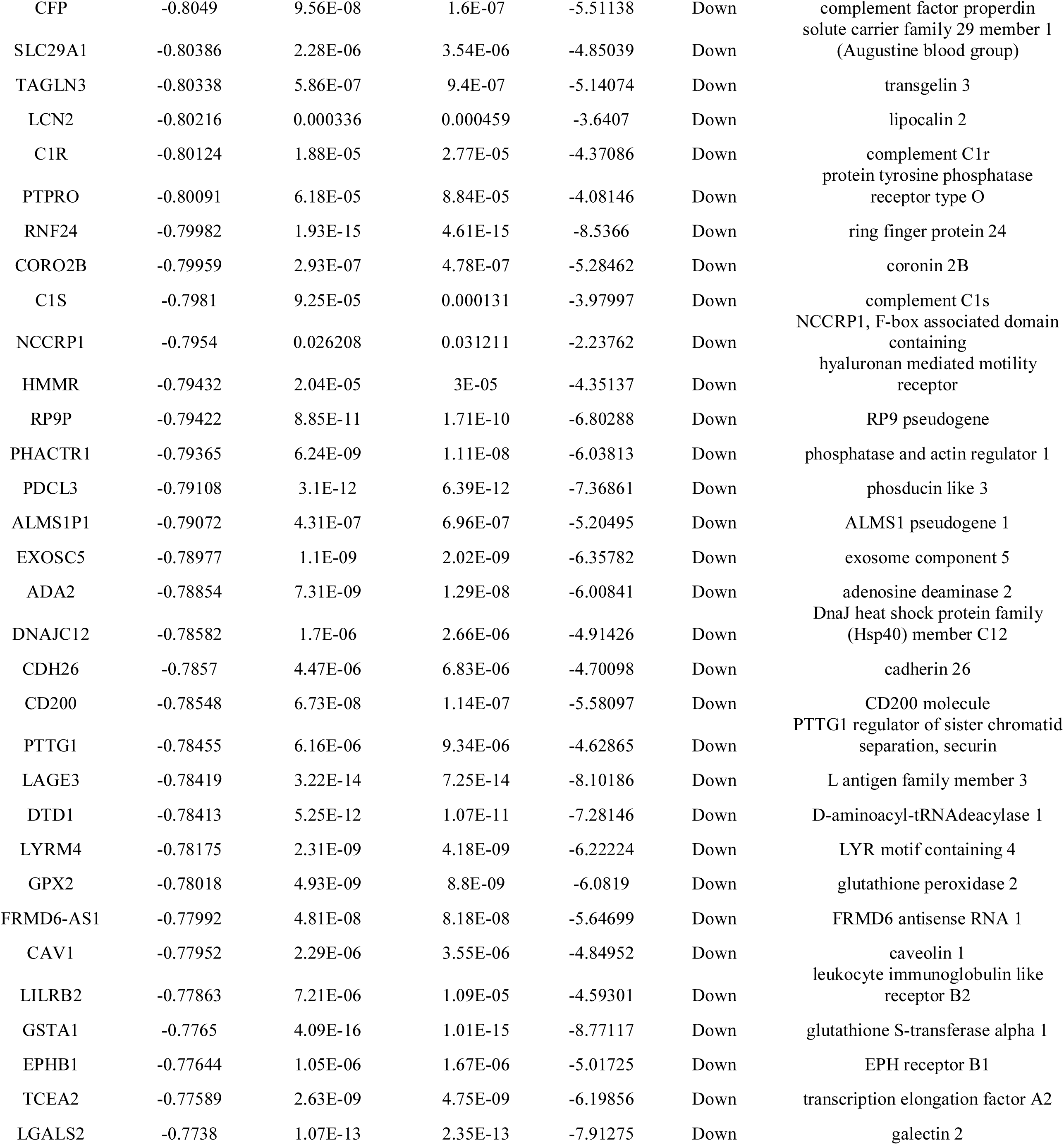
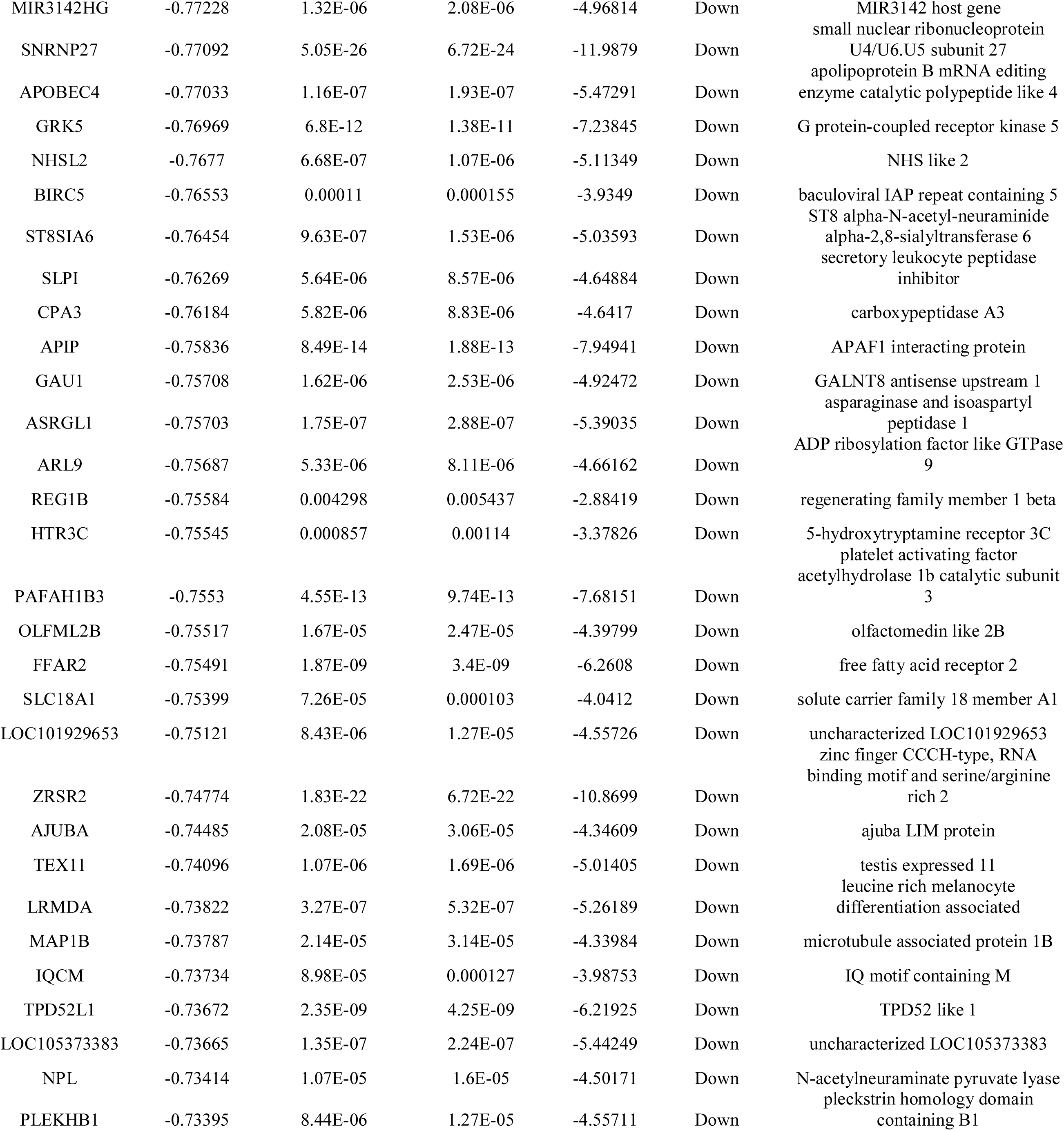
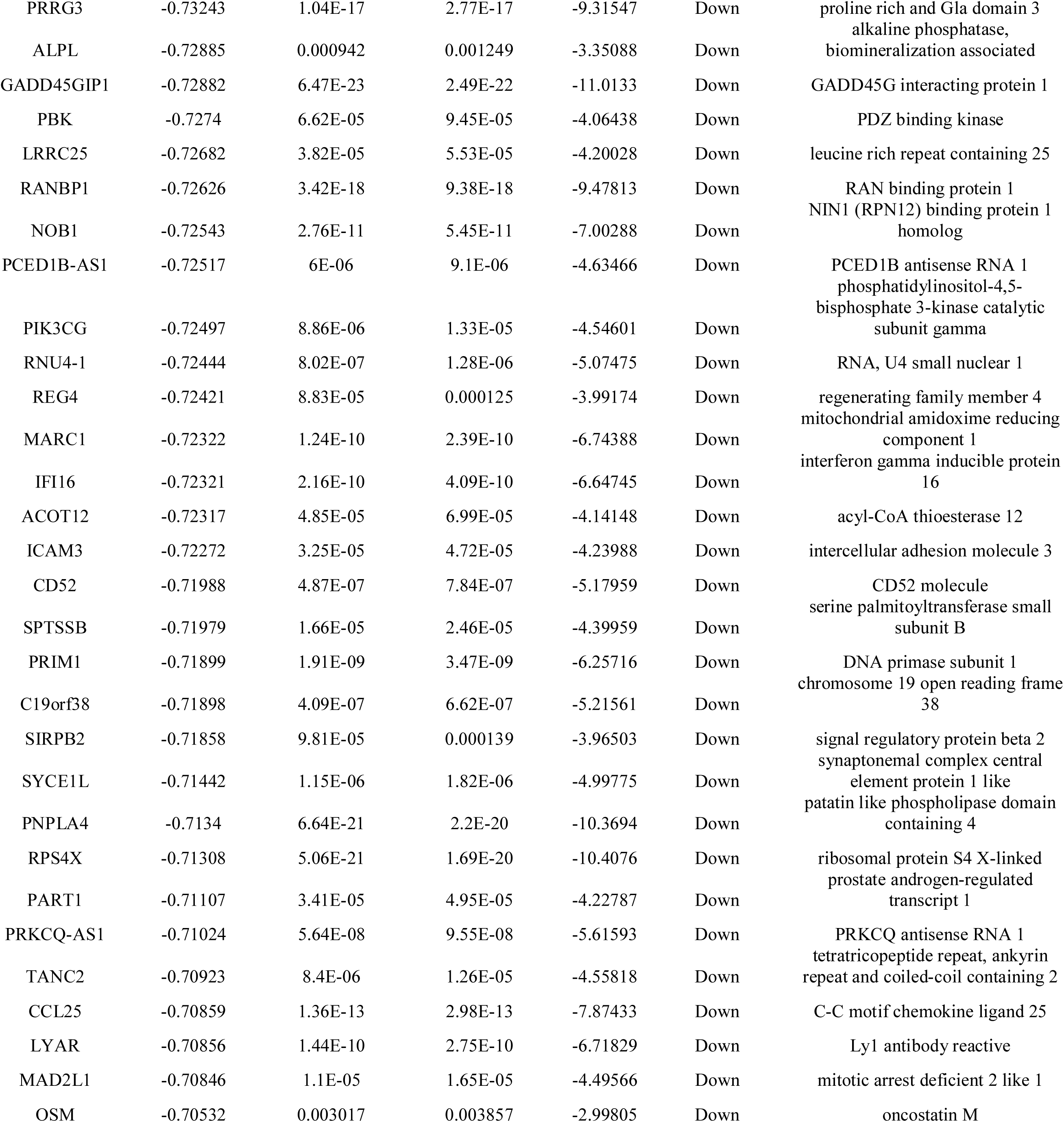
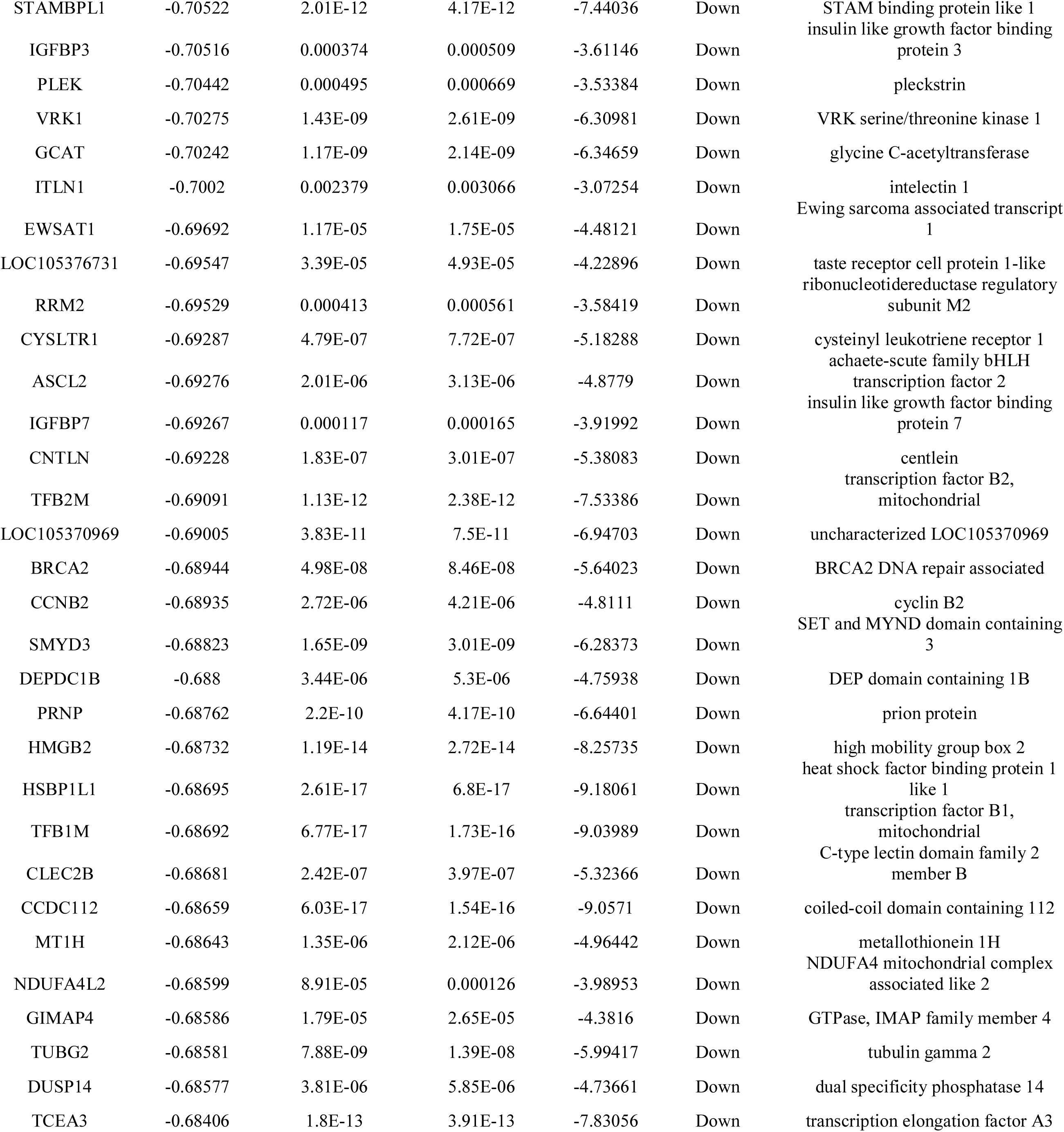
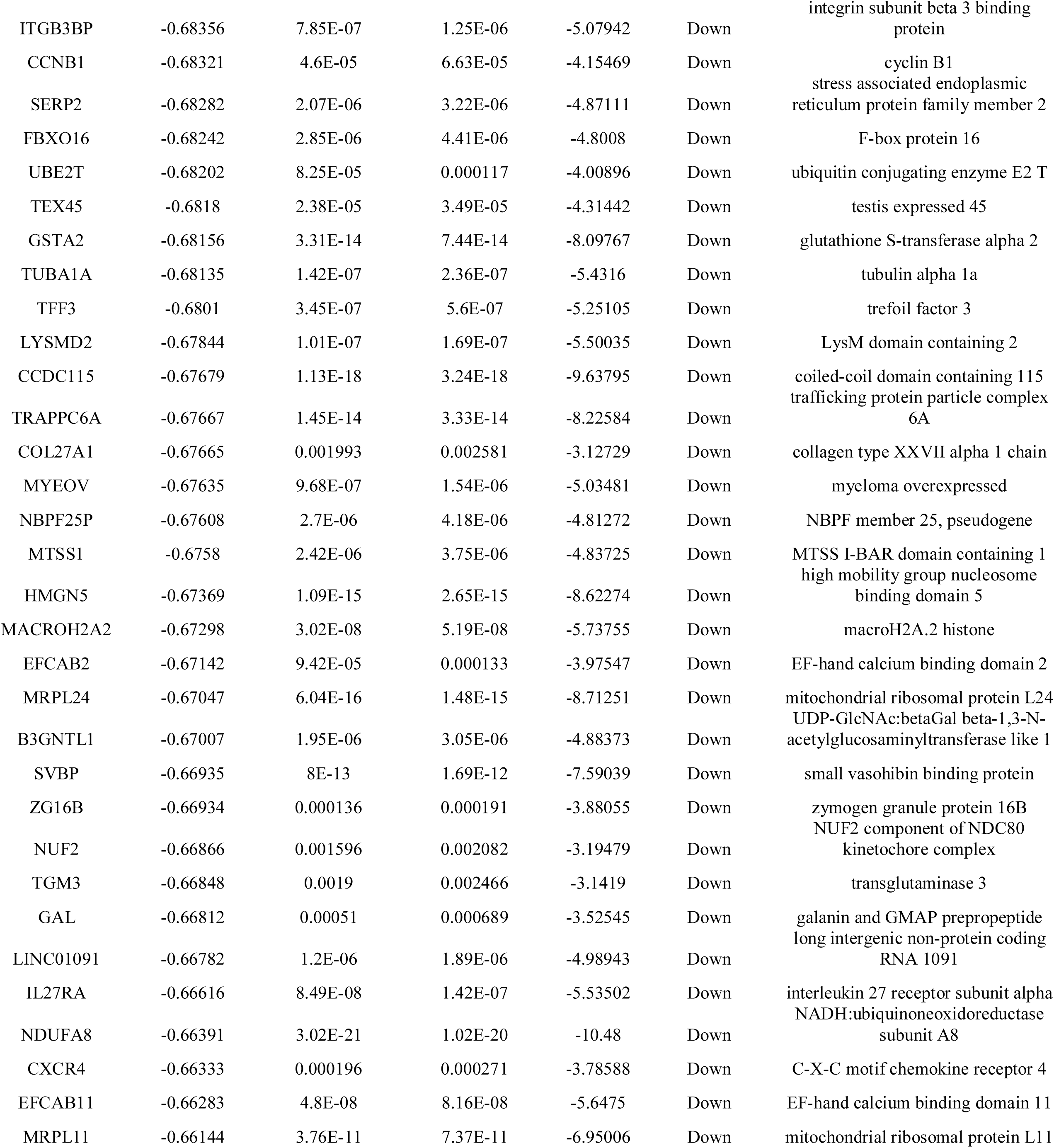
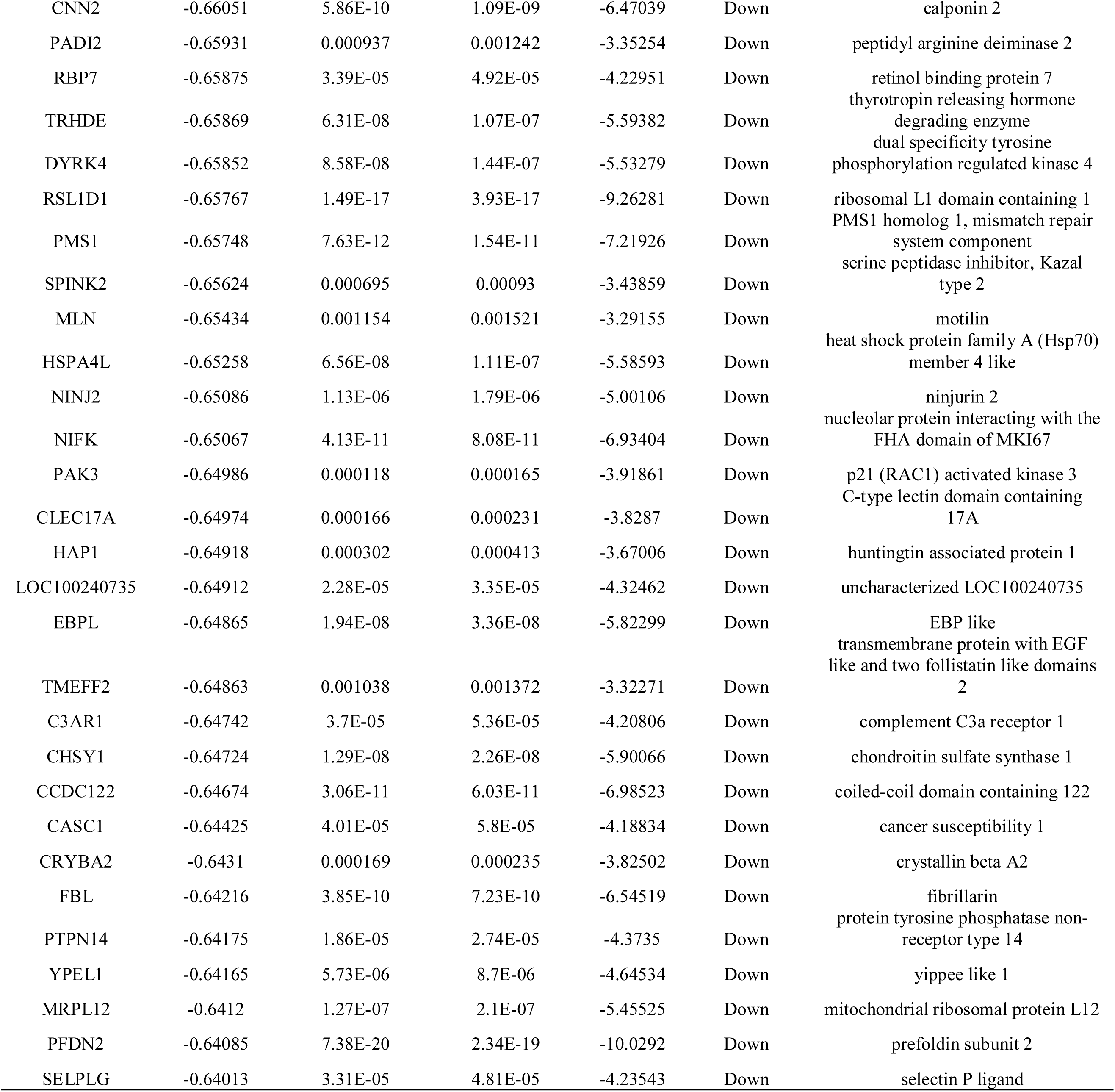
The statistical metrics for key differentially expressed genes (DEGs)

### Molecular docking studies

Using perpetual software module SYBYL-X 2.0, Surflex-Docking docking studies were conducted on active constituents. The molecules were sketched using ChemDraw software, imported and saved into sdf. format using Open Babelfree software. CLTC (Clathrin heavy chain), DCTN1 (Dynactin subunit 1), ERBB2 (Erb-b2 receptor tyrosine kinase 2), FLNA (Filamin A), MYH9 (Myosin heavy chain 9) and their X-RAY crystallographic structure and co-crystallized PDB code 5M5R, 2HLC, 1MFG, 3HOC and 6LKM respectively were selected for docking and were extracted from Protein Data Bank [29–30]. Optimization of the molecules were performed by transforming the 3D concord structure, applying TRIPOS force field and applying GasteigerHuckel (GH) charges, In addition, MMFF94s and MMFF94 algorithm processes have been used for energy minimization. Protein processing was performed after introduction of the protein. The co-crystallized ligand and all the water molecule were ejected from the crystal structure; added more of hydrogen and refined the side chain. To minimize structure complexity, the TRIPOS force field was used and the interaction efficiency of the compounds with the receptor was represented by the Surflex-Dock score in kcal/mol units. The best position was inserted into the molecular area between the protein and the ligand. Using Discovery Studio Visualizer, the simulation of ligand interaction with receptors is accomplished

## Results

### Identification of DEGs

According to the cut-off criteria of a P-value <0.05, and |logFC| > 1.112 for up regulated genes and |logFC| < −0.64 for down regulated genes, a total of 872 DEGs were obtained, including 439 up regulated and 433 down regulated genes and are listed in Table 1. The volcano plot is presented in Fig. 1. The heat map of hierarchical clustering analysis for the DEGs were showed in Fig.2.

**Fig. 1.**
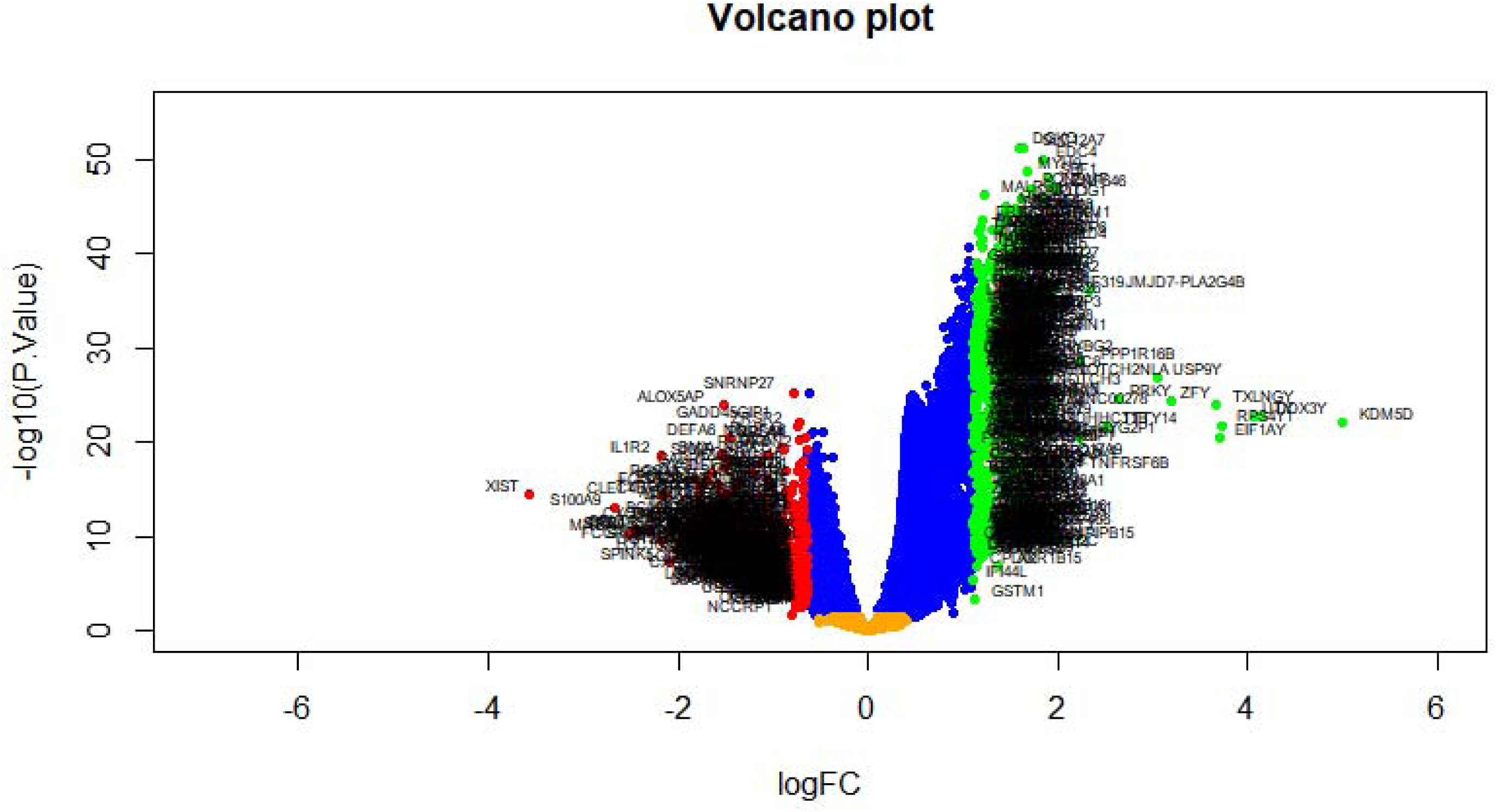
Volcano plot of differentially expressed genes. Genes with a significant change of more than two-fold were selected. Green dot represented up regulated significant genes and red dot represented down regulated significant genes.

**Fig. 2.**
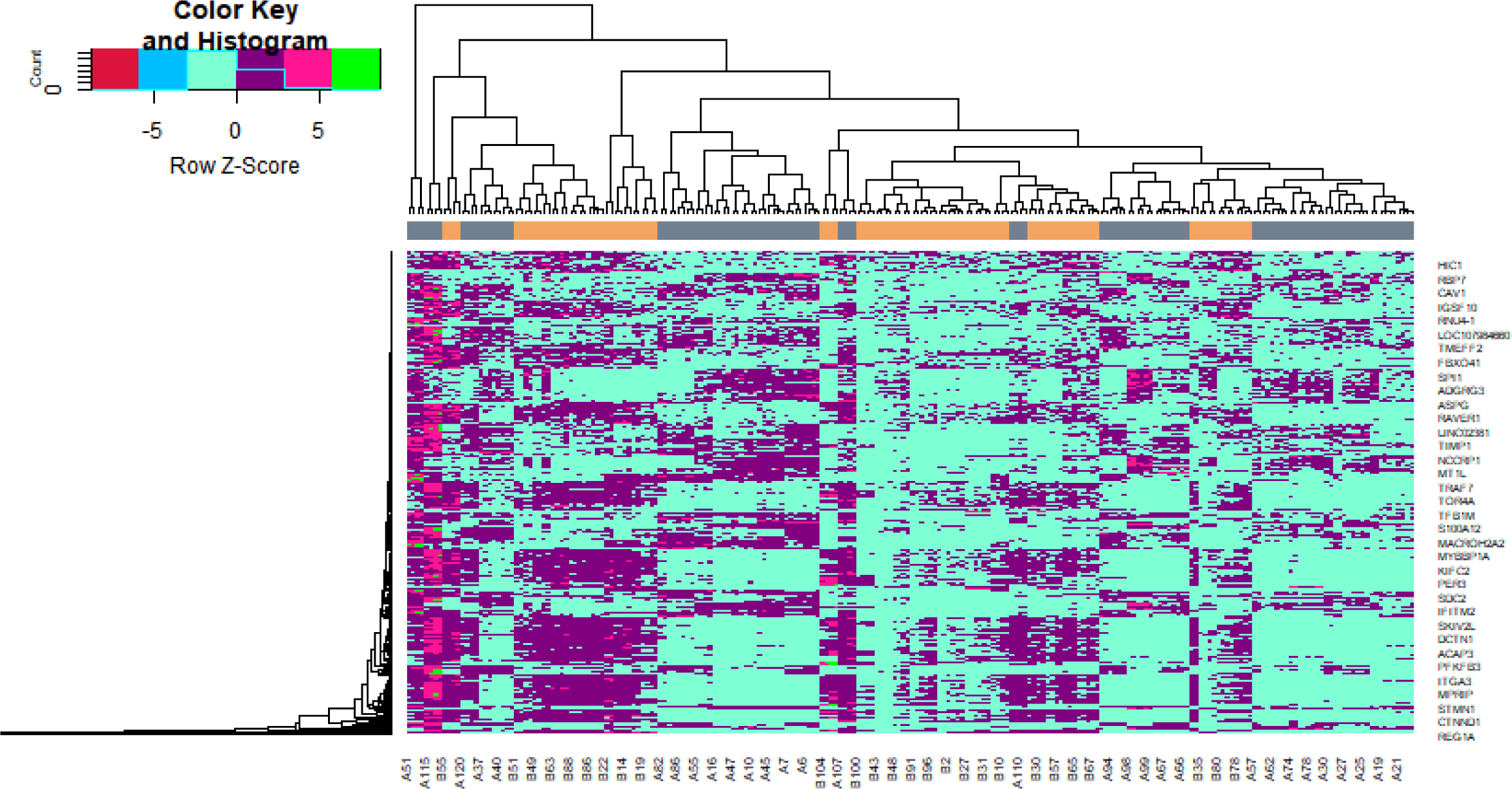
Heat map of differentially expressed genes. Legend on the top left indicate log fold change of genes. (A1 – A120 = non diabetic obese samples; B1 – B104 = diabetic obese samples)

### Gene Ontology (GO) and Reactome pathway enrichment analysis of DEGs

The enriched GO terms and REACTOME pathways for DEGs were identified. According to the P-values, the top enriched terms were listed in Table 3 and Table 4. In the whole up regulated genes, BP terms (plasma membrane bounded cell projection organization and neurogenesis), CC terms (neuron projection and golgi apparatus), MF terms (drug binding and ribonucleotide binding), and pathways (axon guidance and extracellular matrix organization.) were enriched. In the down regulated genes, BP terms (cell activation and secretion), CC terms (secretory granule and secretory vesicle), MF terms (signaling receptor binding and molecular transducer activity), and pathways (neutrophil degranulation and innate immune system) were obtained.

**Table 3.**
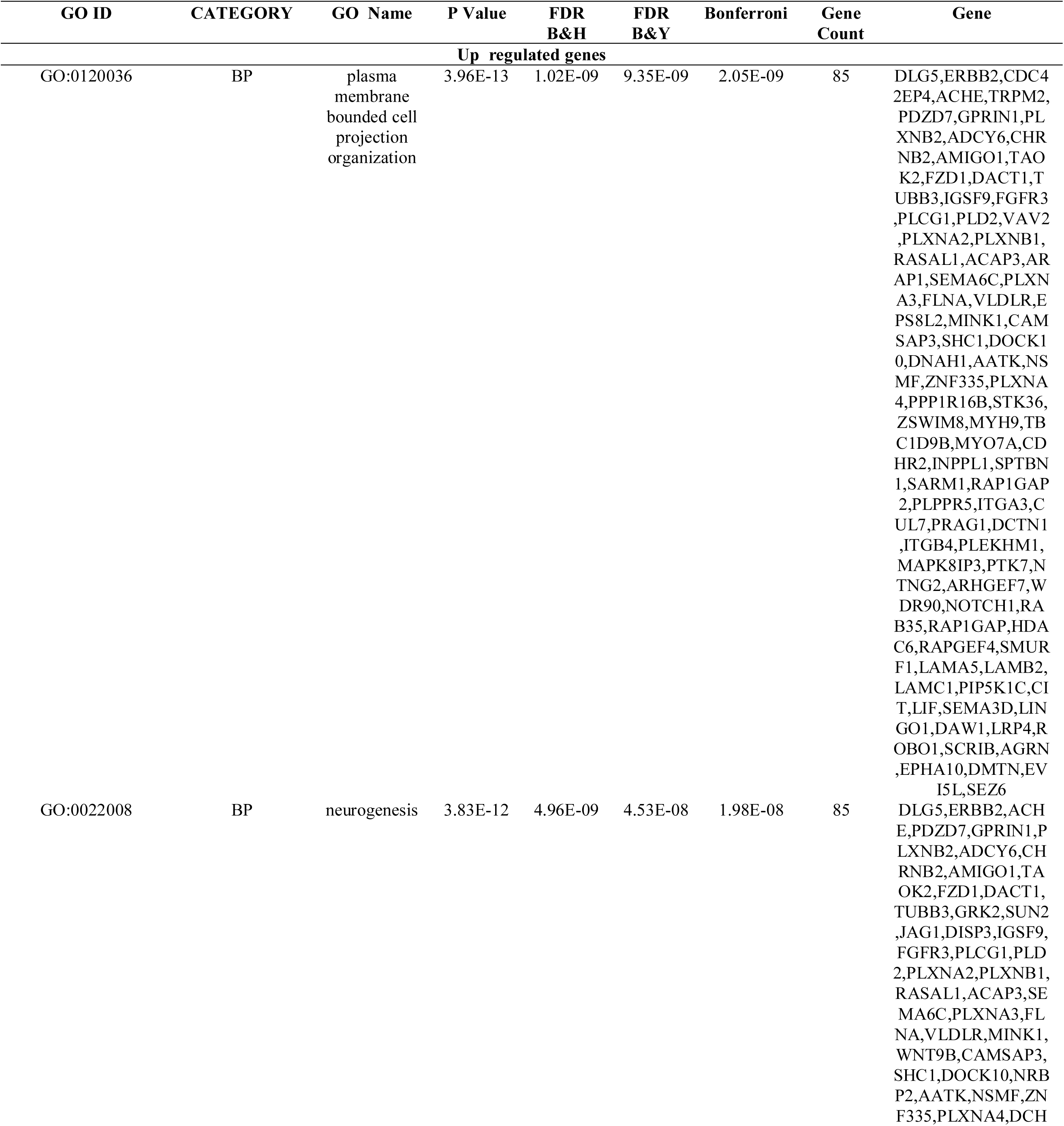
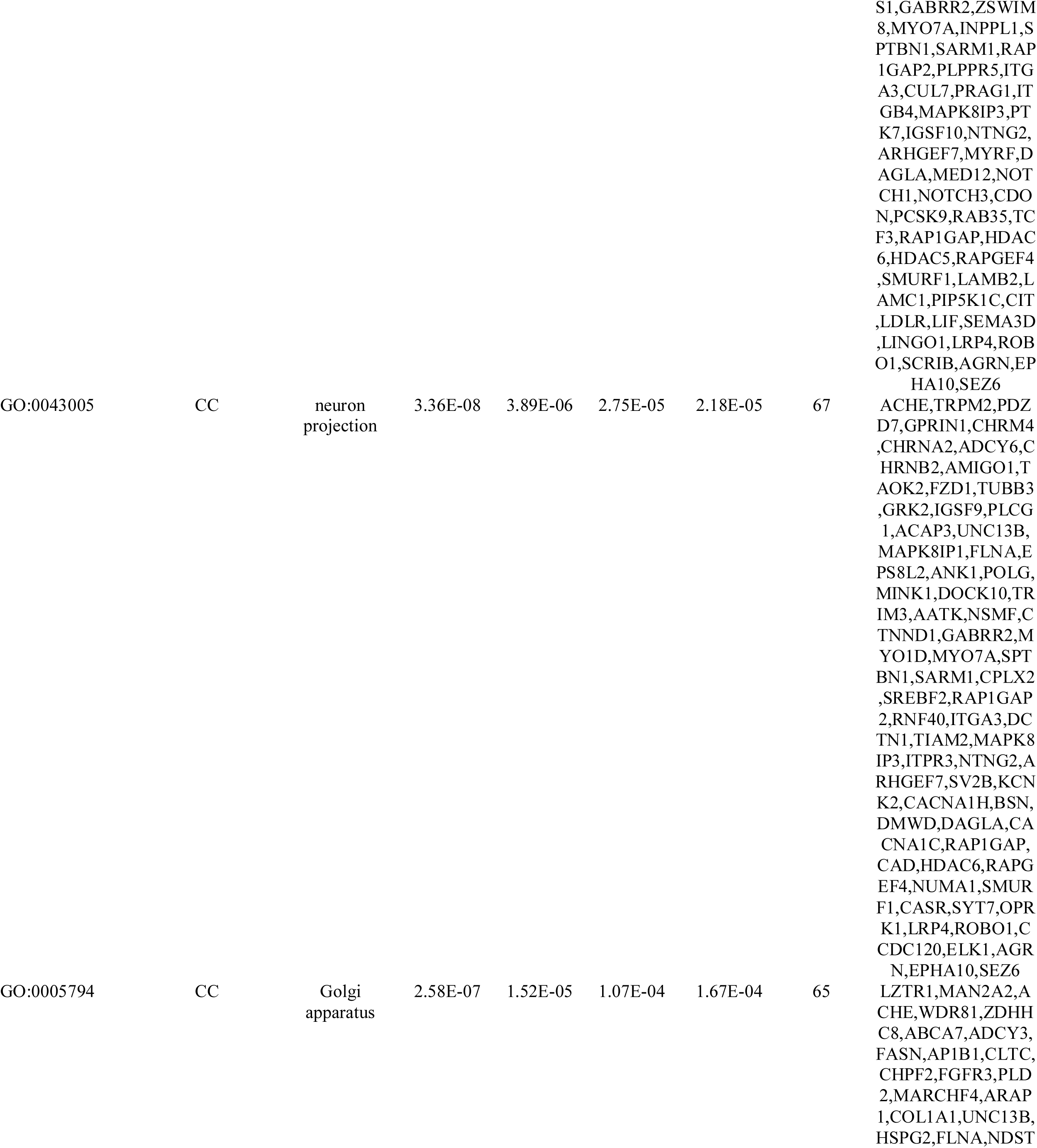
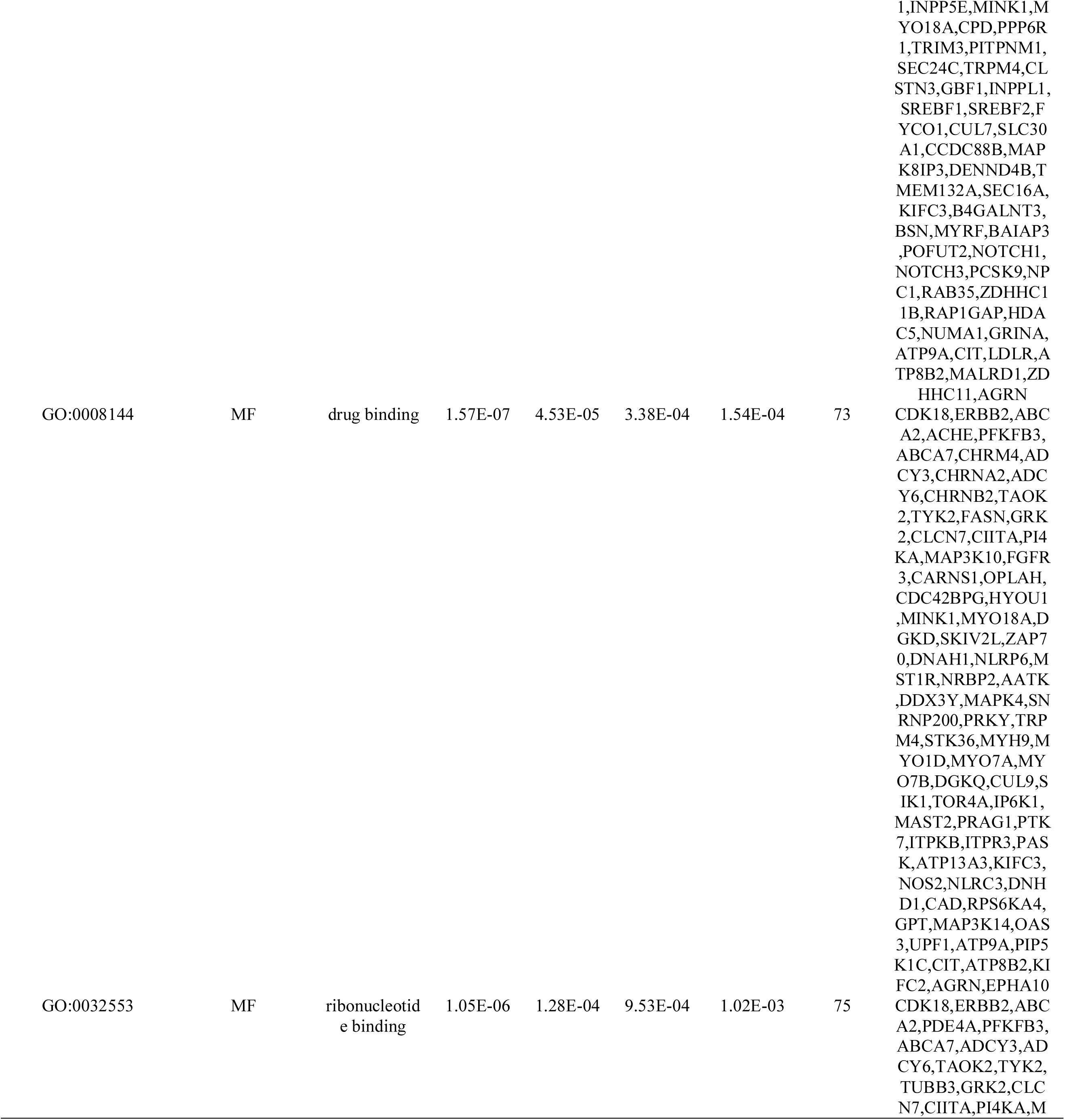
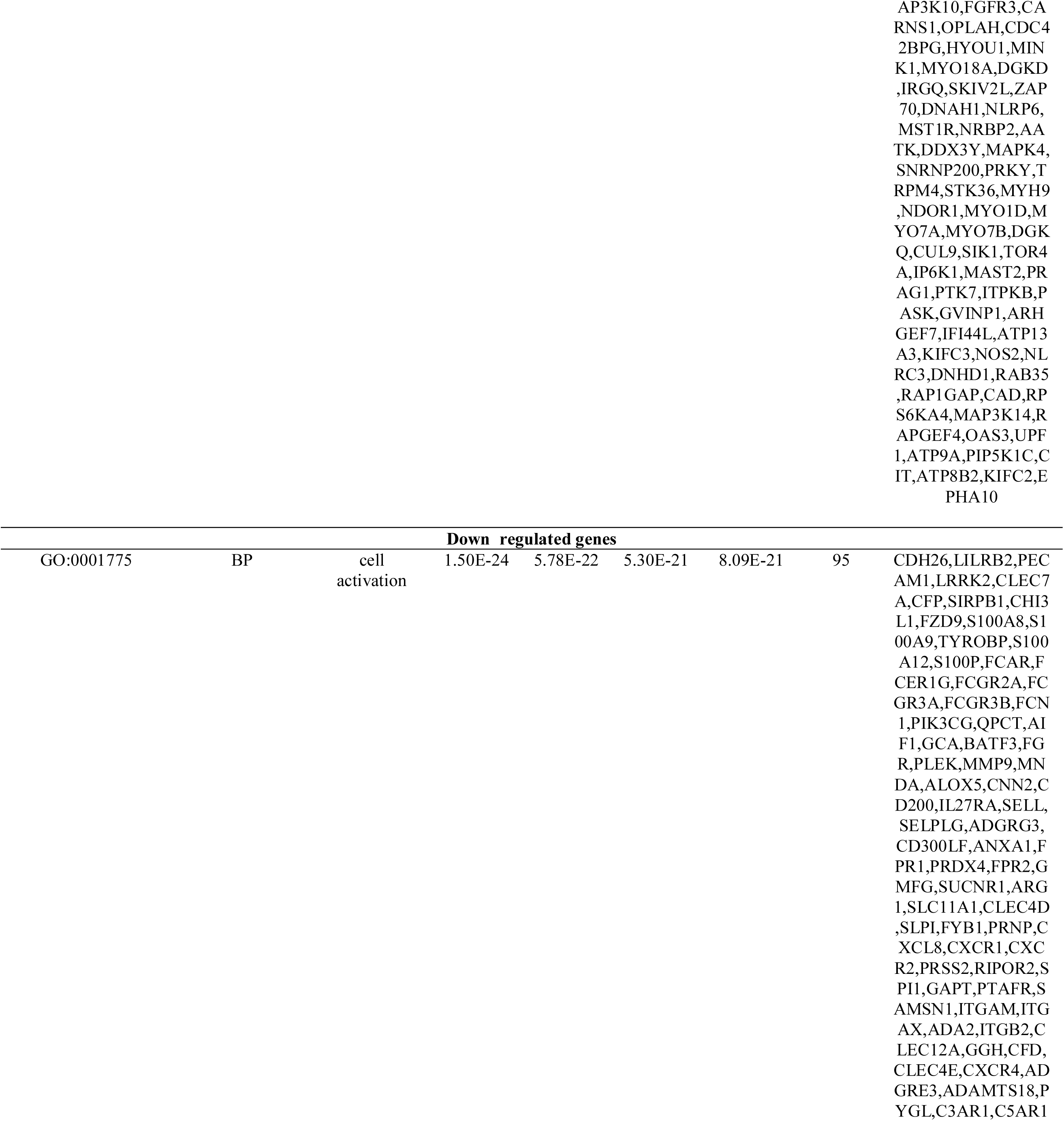
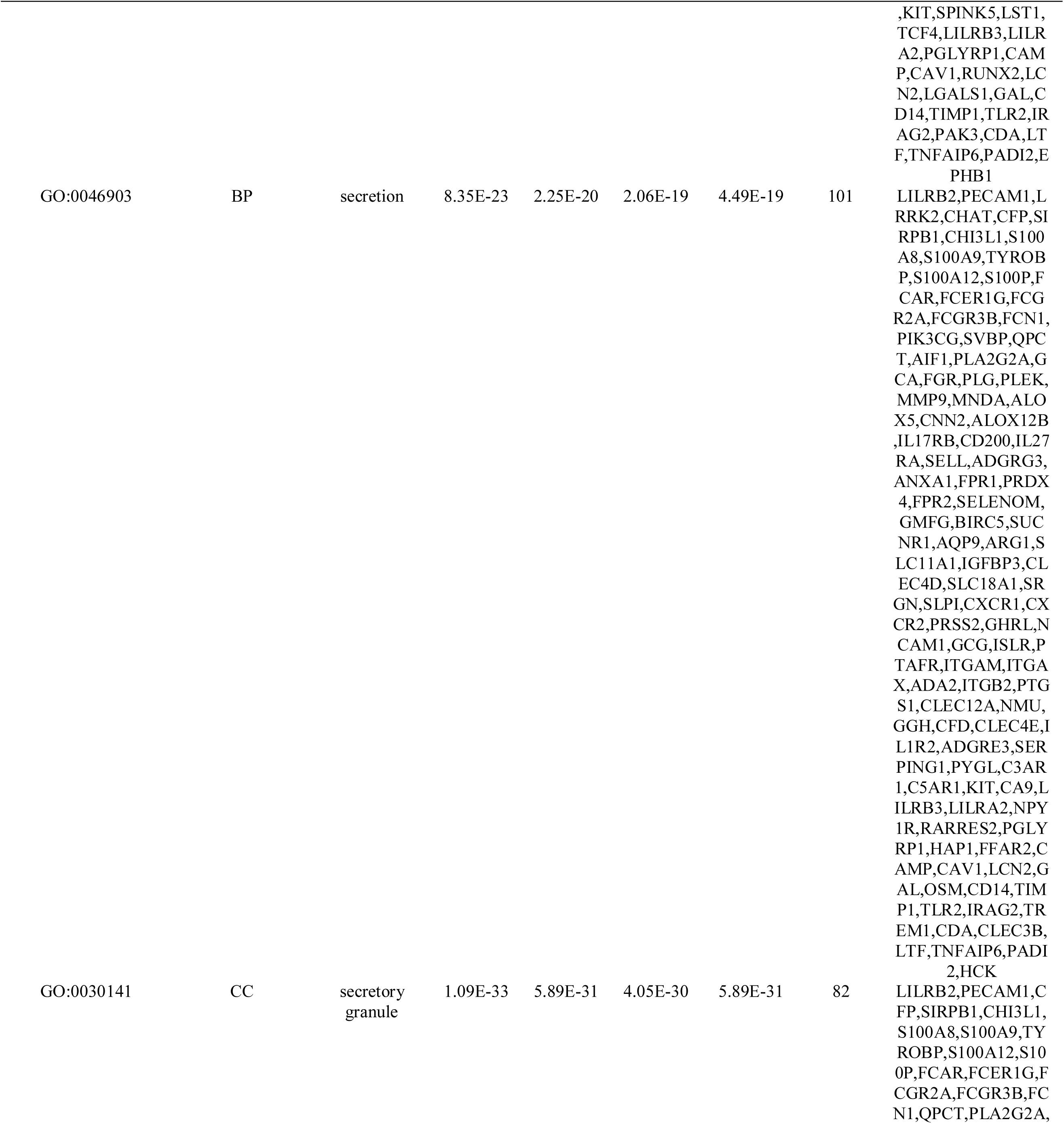
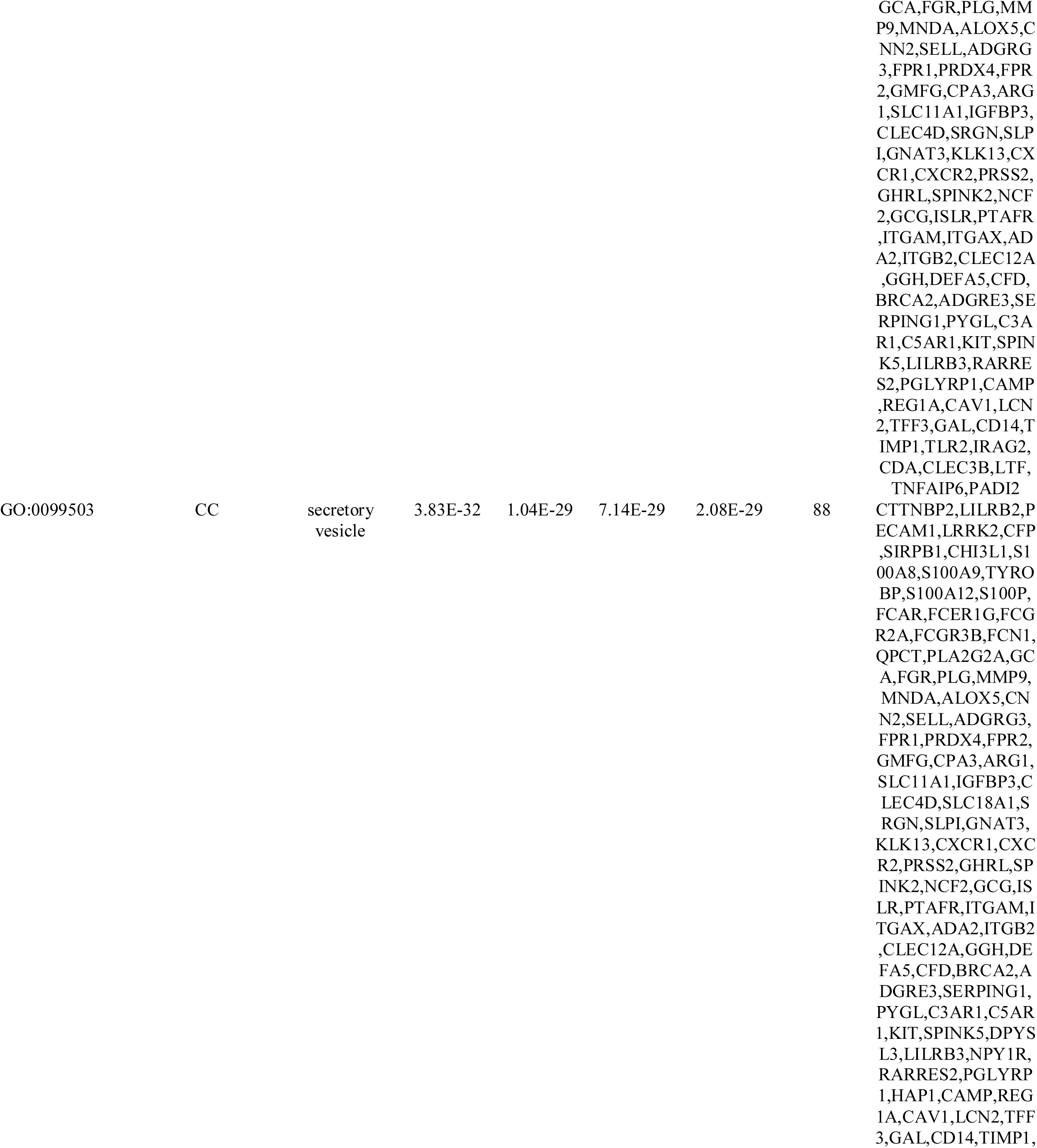
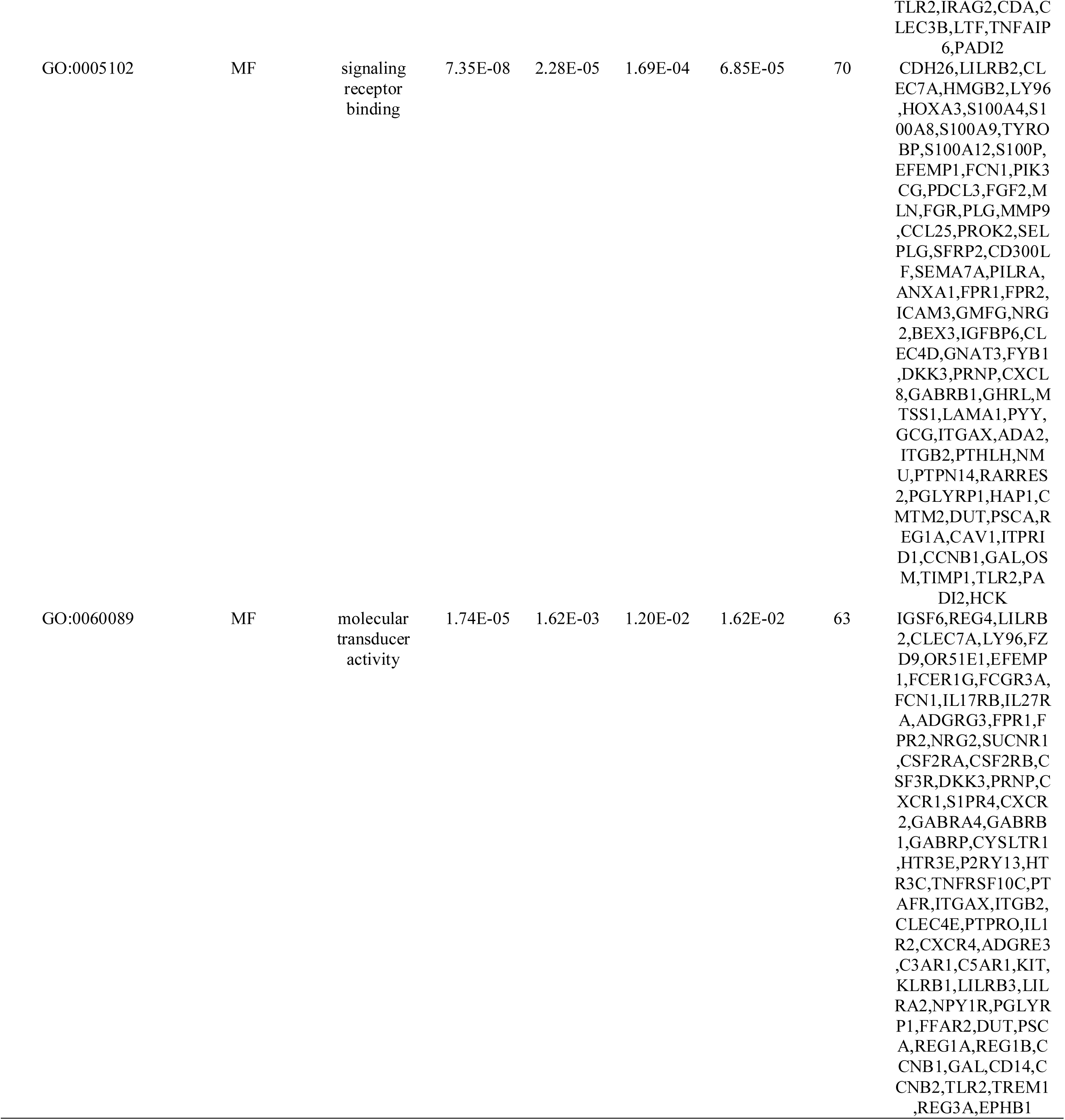
The enriched GO terms of the up and down regulated differentially expressed genes.

**Table 4.**
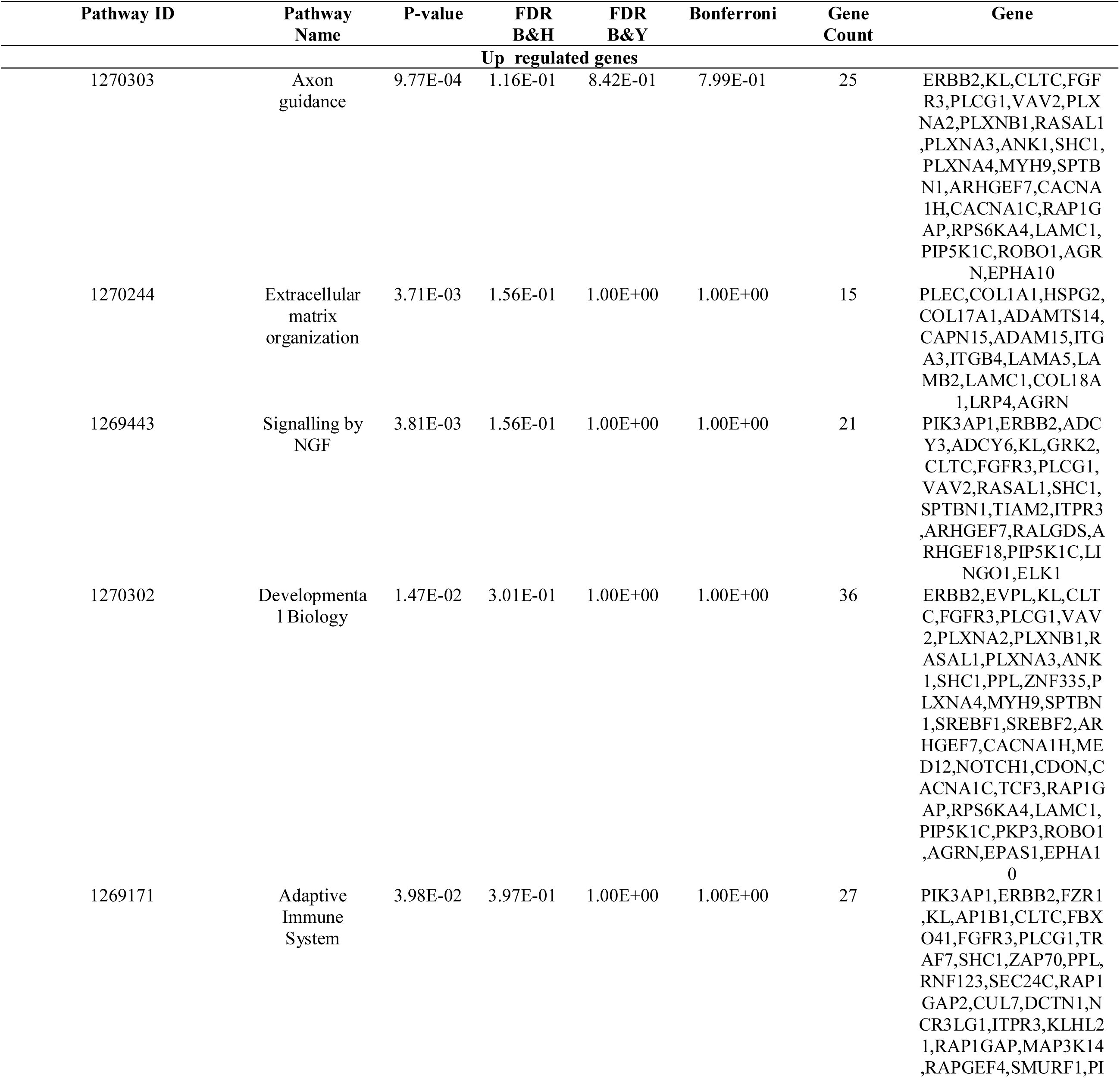
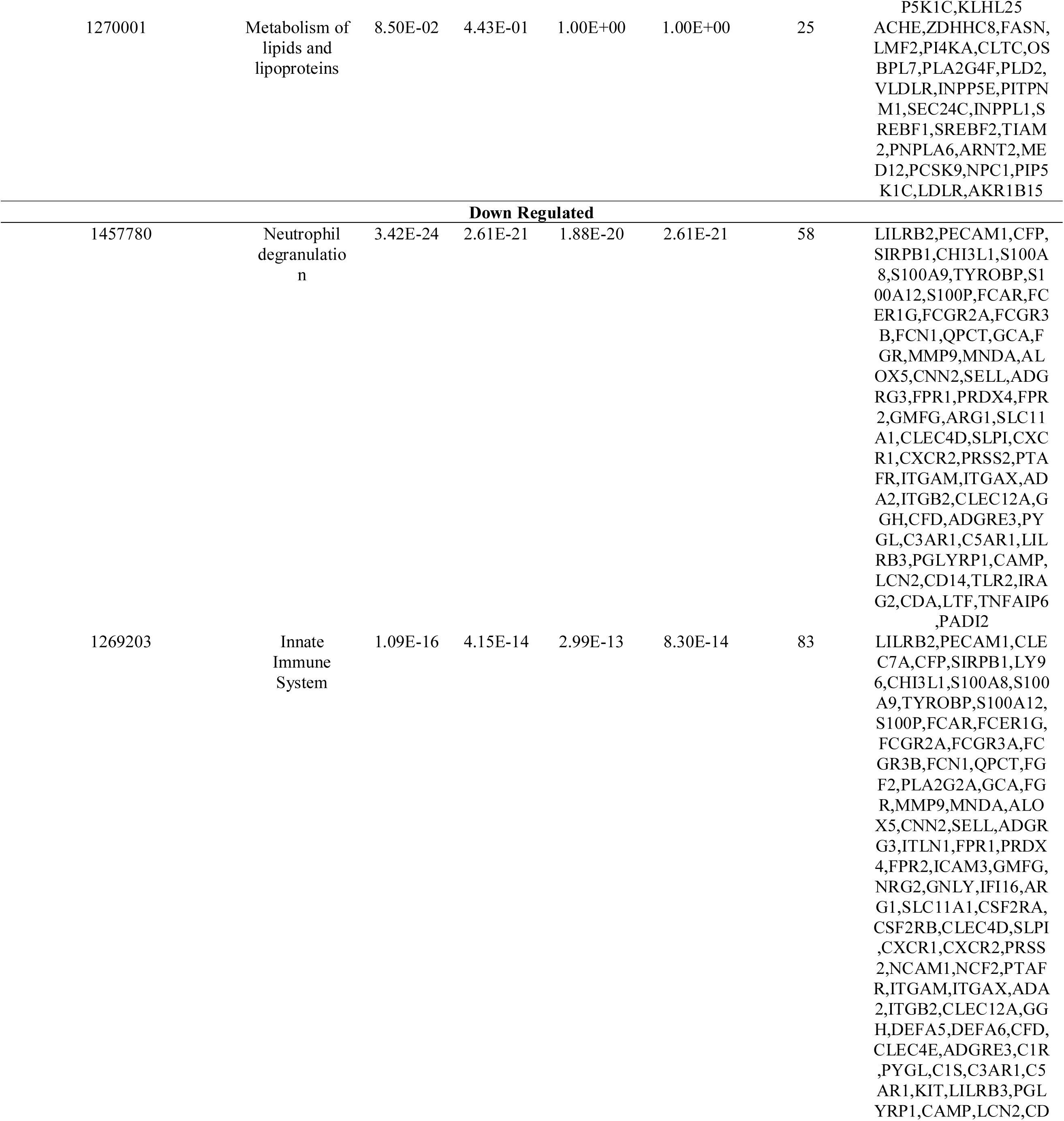
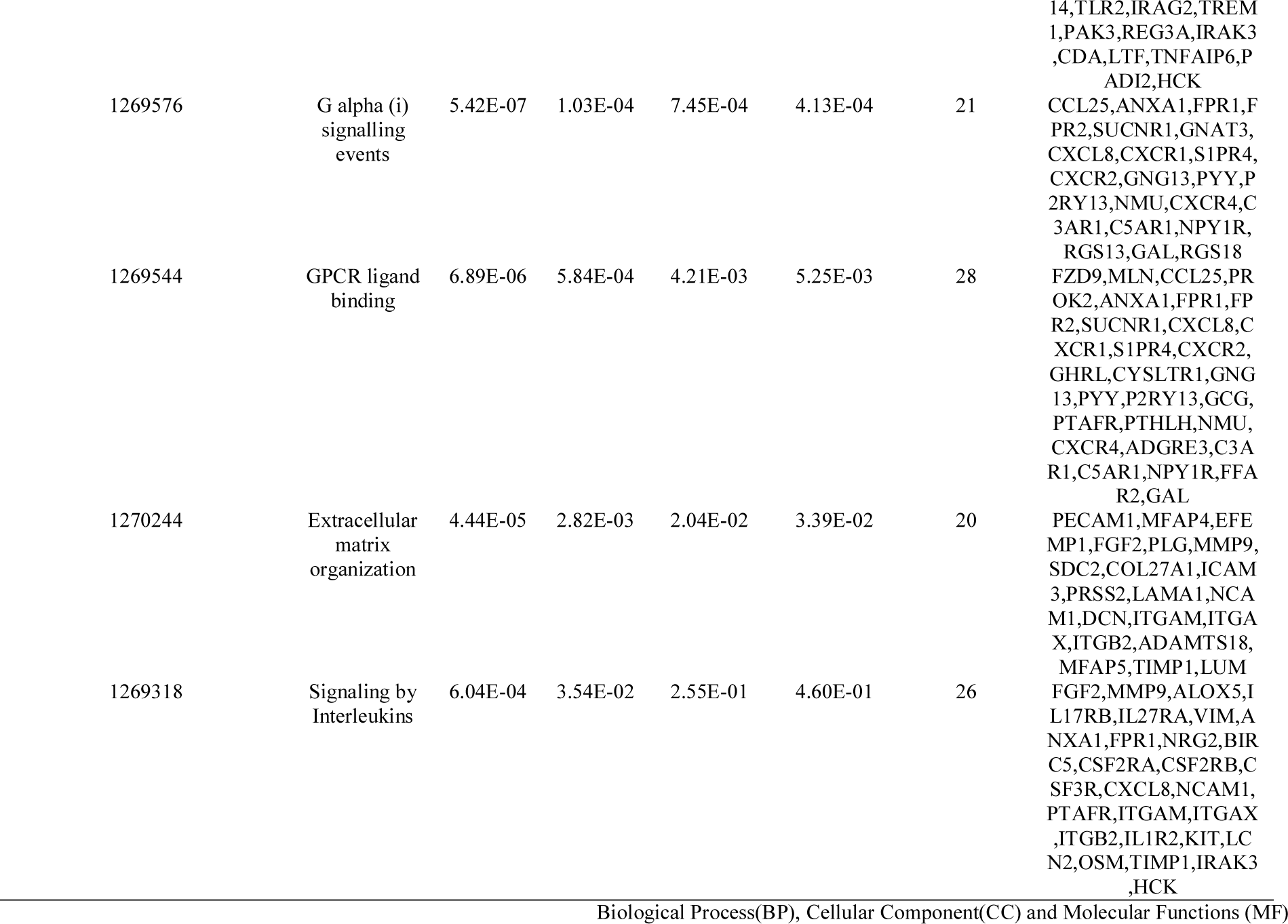
The enriched pathway terms of the up and down regulated differentially expressed genes.

### Protein-protein interaction (PPI) network and module analysis

IID interactome mapped 872 DEGs into PPI network containing 3894 nodes and 7142 edges (Fig. 3A). 10 genes (5 up regulated and 5 down regulated) were identified as hub genes with the high node degree, betweenness centrality, stress centrality and closeness centrality and are listed in Table 5. According to the topological rank, the ten hub genes were MYH9, FLNA, DCTN1, CLTC, ERBB2, TCF4, VIM, LRRK2, IFI16 and CAV1. Hub genes such as MYH9, FLNA, DCTN1, CLTC and ERBB2 up regulated, while TCF4, VIM, LRRK2, IFI16 and CAV1 down regulated. Then PEWCC1 was used to find clusters in the network. Two modules were calculated according to k-core = 2. Among them, module 1 contained 16 nodes and 39 edges (Fig. 3B), module 2 contained 13 nodes and 24 edges (Fig. 3C). These results suggested that the 872 DEGs had effects on obesity associated type 2 diabetes mellitus. In addition, GO terms and pathways were significantly enriched in module 1, including axon guidance, signalling by NGF, plasma membrane bounded cell projection organization and neurogenesis. Meanwhile, pathway was dramatically enriched in module 2, innate immune system.

**Fig. 3.**
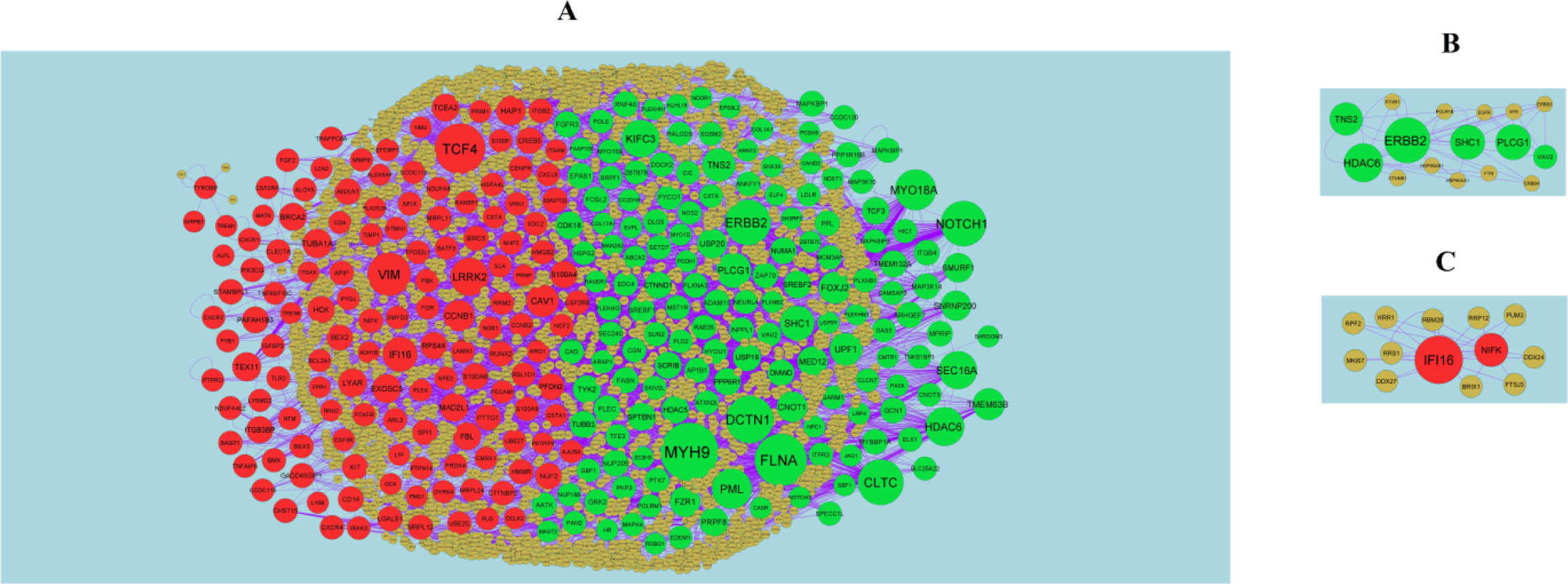
PPI network and the most significant modules of DEGs. (A) The PPI network of DEGs was constructed using Cytoscape (B) The most significant module was obtained from PPI network with 16 nodes and 39 edges for up regulated genes (C) The most significant module was obtained from PPI network with 13 nodes and 24 edges for down regulated genes. Up regulated genes are marked in green; down regulated genes are marked in red

**Table 5.**
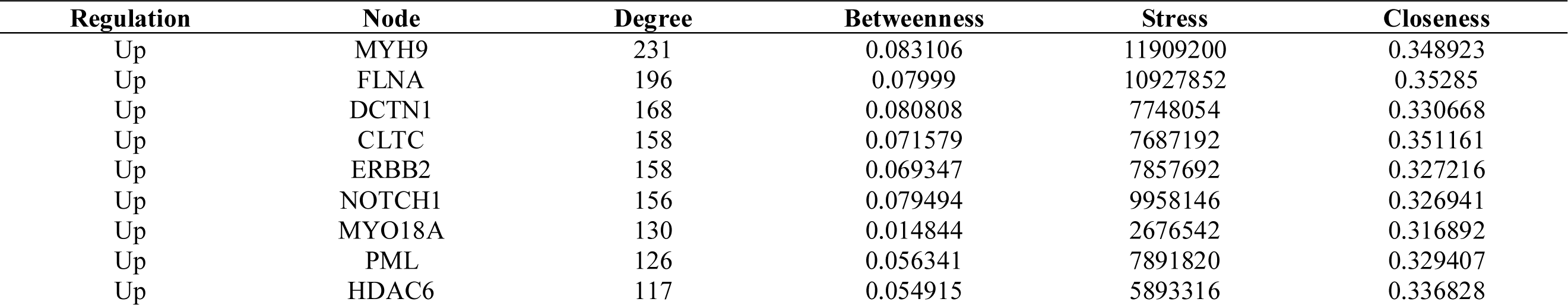
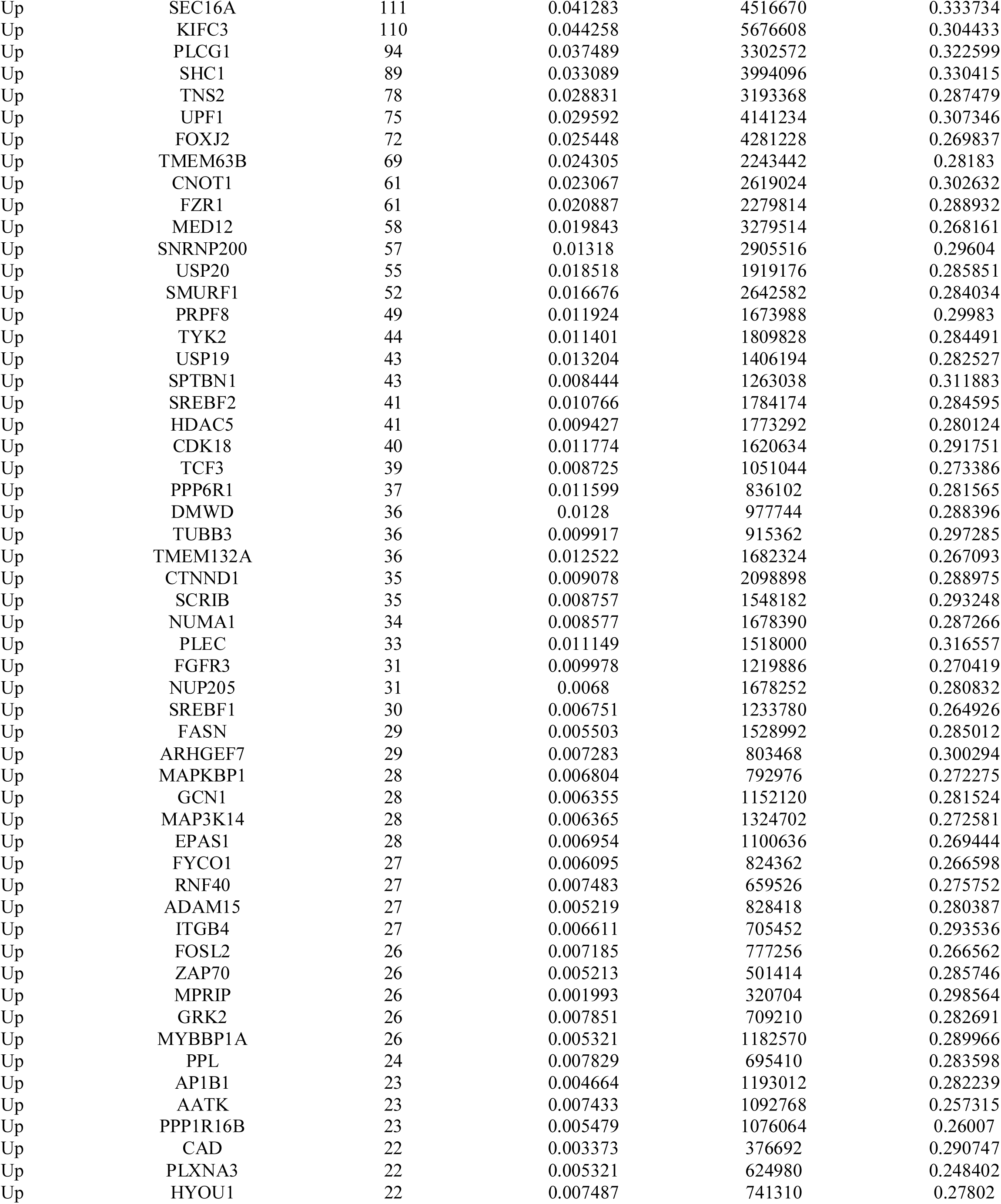
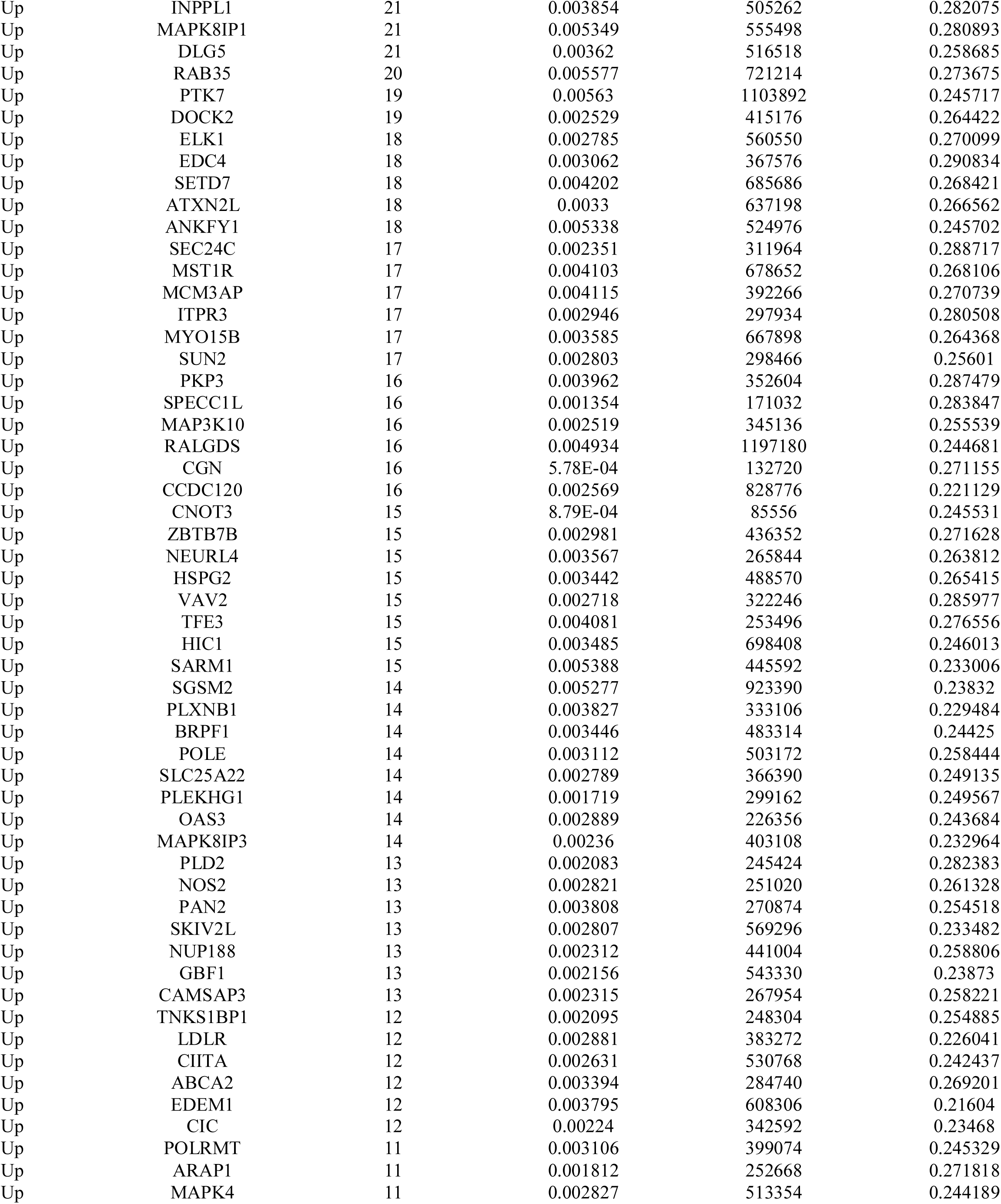
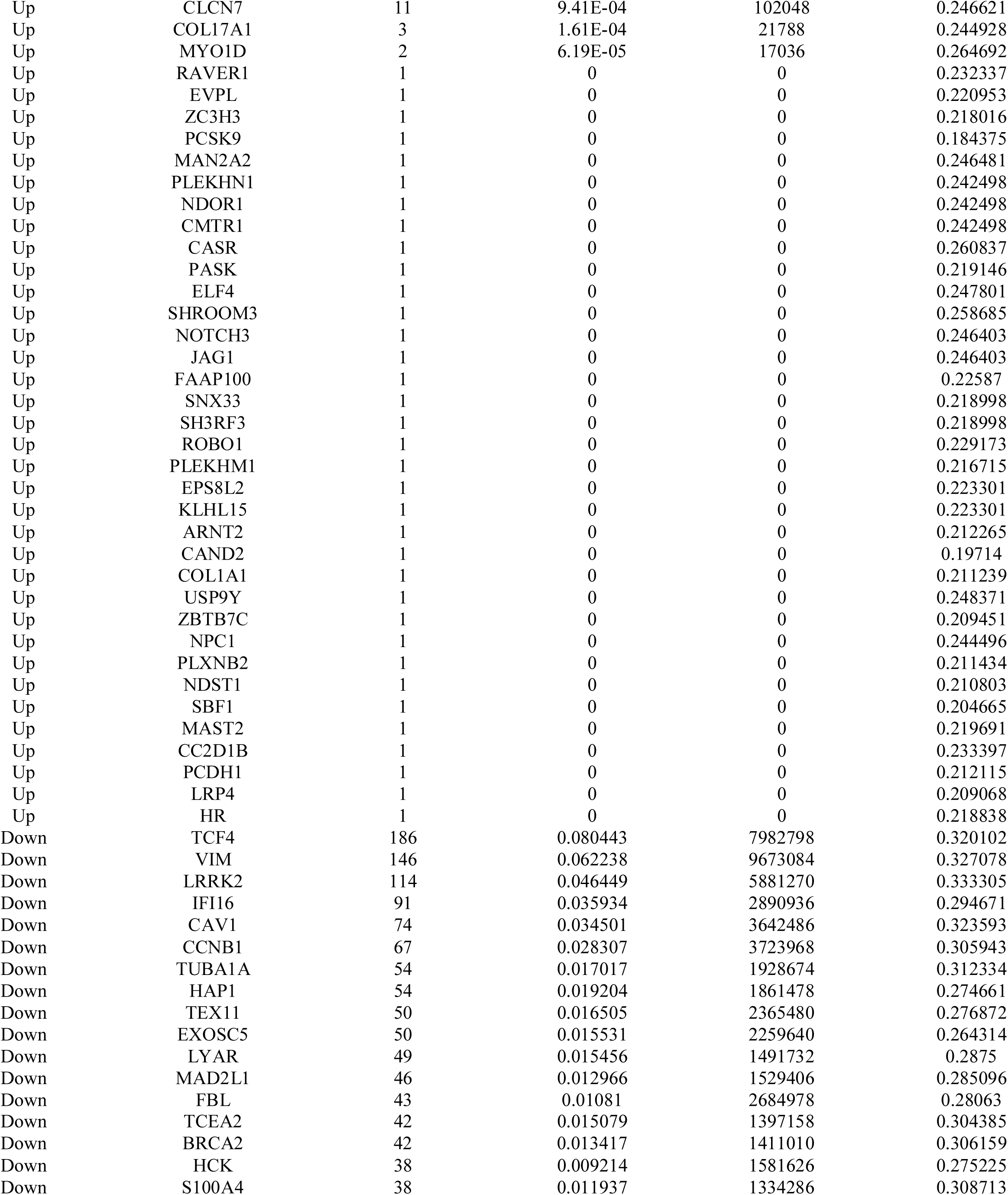
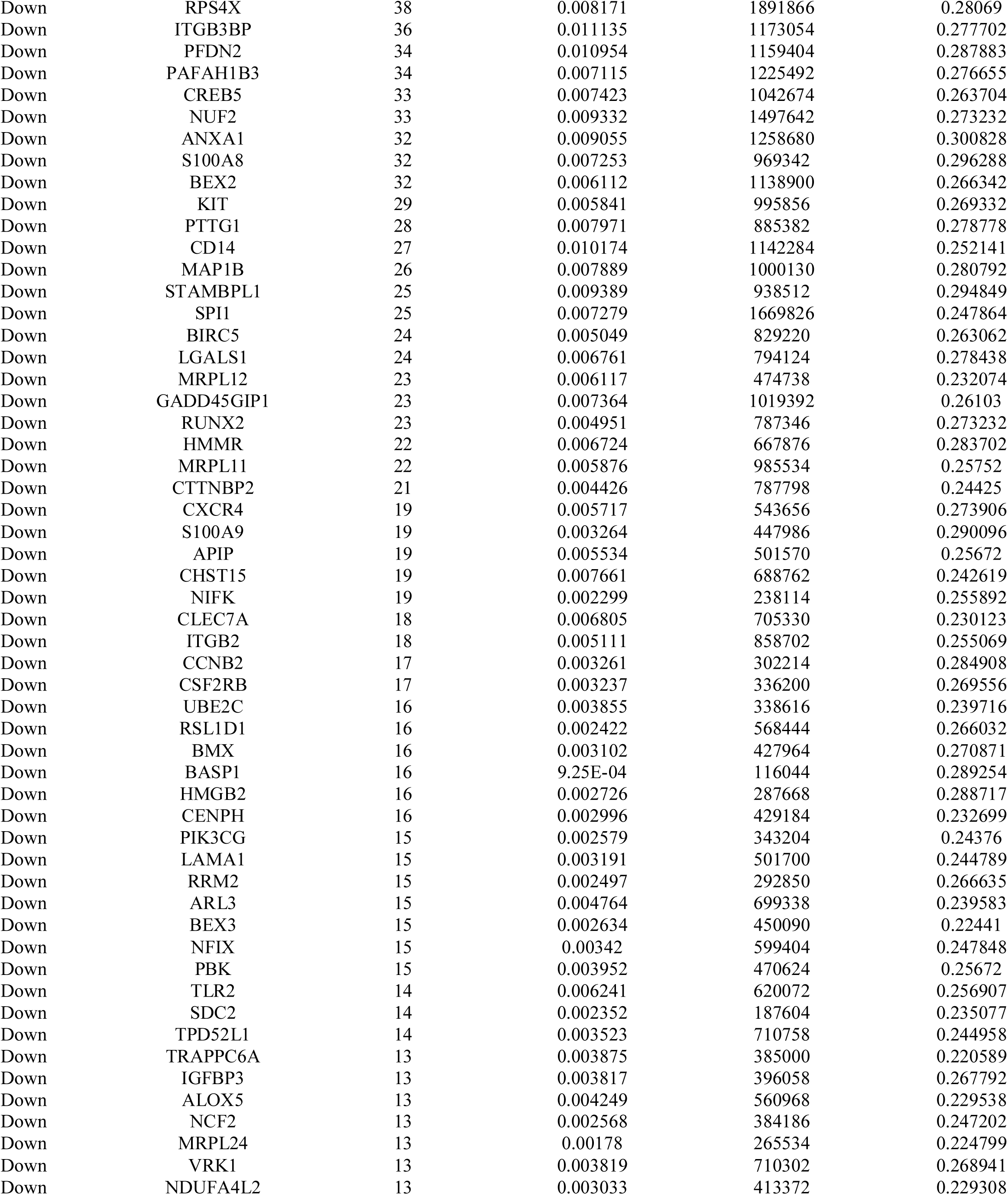
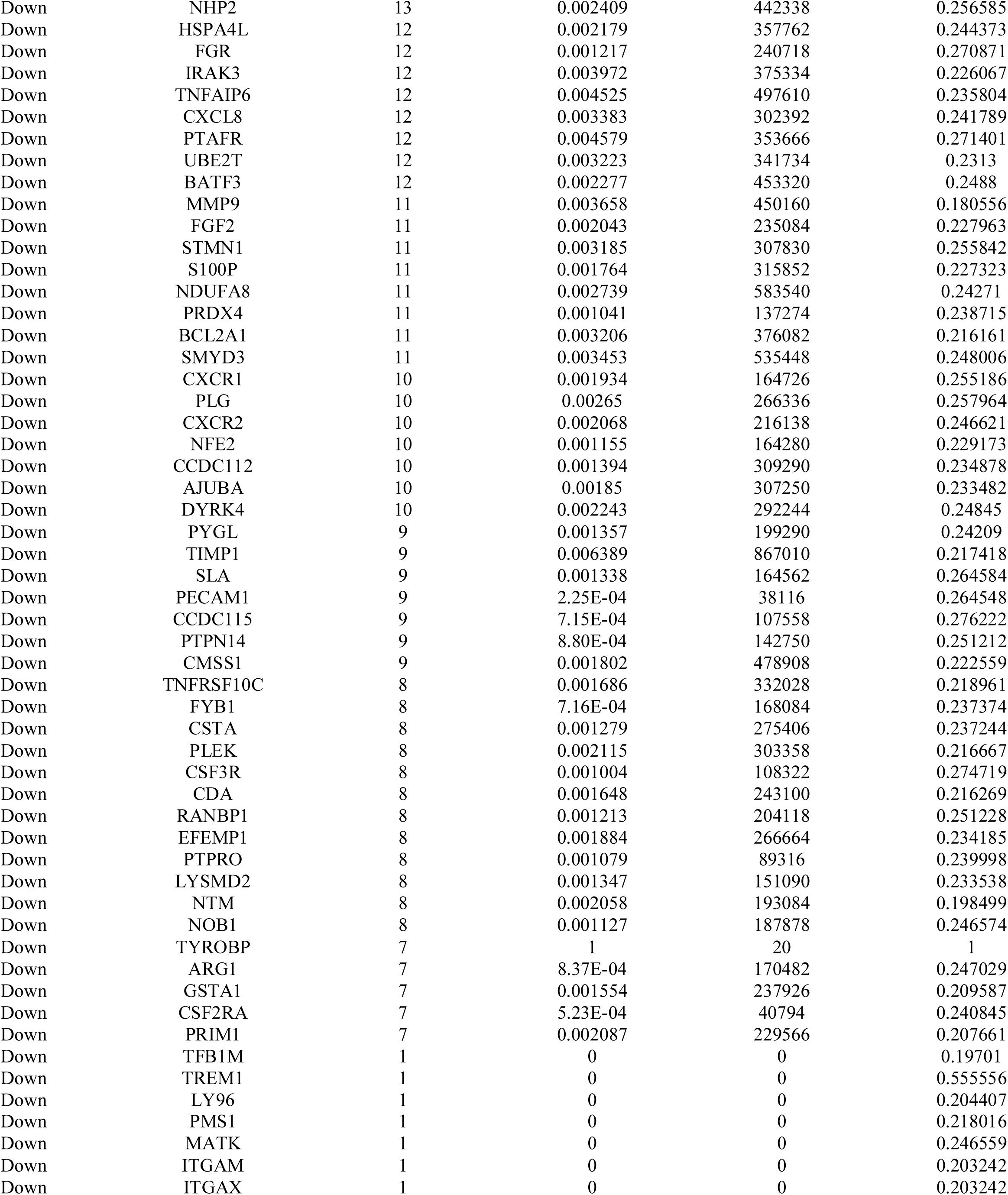
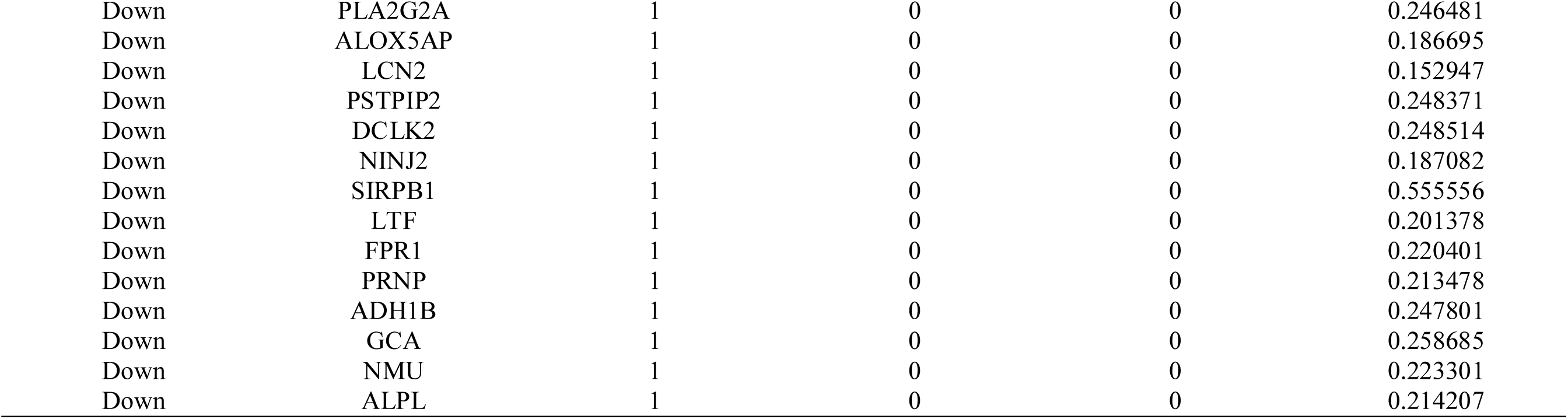
Topology table for up and down regulated genes.

### Target gene - miRNA regulatory network

A total of 296 DEGs, 2224 miRNAs, and 15485 edges were contained in the target gene - miRNA regulatory network (Fig. 4). The nodes with degrees were listed in Table 6. Among them, the degrees of hsa-mir-4329, hsa-mir-4315, hsa-mir-1299, hsa-mir-4779, hsa-mir-3685, hsa-mir-6134, hsa-mir-4459, hsa-mir-6124, hsa-mir-9500 and hsa-mir-1297 were higher in the target gene - miRNA regulatory network. Besides, target genes with higher degrees such as MYH9, ERBB2, MYO18A, SEC16A, PLCG1, CCNB1, CAV1, VIM, HAP1 and MAD2L1.

**Fig. 4.**
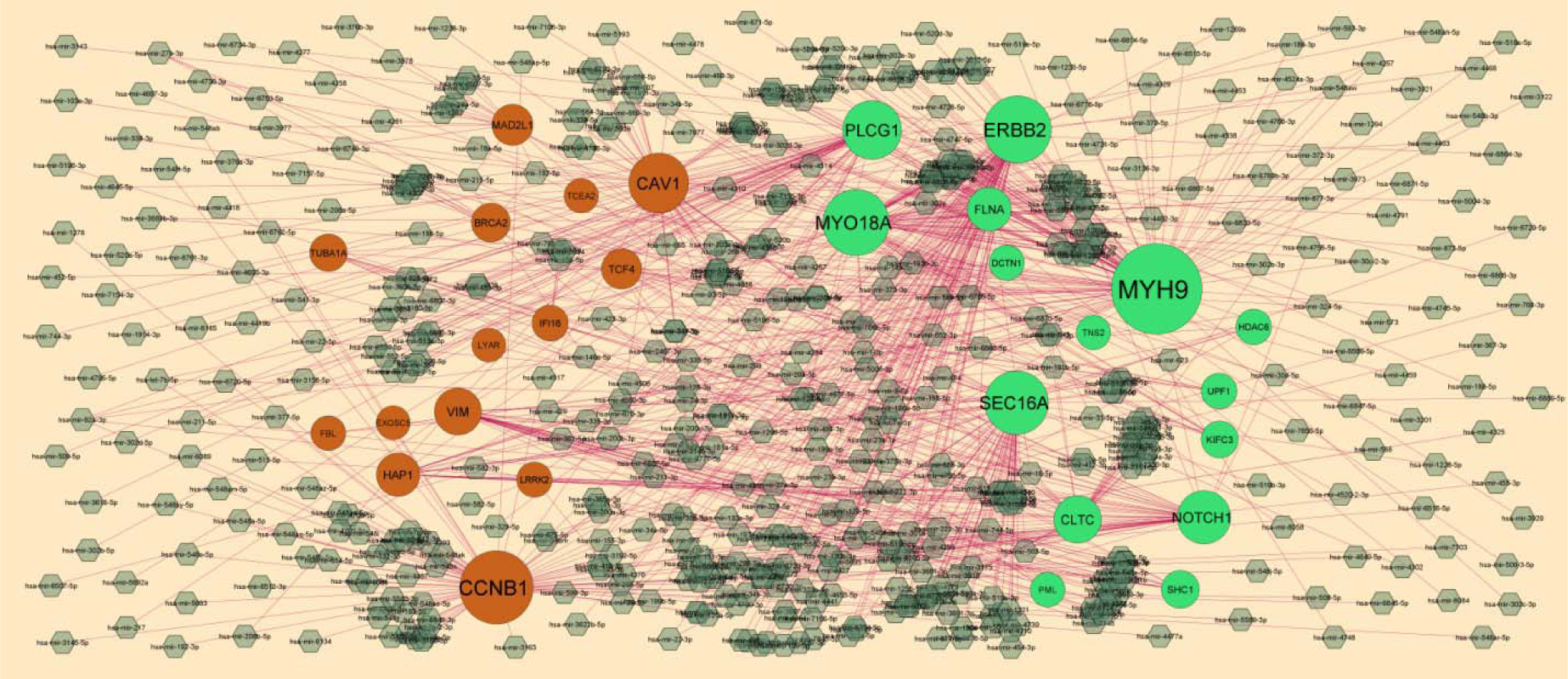
Target gene - miRNA regulatory network between target genes. The black color diamond nodes represent the key miRNAs; up regulated genes are marked in green; down regulated genes are marked in orange.

**Table 6.** miRNA - target gene and TF - target gene interaction.

| Regulation | Target Genes | Degree | MicroRNA | Regulation | Target Genes | Degree | TF |
| --- | --- | --- | --- | --- | --- | --- | --- |
| Up | MYH9 | 116 | hsa-mir-4329 | Up | CLTC | 59 | SMARCA4 |
| Up | ERBB2 | 73 | hsa-mir-4315 | Up | MYH9 | 53 | TCF7 |
| Up | MYO18A | 71 | hsa-mir-1299 | Up | NOTCH1 | 45 | MYB |
| Up | SEC16A | 66 | hsa-mir-4779 | Up | SHC1 | 42 | E2F4 |
| Up | PLCG1 | 56 | hsa-mir-3685 | Up | KIFC3 | 42 | CUX1 |
| Up | NOTCH1 | 44 | hsa-mir-623 | Up | MYO18A | 40 | SALL4 |
| Up | CLTC | 31 | hsa-mir-543 | Up | PML | 32 | CTCF |
| Up | FLNA | 23 | hsa-mir-155-5p | Up | PLCG1 | 29 | SOX17 |
| Up | SHC1 | 10 | hsa-mir-5582-5p | Up | ERBB2 | 24 | MTF2 |
| Up | KIFC3 | 8 | hsa-mir-5193 | Up | UPF1 | 23 | ZFX |
| Up | HDAC6 | 5 | hsa-mir-548m | Up | HDAC6 | 22 | TFAP2A |
| Up | UPF1 | 5 | hsa-mir-4635 | Up | SEC16A | 21 | CEBPB |
| Up | DCTN1 | 4 | hsa-mir-361-5p | Up | FLNA | 18 | SETDB1 |
| Up | PML | 3 | hsa-mir-378a-3p | Up | DCTN1 | 15 | RBPJ |
| Up | TNS2 | 2 | hsa-mir-26b-5p | Down | TCF4 | 50 | NANOG |
| Down | CCNB1 | 84 | hsa-mir-6134 | Down | VIM | 43 | GFI1B |
| Down | CAV1 | 58 | hsa-mir-4459 | Down | CAV1 | 36 | GATA4 |
| Down | VIM | 30 | hsa-mir-6124 | Down | FBL | 33 | HIF1A |
| Down | HAP1 | 22 | hsa-mir-9500 | Down | TUBA1A | 32 | CLOCK |
| Down | MAD2L1 | 17 | hsa-mir-1297 | Down | EXOSC5 | 29 | FLI1 |
| Down | TCF4 | 15 | hsa-mir-203a-3p | Down | MAD2L1 | 27 | AP1S2 |
| Down | BRCA2 | 12 | hsa-mir-1245a | Down | TCEA2 | 26 | MYCN |
| Down | TUBA1A | 9 | hsa-mir-744-5p | Down | BRCA2 | 26 | E2F1 |
| Down | IFI16 | 6 | hsa-mir-136-5p | Down | LYAR | 22 | AR |
| Down | LRRK2 | 4 | hsa-mir-582-3p | Down | IFI16 | 19 | REST |
| Down | FBL | 3 | hsa-mir-181b-5p | Down | CCNB1 | 18 | KLF4 |
| Down | TCEA2 | 2 | hsa-mir-17-5p | Down | LRRK2 | 16 | SUZ12 |
| Down | LYAR | 1 | hsa-mir-16-5p | Down | HAP1 | 15 | RUNX1 |
| Down | EXOSC5 | 1 | hsa-mir-31-5p | Down | TEX11 | 6 | SMAD4 |

**Table 7.** Docking results of Designed Molecules on Over expressed Proteins.

| Sl. No/<br>Code | CLTC (Clathrin heavy chain) | | | DCTN1 (Dynactin subunit 1) | | | ERBB2 (Erb-β2 receptor tyrosine kinase 2) | | | FLNA (Filamin A) | | | MYH9 (Myosin heavy chain 9) | | |
| --- | --- | --- | --- | --- | --- | --- | --- | --- | --- | --- | --- | --- | --- | --- | --- |
|  | PDB: 5M5R |  |  | PDB: 2HL3 |  |  | PDB: 1MFG |  |  | PDB: 3HOC |  |  | PDB: 4ZS4 |  |  |
|  | Total Score | Crash (-Ve) | Polar | Total Score | Crash (-Ve) | Polar | Total Score | Crash (-Ve) | Polar | Total Score | Crash (-Ve) | Polar | Total Score | Crash (-Ve) | Polar |
| FLA | 4.337 | -0.196 | 1.130 | 2.896 | -0.195 | 2.236 | 3.347 | -1.149 | 0.336 | 3.181 | -0.321 | 1.146 | 5.331 | -0.340 | 2.985 |
| FLS | 5.299 | -0.700 | 3.033 | 3.960 | -0.836 | 3.690 | 3.638 | -0.591 | 1.527 | 3.216 | -0.648 | 1.146 | 4.862 | -0.588 | 2.112 |
| FLO | 5.527 | -1.348 | 3.910 | 4.340 | -0.619 | 4.061 | 3.638 | -0.591 | 1.527 | 4.143 | -0.465 | 2.139 | 4.502 | -0.193 | 1.656 |
| FAS | 5.304 | -0.569 | 2.387 | 3.969 | -0.842 | 2.733 | 3.477 | -1.106 | 0 | 3.241 | -0.779 | 2.059 | 4.657 | -1.529 | 1.457 |
| ANC | 6.485 | -1.200 | 5.319 | 5.082 | -1.036 | 1.942 | 3.915 | -1.639 | 3.305 | 3.588 | -0.821 | 3.532 | 5.762 | -1.013 | 3.925 |
| ERI | 4.863 | -0.737 | 2.806 | 3.650 | -1.287 | 0.899 | 4.151 | -0.472 | 2.876 | 4.476 | -0.996 | 5.665 | 5.935 | -0.562 | 4.321 |
| ISO | 3.808 | -0.576 | 0.074 | 2.759 | -0.212 | 3.222 | 3.959 | -1.169 | 0.815 | 2.266 | -0.486 | 0 | 4.080 | -0.479 | 2.012 |
| QRC | 6.685 | -1.572 | 5.384 | 5.272 | -0.706 | 1.763 | 5.728 | -1.046 | 4.023 | 4.472 | -1.082 | 4.375 | 5.233 | -0.985 | 3.188 |
| FIS | 5.510 | -1.515 | 4.728 | 4.332 | -0.444 | 3.662 | 4.245 | -0.481 | 3.342 | 4.057 | -0.705 | 5.309 | 5.347 | -0.485 | 3.754 |
| MOR | 6.142 | -1.413 | 5.757 | 4.818 | -0.548 | 5.218 | 4.932 | -0.839 | 2.943 | 5.520 | -0.903 | 6.341 | 6.566 | -0.540 | 4.981 |
| NAR | 5.588 | -0.487 | 2.548 | 4.341 | -0.259 | 3.278 | 4.584 | -0.347 | 2.910 | 4.615 | -0.586 | 5.288 | 5.159 | -0.457 | 2.929 |
| DIO | 10.047 | -2.772 | 7.675 | 6.490 | -2.201 | 7.258 | 7.457 | -1.358 | 6.205 | 7.250 | -1.807 | 7.062 | 8.947 | -3.961 | 6.441 |
| CHR | 5.247 | -1.848 | 3.680 | 3.932 | -0.419 | 4.022 | 4.097 | -0.431 | 1.830 | 3.553 | -0.454 | 2.594 | 4.598 | -0.888 | 1.588 |
| HES | 10.811 | -2.994 | 10.042 | 7.400 | -1.292 | 8.085 | 5.331 | -2.175 | 2.727 | 6.591 | -1.921 | 6.877 | 9.459 | -3.331 | 7.279 |
| CYA | 5.950 | -0.917 | 5.305 | 4.778 | -0.441 | 5.109 | 4.449 | -0.881 | 4.118 | 5.300 | -1.199 | 5.973 | 5.012 | -0.973 | 3.818 |
| API | 4.863 | -0.737 | 2.806 | 3.345 | -0.403 | 2.756 | 3.795 | -0.951 | 2.092 | 4.410 | -0.662 | 5.361 | 4.816 | -0.688 | 2.395 |
| BAI | 4.680 | -0.575 | 2.886 | 3.252 | -1.720 | 1.810 | 3.926 | -0.655 | 1.980 | 3.668 | -0.512 | 4.063 | 4.671 | -0.518 | 3.621 |
| LUT | 5.044 | -1.731 | 4.513 | 3.667 | -0.456 | 3.878 | 4.440 | -0.671 | 2.932 | 4.444 | -0.684 | 4.383 | 5.480 | -0.805 | 3.817 |
| KAE | 5.459 | -0.512 | 3.978 | 4.098 | -0.325 | 4.067 | 4.597 | -1.132 | 1.831 | 5.172 | -0.759 | 6.381 | 6.168 | -0.389 | 4.820 |
| PEL | 6.143 | -0.711 | 5.091 | 4.967 | -0.837 | 2.047 | 2.806 | -0.285 | 2.912 | 3.914 | -0.881 | 3.213 | 4.889 | -0.888 | 2.343 |
| TAN | 5.313 | -0.780 | 2.292 | 3.976 | -0.570 | 4.999 | 5.845 | -1.719 | 1.923 | 3.822 | -0.607 | 2.551 | 6.069 | -0.771 | 2.843 |
| WOG | 5.103 | -1.196 | 3.749 | 3.929 | -0.426 | 4.091 | 5.889 | -0.926 | 2.762 | 4.435 | -0.935 | 2.793 | 5.947 | -1.526 | 4.157 |
| ISH | 5.605 | -1.025 | 3.033 | 4.488 | -0.986 | 0.051 | 5.490 | -1.485 | 4.304 | 5.090 | -0.828 | 3.994 | 5.841 | -1.464 | 4.203 |
| DAI | 4.690 | -0.951 | 2.151 | 3.283 | -0.317 | 1.941 | 4.456 | -1.703 | 1.907 | 2.918 | -0.500 | 2.677 | 5.128 | -0.841 | 2.251 |
| GEN | 4.538 | -0.232 | 4.426 | 2.944 | -0.608 | 0.526 | 4.529 | -1.124 | 1.344 | 4.028 | -0.554 | 3.053 | 5.676 | -0.780 | 3.799 |
| RUT | 10.657 | -1.576 | 11.201 | 6.716 | -1.292 | 2.821 | 6.147 | -2.850 | 4.021 | 6.744 | -2.346 | 8.022 | 9.432 | -3.408 | 7.926 |
| DEL | 5.942 | -1.187 | 4.706 | 4.501 | -0.490 | 4.425 | 3.903 | -0.621 | 3.159 | 5.151 | -1.447 | 5.207 | 5.208 | -2.253 | 3.572 |
| PHL | 4.162 | -0.959 | 4.484 | 2.799 | -0.364 | 2.369 | 5.199 | -0.378 | 3.220 | 3.491 | -0.665 | 4.755 | 3.173 | -0.966 | 2.568 |

### Target gene - TF regulatory network

A total of 292 DEGs, 195 TFs, and 7094 edges were contained in the target gene - TF regulatory network (Fig. 5). The nodes with degrees were listed in Table 6. Among them, the degrees of SMARCA4, TCF7, MYB, E2F4, CUX1, NANOG, GFI1B, GATA4, HIF1A and CLOCK were higher in the target gene - TF regulatory network. Besides, target genes with higher degrees such as CLTC, MYH9, NOTCH1, SHC1, KIFC3, TCF4, VIM, CAV1, FBL and TUBA1A.

**Fig. 5.**
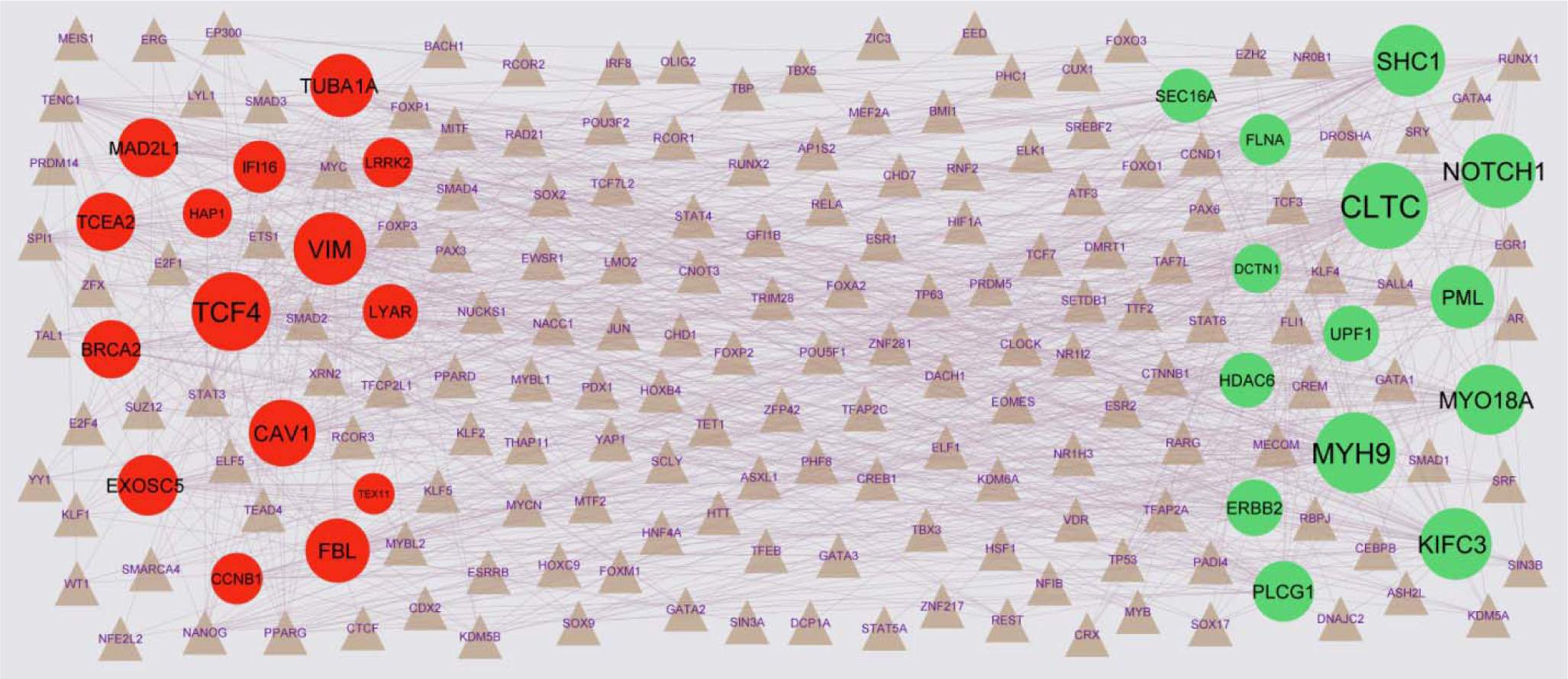
Target gene - TF regulatory network between target genes. The gray color triangle nodes represent the key TFs; up regulated genes are marked in green; down regulated genes are marked in red.

### Receiver operating characteristic (ROC) analysis

ROC curve analysis was performed and the AUC calculated to assess the discriminatory ability of hub genes (Fig. 6). Validated by ROC curves, we found that hub genes had high sensitivity and specificity, including MYH9 (AUC = 0.877), FLNA (AUC = 0.796), DCTN1 (AUC = 0.773), CLTC (AUC = 0.791), ERBB2 (AUC = 0.828), TCF4 (AUC = 0.764), VIM (AUC = 0.838), LRRK2 (AUC = 0.767), IFI16 (AUC = 0.816) and CAV1 (AUC = 0.749). The three genes may be biomarkers of obesity associated type 2 diabetes mellitus and have positive implications for early medical intervention of the disease.

**Fig. 6.**
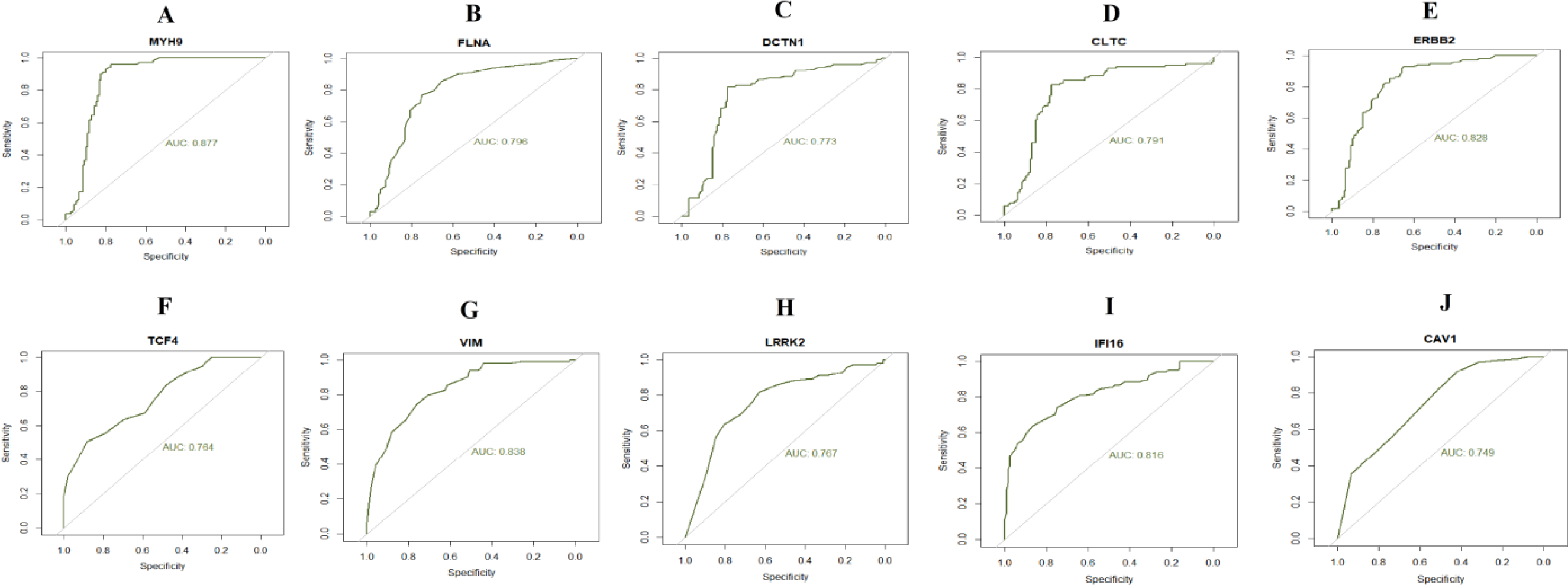
ROC curve analyses of hub genes. A) MYH9 B) FLNA C) DCTN1 D) CLTC E) ERBB2 F) TCF4 G) VIM H) LRRK2 I) IFI16 J) CAV1

### Validation of the expression levels of candidate genes by RT-PCR

MYH9, FLNA, DCTN1, CLTC, ERBB2, TCF4, VIM, LRRK2, IFI16 and CAV1 were selected to validate the expression of integrated analysis. The RT-PCR results showed that MYH9, FLNA, DCTN1, CLTC and ERBB2 were up regulated, which were consistent with that of integrated analysis. While TCF4, VIM, LRRK2, IFI16 and CAV1 were down regulated, these were consistent with that of integrated analysis. The result of RT - PCR was shown in Fig. 7.

**Fig. 7.**
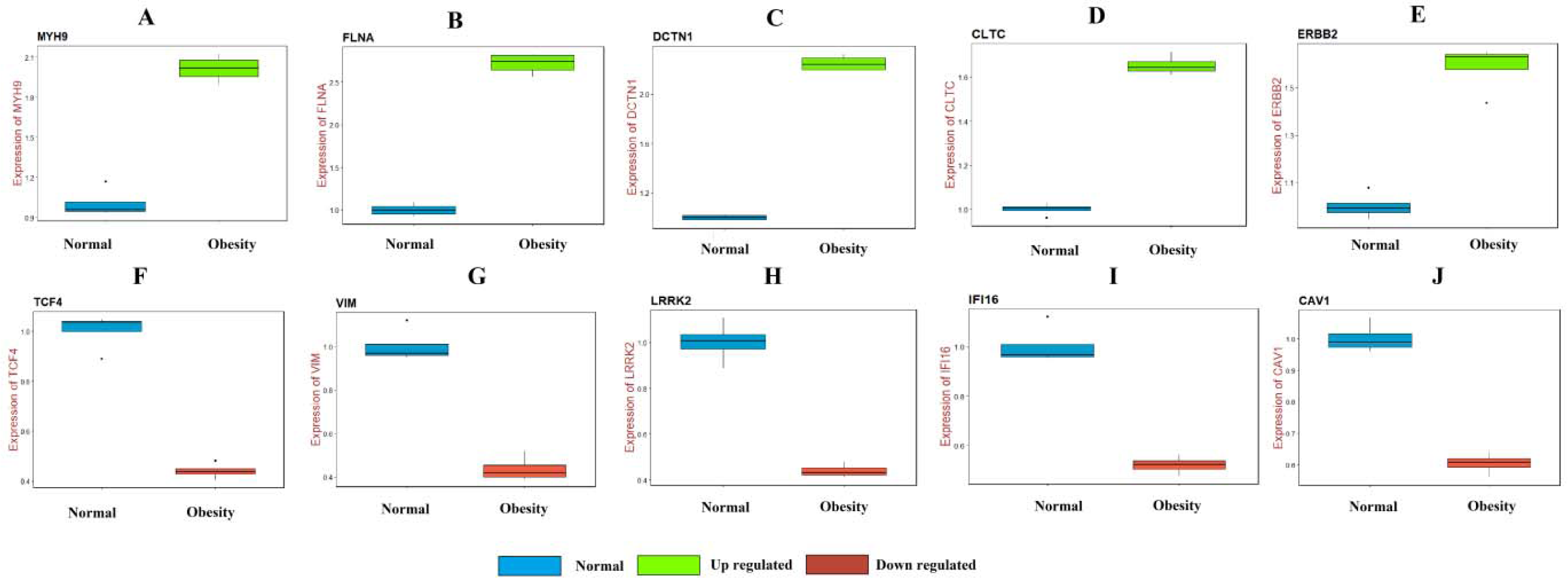
RT-PCR analyses of hub genes. A) MYH9 B) FLNA C) DCTN1 D) CLTC E) ERBB2 F) TCF4 G) VIM H) LRRK2 I) IFI16 J) CAV1

### Molecular docking studies

In the recent findings, the docking simulation was performed to find the site conformation and significant interactions responsible for complicated stability with the receptor of the binding sites. Docking experiments were performed using Sybyl X 2.1 software on natural plants products such as herbs have the ability to lower blood glucose levels and ameliorate diabetes with decreased adverse side effects. They accomplish this owing to the presence of phyto chemicals such as flavonoids (FLA, FLS, FLO, FAS, ANC, ERI, ISO, FIS, MOR NAR, DIO, CHR, CYA, API, BAI, LUT, KAE, PEL, TAN, WOG, ISH, DAI, GEN, DEL etc.), saponins (ARJ),tannins (GAL, PHL), glycosides (HES, RUT) etc. The molecules were constructed based on the natural plant products containing these chemical constituents which play vital role in reducing type 2 diabetes mellitus^3^.The traditional plant products are used in conjunction with allopathic drug to reduce the dose of the allopathic drugs and or to increase the efficacy of allopathic drugs. common and most prominent antidiabetic plants and active principles were selected from their phyto chemicals for docking studies in the present research. For docking experiments, a total of common 28 active constituents few from each of flavonoids, saponins, tannins and glycosides etc., present in plant extracts responsible for antidiabetic function, were chosen for docking studies on over expressed proteins and the structures are depicted in Fig. 8 respectively. The one protein from each over expressed genes in type 2 diabetes mellitus such as CLTC (Clathrin heavy chain), DCTN1 (Dynactin subunit 1), ERBB2 (Erb-b2 receptor tyrosine kinase 2), FLNA (Filamin A), MYH9 (Myosin heavy chain 9) and their X-RAY crystallographic structure and co-crystallized PDB code 5M5R, 2HLC, 1MFG, 3HOC and 6LKM respectively were constructed for docking. The docking on natural active constituents was conducted to classify the potential molecule and their binding affinity to proteins. A C-score greater than5are said to be active and few constituents with particular proteins obtained greater than 8 respectively. Docking experiments were carried out on a total of 28 constituents from plant products, few constituents obtained excellent binding energy (C-score) greater than 8 and few constituents obtained an optimum binding score of 4-4.9 then after a few constituents obtained less binding score of 2.0-3.0 respectively. The natural constituents HES, RUT, DIO obtained excellent binding score of more than 10 with two PDB proteins 5M5R and 4ZS4 respectively. The constituents obtained good binding score 5 to 7.99 are QUE, ANT, PEL, MOR, CYA, DEL, ISO, NAR, FLO, FIS, KAE, TAN, FLS, FLO, CHR, WOG, LUT, ERI and DIO, HES, RUT, QUE, CHR andDIO, RUT, WOG, TAN, QUE, ISO, HES,PEL and DIO, RUT, HES,MOR, CYA, KAE, DEL, ISO and MOR,KAE, TAN, WOG, ERI, ISO, ANT,GEN, LUT, FIS, FLO, QUE, DEL, NAR,DAI, CYA with PDB protein of 5M5R, 2HL3, 1MFG, 3HOC and 4ZS4 respectively. Constituents obtained optimum binding score are API,DAI, BAI, GEN, FLA, PHL and API, ISO, CYA, DEL, ISO, TAN, ERI, LUT, PEL and MOR, KAE, NAR, GEN, DAI, CYA, LUT, FIS, ERI, CHR, and NAR, ERI, QUE, LUT, WOG, API, FLO, FIS, GEN, and PEL, FLS, API, BAI, FAS, CHR, FLO, ISO, PHL with PDB code of 5M5R, 2HL3, 1MFG, 3HOC and 4ZS4 and following constituents obtained very less binding are ISO and FLO, DAI, NAR, PHL and PHL and DAI, ISO with PDB code 5M5R, 2HL3 and 1MFG respectively the values are depicted in Table 1. The constituents HES, RUT, DIO obtained good binding score with all proteins. The hydrogen bonding interactions and other bonding interactions with amino acids with protein code 5M5Rof molecule HESare depicted by 3D and 2D (Fig. 9 and Fig. 10).

**Fig. 8.**
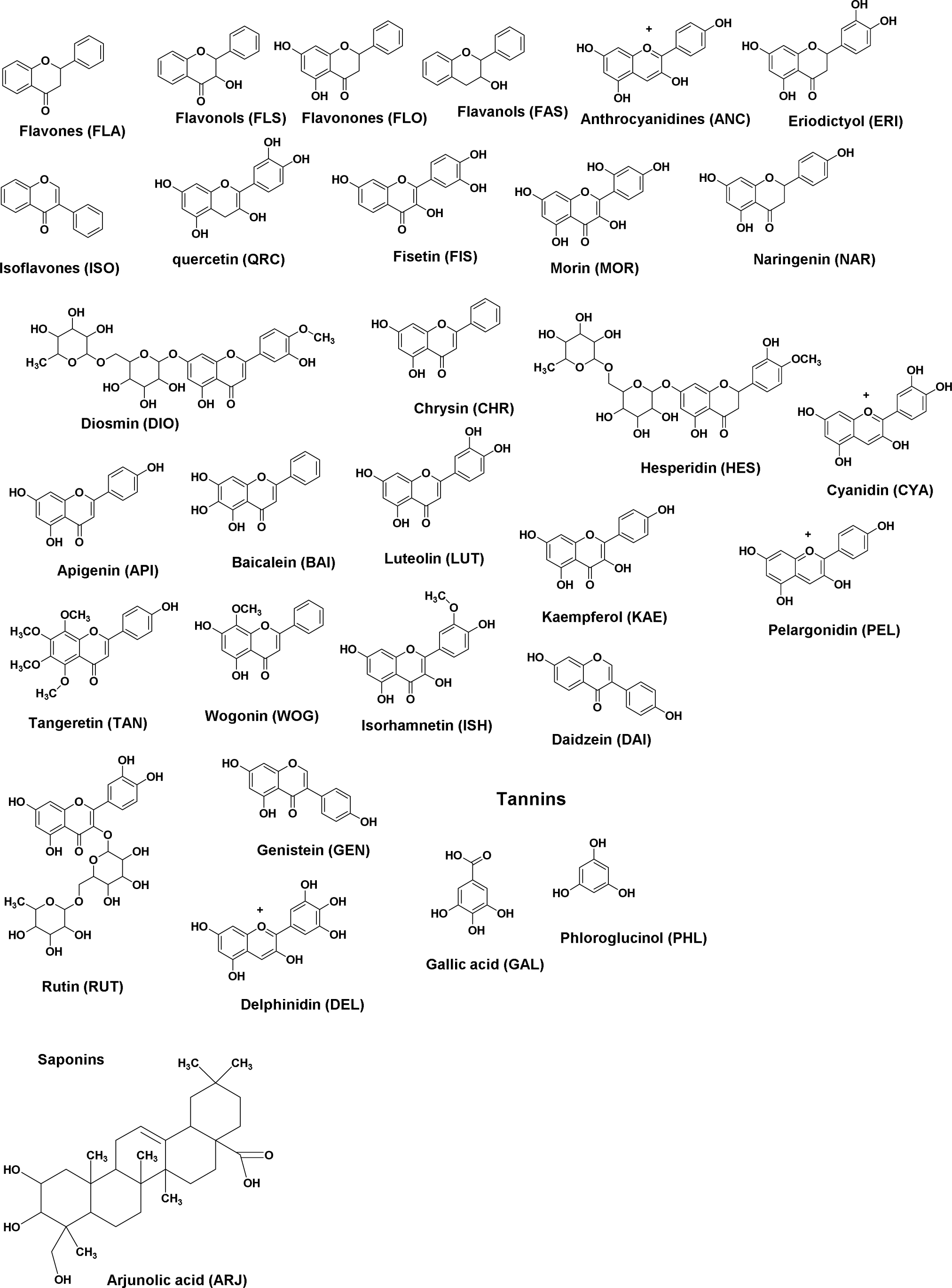
Structures of Designed Molecules

**Fig. 9.**
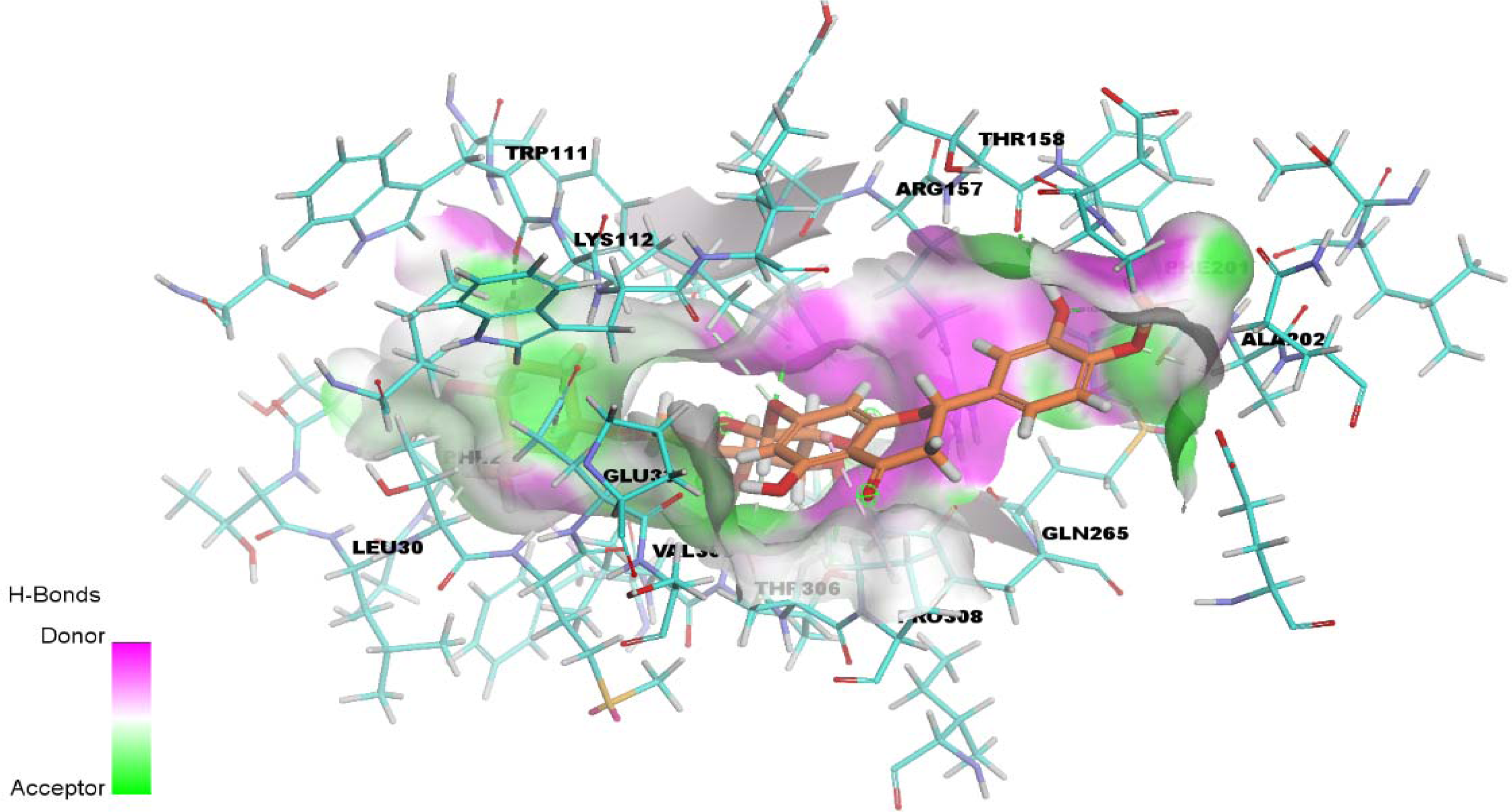
3D Binding of Molecule HES with 5M5R

**Fig.10.**
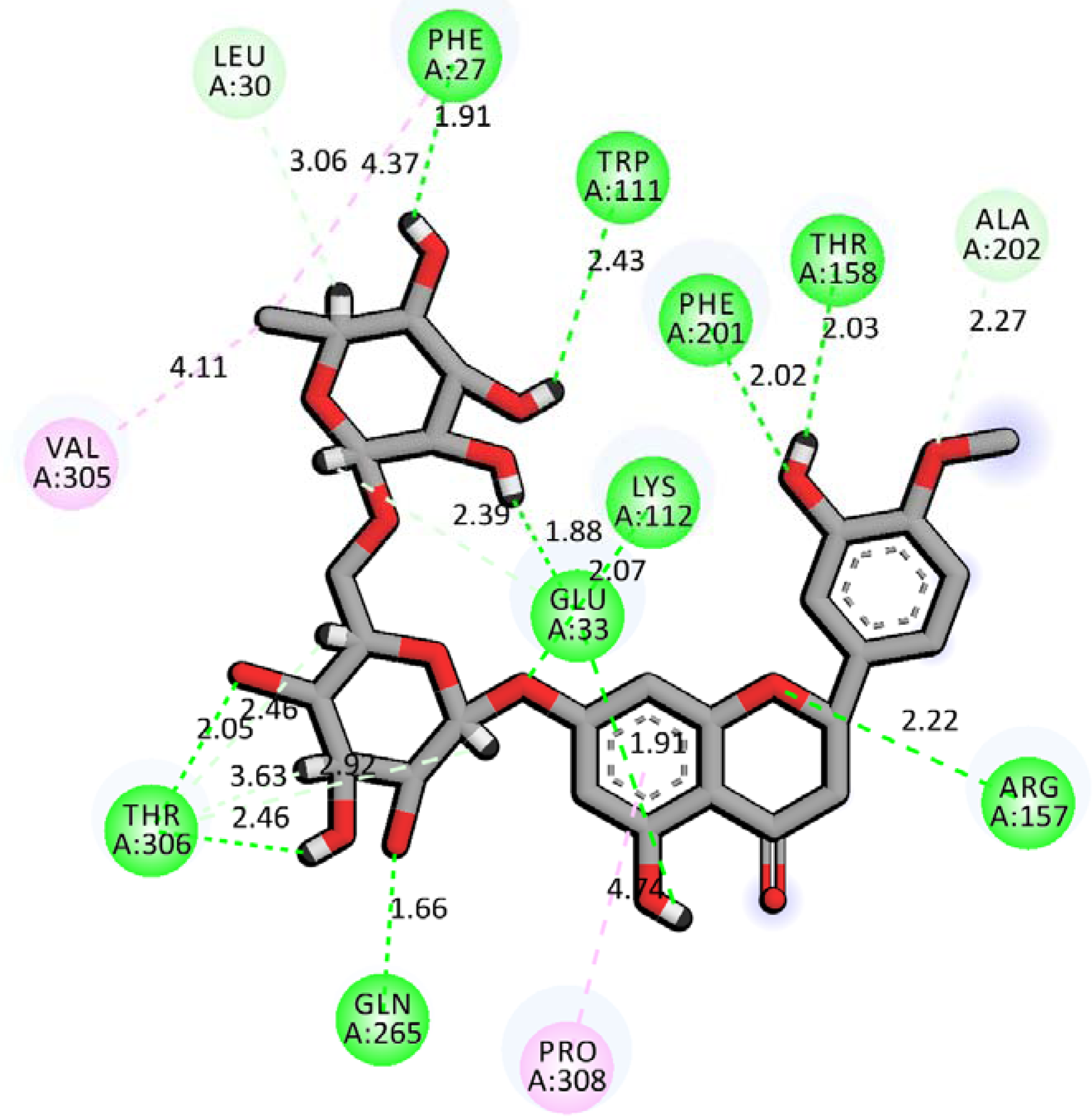
2D Binding of Molecule HES with 5M5R

## Discussion

The identification of key genes can aid our understanding of obesity associated type 2 diabetes mellitus and the discovery of novel targets for therapy of obesity associated type 2 diabetes mellitus. In the current investigation, we first integrated expression profiling by high throughput sequencing data of obesity associated type 2 diabetes mellitus, and found that 872 genes (438 up regulated and 439 down regulated genes) were differentially expressed compared with non diabetic obese samples. XIST (X inactive specific transcript) [31] and SELL (selectin L) [32] expression are significantly associated with type 2 diabetes mellitus. S100A9 and S100A8 acts in the progression of obesity associated type 2 diabetes mellitus [33]. IL1R2 [34] and SPINK5 [35] are involved in the development of asthma, but these genes might be associated with development of obesity associated type 2 diabetes mellitus.

DEGs in GO terms and pathways might be linked with advancement of obesity associated type 2 diabetes mellitus. ERBB2 [36], DACT1 [37], ARAP1 [38], MYH9 [39], INPPL1 [40], SARM1 [41], NOTCH1 [42], ROBO1 [43], MAPK8IP1 [44], ANK1 [45], SARM1 [46], SREBF2 [47], SIK1 [48], PASK (PAS domain containing serine/threonine kinase) [49], NOS2 [50], OAS3 [51], KL (klotho) [52], PECAM1 [53], S100A12 [54], S100P [55], BATF3 [56], PLEK (pleckstrin) [57], ALOX5 [58], ARG1 [59], CXCL8 [60], CXCR1 [61], PTAFR (platelet activating factor receptor) [62], PYGL (glycogen phosphorylase L) [63], TCF4 [64], CAMP (cathelicidin antimicrobial peptide) [65], RUNX2 [66], PLA2G2A [67], GCG (glucagon) [68], RARRES2 [69] and HAP1 [70] were involved in the genesis of type 2 diabetes mellitus. Recent studies have reported that ACHE (acetylcholinesterase) [71], FGFR3 [72], VLDLR (very low density lipoprotein receptor) [73], SHC1 [74], HDAC6 [75], CHRNA2 [76], CASR (calcium sensing receptor) [77], ELK1 [78], TYK2 [79], CIITA (class II major histocompatibility complex transactivator) [80], ZAP70 [81], GPT (glutamic--pyruvic transaminase) [82], CHI3L1 [83], AIF1 [84], MMP9 [85], ITGB2 [86], CFD (complement factor D) [87], C3AR1 [88], LGALS1 [89], CD14 [90], TIMP1 [91], TLR2 [92], LTF (lactotransferrin) [93], BRCA2 [94] and IGFBP3 [95] promotes the development of obesity. Sun et al [96] found that TRPM2 was significantly associated in obesity associated type 2 diabetes mellitus. Richter et al. [97], Suchkova et al [98], Qureshi et al [99], Wang et al [100], Wang et al [101], Aoki-Suzuki et al [102], Ohno et al [103], Richter et al [104], Rahman and Copeland [105], Congiu et al [106], Ji et al [107], Wollmer et al [108], Yamazaki et al [109], Bardien et al [110], Comella Bolla et al [111], Horvath et al [112], Watanabe et al [113], Kushima et al [114], Grünblatt et al [115] and Sato and Kawata [116] have shown that TAOK2, ACAP3, PLXNA3, PLXNA4, DCTN1, NTNG2, LRP4, AGRN (agrin), TAOK2, POLG (DNA polymerase gamma, catalytic subunit), KCNK2, OPRK1, ABCA2, ABCA7, LRRK2, CD200, PAK3, PADI2, EPHB1, CHAT (choline O-acetyltransferase) and SLC18A1 key for progression of neurological and neuropsychiatric disorders, but these genes might be liable for obesity associated type 2 diabetes mellitus. PLD2 [117], FLNA (filamin A) [118], SMURF1 [119], LINGO1 [120], CACNA1H [121], NLRP6 [122], NLRC3 [123], CXCR2 [124] and C5AR1 [125] are involved in hypertension, but these genes might be essential for progression of obesity associated type 2 diabetes mellitus. Studies conducted by Sauzeau et al [126], Xu et al [127], Hirota et al [128], Alharatani et al [129], Beitelshees et al [130], Zhu et al [131], Gil-Cayuela et al [132], Liu et al [133], Xie et al [134], Kroupis et al [135], López-Mejías et al [136], Gremmel et al [137], Yamada and Guo [138], Petri et al [139], DeFilippis et al [140], Rocca et al [141] and Tur et al [142] indicated that of VAV2, RASAL1, LIF (LIF interleukin 6 family cytokine), CTNND1, CACNA1C, MAP3K10, NRBP2, TRPM4, LILRB2, FCGR2A, PIK3CG, SELPLG (selectin P ligand), PRDX4, FPR2, PLG (plasminogen), SELENOM (selenoprotein M) and NCAM1were associated with cardiovascular diseases, but these genes might be responsible for progression of obesity associated type 2 diabetes mellitus. Previous studies indicated that ITGB4 [143], SEMA3D [144], FCAR (Fc fragment of IgA receptor) [145], KIT (KIT proto-oncogene, receptor tyrosine kinase) [146], PGLYRP1 [147], IL17RB [148], BIRC5 [149] and PTGS1 [150] plays an important role in asthma, but these genes might be linked with progression of obesity associated type 2 diabetes mellitus. GRK2 [151], ADCY3 [152], FASN (fatty acid synthase) [153], DGKD (diacylglycerol kinase delta) [154], DGKQ (diacylglycerol kinase theta) [154], IP6K1 [155], ANXA1 [156], SUCNR1 [157], PRNP (prion protein) [158], CXCR4 [159], CAV1 [160], LCN2 [161], AQP9 [162], NMU (neuromedin U) [163], NPY1R [164], FFAR2 [165], OSM (oncostatin M) [166] and TREM1 [167] were reported in obesity associated type 2 diabetes mellitus. UNC13B [168], PFKFB3 [169], FCN1 [170] and SLC11A1 [171] were involved in progression of type 1 diabetes, but these genes might be liable for progression of obesity associated type 2 diabetes mellitus.

Protein-protein interaction (PPI) network and its module can be regarded as key to the understanding of progression of obesity associated type 2 diabetes mellitus and might also lead to novel therapeutic way. VIM (vimentin) has been implicated as a principal mediator of obesity associated type 2 diabetes mellitus [172]. IFI16 is crucial for advancement of obesity [173]. CLTC (clathrin heavy chain), TNS2, PLCG1 and NIFK (nucleolar protein interacting with the FHA domain of MKI67) might be the novel biomarkers for obesity associated type 2 diabetes mellitus.

To validate the accuracy of the target genes, miRNAs and TFs identified by target gene - miRNA regulatory network and target gene - TF regulatory network analysis. Yan et al [174], Wang et al [175], Yan et al [176] and Guo et al [177] showed that hsa-mir-4329, hsa-mir-3685, hsa-mir-6124, hsa-mir-1297 and SMARCA4 expression can be associated with cardiovascular diseases progression, but these genes might be essential for advancement of obesity associated type 2 diabetes mellitus. hsa-mir-1299 [178], hsa-mir-4779 [179] and hsa-mir-4459 [180] play a crucial role in type 2 diabetes mellitus. Expression of TCF7 was liable for progression of type 1 diabetes [181], but this gene might be responsible for development of obesity associated type 2 diabetes mellitus. MYB transrption factor was associated with progression of asthma [182], but these genes might be linked with progression obesity associated type 2 diabetes mellitus. E2F4 [183] and transcription factor CLOCK [184] were linked with development of obesity associated type 2 diabetes mellitus. CUX1 [185], NANOG [186], GATA4 [187] and HIF1A [188] were essential for progression of obesity. MYO18A, SEC16A, CCNB1, MAD2L1, hsa-mir-4315, hsa-mir-6134, hsa-mir-9500, KIFC3, FBL (fibrillarin), TUBA1A and GFI1B might be the novel biomarkers for obesity associated type 2 diabetes mellitus.

## Conclusion

Based on integrated bioinformatics analysis including GO and REACTOME pathway enrichment, PPI network, module analysis, target gene - miRNA regulatory network and target gene - TF regulatory network, and hub and target gene identification, the present study showed that DEGs that might have the potential to serve as reliable molecular biomarkers for the diagnosis, prognosis and therapeutic target of obesity associated type 2 diabetes mellitus. Moreover, MYH9, FLNA, DCTN1, CLTC, ERBB2, TCF4, VIM, LRRK2, IFI16 and CAV1 were validated in obesity associated type 2 diabetes mellitus. Further studies are merited to explore the biological functions of these genes and the underlying mechanisms associated in the pathogenesis of obesity associated type 2 diabetes mellitus.

## Acknowledgement

I thank Stephane Le Crom, IBENS, Genomic Core Facility, Paris, France, very much, the author who deposited their expression profiling by high throughput sequencing dataset, GSE113079, into the public GEO database.

## Ethical approval

This article does not contain any studies with human participants or animals performed by any of the authors.

## Informed consent

No informed consent because this study does not contain human or animals participants.

## Availability of data and materials

The datasets supporting the conclusions of this article are available in the GEO (Gene Expression Omnibus) (https://www.ncbi.nlm.nih.gov/geo/) repository. [(GSE132831) (https://www.ncbi.nlm.nih.gov/geo/query/acc.cgi?acc=GSE132831)]

## Consent for publication

Not applicable.

## Author Contributions

**A**

**Author Contributions**

B. V - Writing original draft, and review and editing

A. T - Formal analysis and validation

C. V - Software and investigation

I. K - Supervision and resources

